# Self-organized yolk sac-like organoids allow for scalable generation of multipotent hematopoietic progenitor cells from human induced pluripotent stem cells

**DOI:** 10.1101/2021.04.25.441298

**Authors:** Naritaka Tamaoki, Stefan Siebert, Takuya Maeda, Ngoc-Han Ha, Meghan L. Good, Yin Huang, Suman Kumar Vodnala, Juan J. Haro-Mora, Naoya Uchida, John F. Tisdale, Colin L. Sweeney, Uimook Choi, Julie Brault, Sherry Koontz, Harry L. Malech, Yasuhiro Yamazaki, Risa Isonaka, David S. Goldstein, Masaki Kimura, Takanori Takebe, Jizhong Zou, David F. Stroncek, Pamela G. Robey, Michael J. Kruhlak, Nicholas P. Restifo, Raul Vizcardo

## Abstract

The human definitive yolk sac is an important organ supporting the early developing embryo through nutrient supply and by facilitating the establishment of the embryonic circulatory system. However, the molecular and cellular biology of the human yolk sac remains largely obscure due to the lack of suitable *in vitro* models. Here, we show that human induced pluripotent stem cells (hiPSCs) co-cultured with various types of stromal cells as spheroids self-organize into yolk sac-like organoids without the addition of exogenous factors. Yolk sac-like organoids recapitulated a yolk sac specific cellular complement and structures as well as the functional ability to generate definitive hematopoietic progenitor cells (HPCs). Furthermore, sequential hemato-vascular ontogenesis could be observed during organoid formation. Notably, our organoid system can be performed in a scalable, autologous, and xeno-free condition, thereby providing an important model of human definitive yolk sac development and allows for efficient bulk generation of hiPSC-derived HPCs.

## INTRODUCTION

The mammalian yolk sac supports the developing embryo by nutrient absorption and formation of the embryonic circulatory system which can deliver oxygen and nutrients to the embryo during early embryogenesis (Cindrova-Davies et al., 2017; Ross and Boroviak, 2020; Zohn and Sarkar, 2010). The yolk sac is a double-layered organ composed of an endodermal and a mesodermal layer. The yolk sac endoderm serves the function of nutrient uptake, while the major function of yolk sac mesoderm is to establish the vascular plexus and hematopoiesis (Palis and Yoder, 2001; Sheng and Foley, 2012; Zohn and Sarkar, 2010). Human yolk sac development occurs in two phases including the formation of the transient primary yolk sac followed by the formation of the functional definitive yolk sac (Ross and Boroviak, 2020). It has been postulated that the human definitive yolk sac supports the embryo in ways similar to what has been observed in other mammals, but the cellular and molecular biology remains largely obscure due to the ethical and technical challenges associated with the access to early human embryos. Since basic structures and roles of the yolk sac are highly conserved across species (Cindrova-Davies et al., 2017; Sheng and Foley, 2012; Zohn and Sarkar, 2010), most of our understanding about the yolk sac is based mainly on studies in mice due to the ease of genetic manipulation. However, there are several significant differences between murine and human yolk sacs including anatomical features and the complexity of the developmental process in humans (Ross and Boroviak, 2020). Consequently, direct extrapolation from murine study data to human biology may be inadequate.

The generation of human induced pluripotent stem cells (hiPSCs) has unleashed an enormous potential for the investigation of human developmental processes. hiPSC differentiation *in vitro* can recapitulate the dynamics and complexity of embryonic development without raising ethical concerns (Sharma et al., 2020). Indeed, many research groups have demonstrated that hiPSC differentiation to multipotent hematopoietic progenitor cells (HPCs) follows the hemato-vascular ontogenesis observed in the mouse yolk sac (Ditadi et al., 2017; Slukvin, 2013). These studies provide a foundation for establishing experimental models that mimic human embryonic vasculogenesis and hematopoiesis observed in the yolk sac mesoderm. However, the precise functions of human yolk sac endoderm during embryogenesis are still poorly understood, because an *in vitro* model using hiPSC-derived yolk sac endoderm cells to study their cellular and molecular biology has not been established. Furthermore, the hemato-vascular development in the yolk sac is orchestrated by interactions between yolk sac mesoderm and endoderm (Drevon and Jaffredo, 2014; Giles et al., 2005; Goldie et al., 2008; M.A. Dyer, 2001), therefore modeling both yolk sac compartments is desirable to investigate the signaling molecules involved in this process. Recently, the combination of hiPSC and three-dimensional (3D) culture technologies enabled the generation of hiPSC-derived organoids representing the remarkable complexity of organ-specific cell types and structural features, that also function similarly to their *in vivo* counterparts (McCauley and Wells, 2017). Although many types of organoids, such as brain, liver, and kidney, have been generated successfully from hiPSCs (Camp et al., 2015; Low et al., 2019; Takebe et al., 2013), definitive yolk sac organogenesis from hiPSCs has not been reported. The capacity to engineer yolk sac organoids from hiPSCs would enable *in vitro* monitoring of the interactions between mesoderm and endoderm layers during the hemato-vascular development.

In this study, we developed a simple and scalable system to generate yolk sac-like organoids from hiPSCs without the addition of exogenous factors. Our yolk sac-like organoids recapitulated not only human definitive yolk sac-specific cellular components and structures but also the functional ability to generate HPCs. Similar to hiPSC differentiation *in vitro*, a sequential hemato-vascular developmental process was observed during organoid development. Single-cell RNA sequence (scRNA-seq) analysis revealed transcriptional signatures and marker gene expression consistent with cellular functions that have been reported for mouse yolk sac endoderm as well as genes that regulate the hemato-vascular specification from mesoderm progenitors. Additionally, we demonstrated that organoid induced HPCs possess definitive hematopoiesis to generate erythroid, myeloid and lymphoid cells. Importantly, our organoid system is scalable and can be performed in an autologous and xeno-free condition. In summary, our yolk sac-like organoid system offers the opportunity to study human yolk sac development and represents a new avenue for clinical application of hiPSC-based blood cell therapy allowing for reproducible and cost-effective bulk production of HPCs from patient-specific hiPSCs.

## RESULT

### Hematopoietic spheroid system produces HPCs from hiPSCs in the absence of exogenous cytokines/growth factors

Previous studies have shown that spontaneous differentiation of embryoid bodies (EBs) grown in 3D conditions can represent some yolk sac features and induce hematopoiesis (Poon et al., 2006; Zambidis et al., 2005). However, this approach led to EBs differentiating randomly into all three germ layers, resulting in high variability and low efficacy of hematopoietic specification that is a unique and important function of the human definitive yolk sac. Thus, we sought to engineer a consistent culture system to facilitate the formation of yolk sac-like structures and the induction of hematopoiesis from hiPSCs. Since mouse bone marrow-derived OP9 stromal cells can present a hematopoietic niche-like microenvironment to induce hematopoiesis of human pluripotent stem cells (hPSCs) without the addition of exogenous growth factors (Nakano et al., 1994; Vodyanik et al., 2006), we speculated that this unique feature can be applied to engineer hiPSC-derived hematopoietic organoids. As 3D culture techniques can provide improved physiological growth conditions to promote organoid formation compared to two-dimensional (2D) conditions, we designed a system to co-culture OP9 cells and EBs as spheroids (Figure 1A). EBs mixed with OP9 cells in ultra-low adherent microwells (Aggrewell plates) developed into uniform-sized spheroids (Figure 1B). Small EBs (100-150 cells/EB) were used to ensure more cell-to-cell contact between hiPSCs and OP9 cells, improving intercellular signaling and cell-extracellular matrix interactions. Because EBs sink faster than OP9 cells in microwells, EBs and OP9 cells did not mix homogeneously in the forming spheroids, but this approach produced a mechanical resistant spheroid with firm interactions between EBs and OP9 cells in a 3D manner (Figures 1A and 1B).

**Figure 1.**
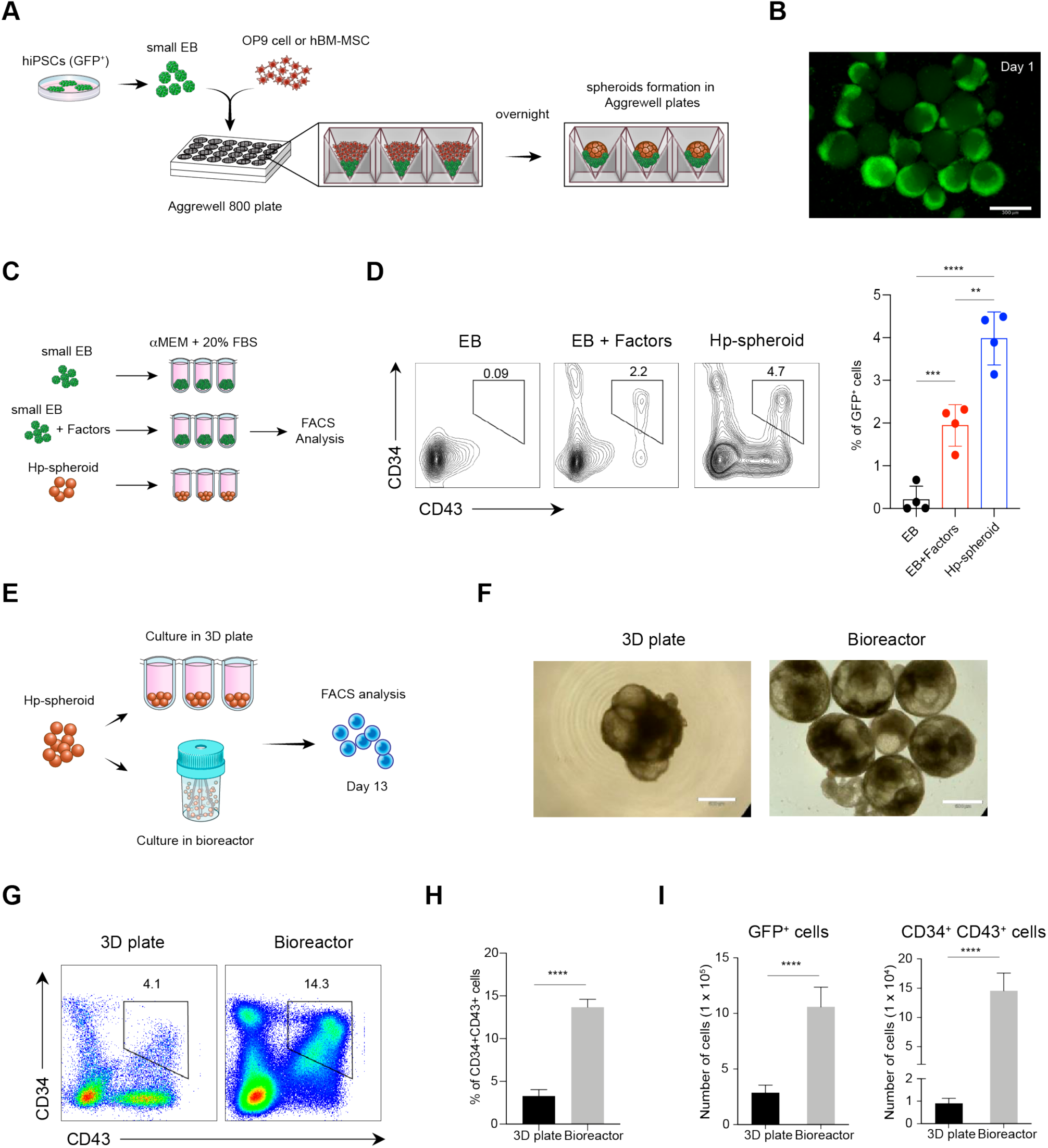
Hp-spheroid system can induce hematopoiesis from hiPSCs without addition of exogenous cytokines/growth factors. (A) Schematic of co-culturing hiPSCs and stromal cells as spheroids in 3D conditions. (B) Representative GFP fluorescence microscopy image of Hp-spheroids on day 1. Scale bar = 500μm. (C) Experimental design to assess the induction of HPCs using the Hp-spheroid system (co-culturing with hBM-MSCs) in 3D culture plates compared to the EB mediated differentiation. (D) Representative flow cytometry analysis of CD34^+^ CD43^+^ cells in GFP^+^ cells isolated from EBs, EBs with exogenous factors, and Hp-spheroids (left). Percentage of CD34^+^CD43^+^ cells in GFP^+^ cells measured on day 13 by flow cytometry analysis (right). Values represent mean ± SD (n = 4, **p<0.005, ***p<0.001, ****p < 0.0001). (E) Schematic representation of the comparative experiment of the Hp-spheroid system using stirred suspension bioreactors and 3D culture plates. (F) Light microscopy image of Hp-spheroids cultured in 3D culture plates (left) and in bioreactors (right) for 13 days. Scale bar = 500μm. (G) Representative flow cytometry analysis of CD34^+^ and CD43^+^ cells in Hp-spheroids on day 13. GFP^+^ cells are gated for analysis. (H) Percentage of CD34^+^CD43^+^ cells in GFP^+^ cells isolated from day 13 Hp-spheroids. Values represent mean ± SD (n = 3, ****p < 0.0001). (I) Quantifications of GFP^+^ cells (left) and CD34^+^CD43^+^ cells (right) in Hp-spheroids on day 13. Graph shows the results obtained from 1 x 10^6^ hiPSCs. Values represent mean ± SD (n = 3, ****p < 0.0001).

We first attempted to identify CD34^+^CD43^+^ HPCs to determine whether our OP9/EB spheroids system can induce hematopoiesis. To track hiPSC-derived cells during co-culture, we used a stable EGFP (enhanced green fluorescent protein)-expressing hiPSC line (NCRM5-AAVS1-CAG-EGFP, hereinafter referred to as NCRM5-EGFP) (Luo et al., 2014). The OP9/EB co-cultured spheroids were transferred from Aggrewell plates to ultra-low attachment 3D culture plates and were cultured in αMEM medium containing 20% FBS for 13 days (**Figures 1A and S1A**). Interestingly, OP9/EB spheroids developed into cystic structures (**Figure S1B**) containing GFP^+^CD34^+^CD43^+^ cells on day 13, similar to the hematopoietic zones observed in the classical OP9 co-culture system in 2D (Timmermans et al., 2009) (**Figure S1C**). Hereinafter this co-culture spheroid that can induce hematopoiesis is referred to as hematopoietic spheroid (Hp-spheroid). To clarify whether providing a hematopoietic niche-like microenvironment in 3D is a key element for our Hp-spheroid system, we decided to use human bone marrow-derived mesenchymal stem/stroma cells (hBM-MSCs), since these cells contribute to the maintenance of the hematopoietic niche in the bone marrow (Kfoury and Scadden, 2015). Additionally, at the time of this study there was no published work indicating whether hBM-MSCs can induce hematopoiesis of hiPSC similar to OP9 cells. Based on our experience of using the classical OP9 co-culture system (Good et al., 2019), we reasoned that the gradual loss of stem cell properties in hBM-MSCs during expansion may be detrimental. Therefore, we limited the expansion of hBM-MSCs to 4-5 passages prior to Hp-spheroid formation (Sabatino et al., 2012). Similar to when OP9 cells were used, NCRM5-EGFP cells co-cultured with hBM-MSCs in the Hp-spheroid system generated GFP^+^CD34^+^CD43^+^ cells by day 13. Strikingly, this process occurred in the absence of exogenous factors which are required for HPC differentiation from EBs (**Figures 1C and 1D**). It is also important to note that hBM-MSCs co-cultured in a 2D monolayer did not generate CD34^+^CD43^+^ cells from hiPSCs (**Figure S1D**), indicating that our Hp-spheroid system was not simply a conversion of the classical OP9 co-culture system to a 3D environment. Significantly, using hBM-MSCs to form Hp-spheroids enabled us to generate hiPSC-derived HPCs consistently without exogenous factors or murine stromal cells.

Next, we sought to build a scalable Hp-spheroid system. Since Hp-spheroids showed a strong resilience to mechanical stress, we were able to culture them in stirred suspension bioreactors after optimizing the rotation speed. Hp-spheroids grown in the bioreactors were homogeneous spheres with internalized cystic structures (**Figures 1E and 1F**). Notably, based on the size of the Hp-spheroids and the number of GFP^+^ cells, the use of bioreactors enhanced the growth of hiPSC as well as the fidelity of HPC induction (**Figures 1F-1I**). A kinetic analysis of HPC markers by flow cytometry showed that HPCs started to appear by day 9 and peaked on day 13-14 of culturing (**Figure S2A**). Since both primitive and definitive types of hematopoiesis can be found in the human definitive yolk sac, we tried to clarify the hematopoietic potential of HPCs generated by this system. The hematopoietic colony-forming unit (CFU) assay demonstrated that Hp-spheroid-derived HPCs cells generated both erythroid and myeloid lineage cells (**Figure S2B**). Furthermore, the T cell differentiation potential was evaluated by co-culturing on OP9/DLL1 in the presence of cytokines (SCF, Flt3-L, and IL7) (Vizcardo et al., 2013). CD5^+^CD7^+^ T cell progenitors were detected by day 17, and CD4^+^CD8^+^ double positive T cells were detected by day 27 of T cell differentiation (**Figure S2C**). Transplantation of NCRM5-EGFP-derived CD34^+^ cells into immunocompromised mice failed to gain engraftment, suggesting that HPCs generated in Hp-spheroids were not as mature as adult hematopoietic stem cells (HSCs) (data not shown). Altogether, these data demonstrated that the use of bioreactors with our Hp-spheroid system could provide a scalable approach to generate definitive HPCs from hiPSCs without using a complicated cocktail of cytokines and growth factors.

### The Hp-spheroids form yolk sac-like structures and recapitulate the dynamics of hemato-vascular developmental processes

Given that day 13 Hp-spheroids were able to functionally generate HPCs similar to the human definitive yolk sac, we explored if there were also structural similarities. Hematoxylin and eosin staining suggested that day 13 Hp-spheroids are composed of an outer layer of cells encompassing multiple inner cystic structures (Figure 2A). Notably, whole-mount immunostaining and confocal imaging revealed that only the outer cell layer expressed AFP (alpha fetoprotein) whereas the PDGFRβ^+^ (platelet-derived growth factor receptor β) cells were located inside of the Hp-spheroids, indicating that the endoderm (AFP) and mesoderm (PDGFRβ) regions are distinctly organized on day 13 (Figure 2B). To visualize the distribution of hematopoietic cells on day 13, we performed immunostaining for CD34 and CD43. Similar to the anatomical feature of hematopoietic cells in the blood island of the human definitive yolk sac, CD43^+^ hematopoietic cells were detected as cell-clusters within the mesodermal region (Ferkowicz and Yoder, 2005) with a subset of CD34^+^CD43^+^ putative HPCs (Figures 2C and 2D). These unique structural features of day 13 Hp-spheroids supported that they were morphologically similar to the human definitive yolk sac (Ross and Boroviak, 2020). Taken together, day 13 Hp-spheroids possessed functional and morphological features of the human definitive yolk sac, indicating that Hp-spheroids developed into yolk sac-like organoids.

**Figure 2.**
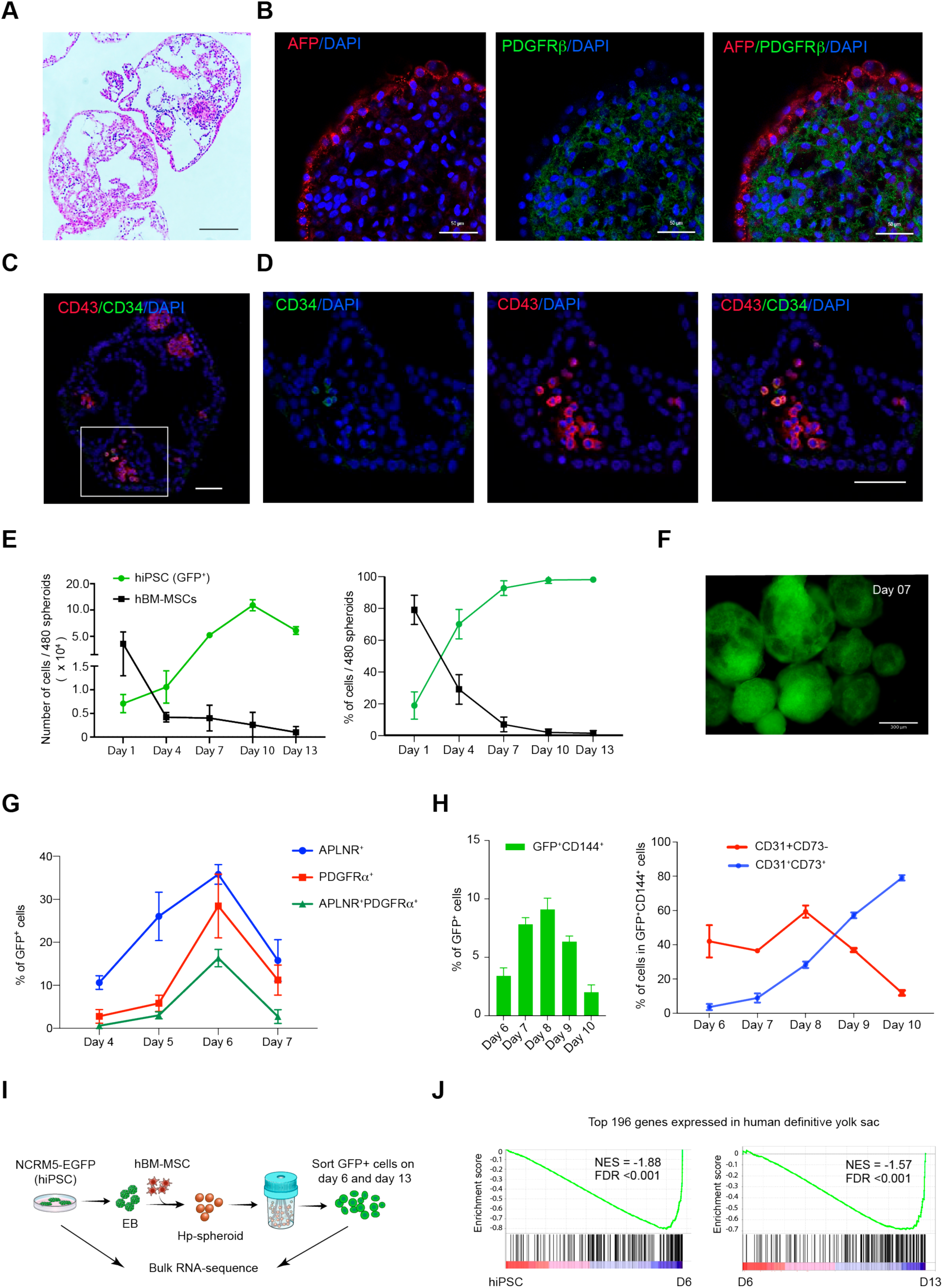
The Hp-spheroid system induces hiPSCs to self-organize into yolk sac-like organoids. (A) Hematoxylin and eosin staining of day 13 Hp-spheroids. Scale bar = 200μm. (B) Whole-mount immunostaining analysis of AFP and PDGFRβ in a Hp-spheroid on day 13. Scale bar = 50μm. (C) Immunostaining of a Hp-spheroid on day 13 for CD34 and CD43. Scale bar = 50μm. (D) Higher magnification of the boxed area in C. Scale bar = 50μm. (E) Quantifications of the number (left) and the percentage (right) of GFP^+^ cells (hiPSCs) and GFP^−^ cells (hBM-MSCs) in Hp-spheroids for 13 days. Values represent mean ± SD (n = 3). Representative flow cytometry analysis is shown in **Figure S3A**. (F) Fluorescence microscopy image of Hp-spheroids on day 7. Scale bar = 500μm. (G) Kinetics analysis of APLNR and PDGFRα expression on GFP^+^ cells in Hp-spheroids from day 4 to day 7 by flow cytometry analysis. Representative flow cytometry analysis is shown in **Figure S3C**. (H) Quantification of flow cytometry analysis of CD144 expression on GFP^+^ cells in Hp-spheroids from day 6 to day 10 of organoids formation (left), and the frequencies of CD31^+^CD73^−^ cells and CD31^+^CD73^+^ cells in GFP^+^CD144^+^ population (right). Values represent mean ± SD (n = 3). Representative flow cytometry analysis is shown in **Figure S3D**. (I) Schematic outline of bulk RNA-seq analysis. (J) Representative GSEA plots of the top 196 genes which are expressed in the human yolk sac. NES; normalized enrichment score; FDR, false-discovery rate.

During embryogenesis, vasculogenesis proceeds in parallel with hematopoiesis in the yolk sac mesoderm. We sought to understand whether the dynamics of hemato-vascular ontogenesis is also recapitulated during the organoid development. To this end, we traced NCRM5-EGFP cells (GFP^+^ cells) by kinetic flow cytometry analysis. Although we used equivalent numbers of both NCRM5-EGFP cells and hBM-MSCs (GFP^−^ cells) to establish Hp-spheroids, we found almost all cells in day 13 organoids to be GFP^+^ cells (**Figure S3A**). The same result was obtained when OP9 cells were used (data not shown). To determine how long hBM-MSCs contributed to the organoid development, we tracked both GFP^+^ and GFP^−^ cell populations through day 13 of co-culture. Remarkably, kinetic flow cytometry analysis revealed that hBM-MSCs decreased dramatically through day 4 of co-culture, whereas EGFP-labeled hiPSCs continued to grow to eventually form the entirety of the organoid (**Figures 2E, 2F and Figure S3A**). GFP^+^ cells increased the expression of stromal cell markers, such as CD44 and CD73, after day 4 of co-culture, suggesting that hiPSCs differentiated into stromal type cells to compensate for the loss of hBM-MSCs (**Figure S3B**). All these observations also proposed that our organoid system induces hematopoiesis in a very different way compared to the classical OP9 co-culture system in 2D.

Embryonic hemato-vascular development begins from the induction of multipotent mesodermal progenitor cells with hematoendothelial potential (Slukvin, 2013; Uenishi et al., 2014). These specific mesodermal progenitors can be detected by co-expression of APNLR^+^ PDGFRα^+^ during hPSC differentiation (Choi et al., 2012; Uenishi et al., 2014; Yu et al., 2012). Kinetic flow cytometry analysis showed that GFP^+^APNLR^+^ PDGFRα^+^ cells were detected from day 5 of the organoid development, then peaked on day 6 and declined towards day 7 (**Figure 2G and Figure S3C**). The next developmental stage is the commitment of mesodermal progenitors to become hemogenic endothelium (HE) cells, which are common precursors of endothelial and blood cells, subsequently followed by endothelial-to-hematopoietic transition (EHT) to branch into hematopoietic and endothelial cells (Choi et al., 2012; Ditadi et al., 2015; Slukvin, 2013). To observe a sequential process in this stage, we analyzed various endothelial cell markers, CD144 (VE-cadherin), CD31, and CD73 by kinetic flow cytometry. When mesoderm progenitors become committed to the bipotential HE cell stage, they start to express pan-endothelial markers CD144 and CD31, and then differentiate into mature endothelial cells (CD144^+^CD31^+^CD73^+^) and hematopoietic cells (CD144^−^CD31^−^CD73^−^) after the EHT process (Choi et al., 2012; Ditadi et al., 2015). Similarly, GFP^+^ cells increased CD144 expression from day 6 and peaked on day 8 (**Figure 2H and Figure S3D**), followed by a decline towards day 10. Further analysis of these CD144^+^ cells revealed that most CD144^+^ cells were CD31^+^CD73^−^ until day 8, and then switched to CD31^+^CD73^+^ in conjunction with the decrease of CD144 expression. Our observations indicated that the peak of HE cell formation was on day 8, and then most HE cells completed the transition process by day 10 (**Figure 2H**). This was consistent with analysis of HPC marker kinetics which showed a dramatic increase of HPCs from day 9 to day 12 (**Figure S2A**).

To ascertain the global gene expression signatures of developing organoids, we performed bulk RNA sequencing of NCRM5-EGFP cells (hiPSCs) and GFP^+^ cells isolated from organoids on day 6 and on day 13, which represent the intermediate and final stages of hemato-vascular development respectively (**Figure 2I**). Gene set enrichment analysis (GSEA) using a human definitive yolk sac gene set comprising the top 196 differentially expressed genes (Cindrova-Davies et al., 2017) indicated that the gene expression signature of organoids became more similar to that of human definitive yolk sac as development progressed (**Figure 2J; Table S1**). Collectively, our data indicate that our Hp-spheroid system stimulated hiPSCs to initiate self-assembly of yolk sac-like organoids and generate HPCs in a manner representative of hemato-vascular development in the human definitive yolk sac.

### Single-cell RNA sequencing analysis reveals a day 6 organoid cell state composition similar to the early human definitive yolk sac

Since the yolk sac mesoderm dramatically changes its cell composition during hemato-vascular development, we sought to describe the cell complement of day 6 and day 13 organoids by single-cell transcriptome analysis. We harvested GFP+ cells from day 6 and day 13 organoids and performed scRNA-seq analysis (Figure 3A). This yielded 4,895 day 6 cells, and 4,401 day 13 cells with median counts of 5,292 genes and 30,096 transcripts (Unique Molecular Identifiers) per cell after filtering (Figure S4A). We clustered cells from both days separately and in combination, identified biomarkers and annotated the obtained clusters by using expression of previously described marker genes (**Figures 3B, S4B, S4C, S4E and S5A-C; Tables S2 and S3**) (Bernardo et al., 2011; Choi et al., 2012; Cindrova-Davies et al., 2017; Krendl et al., 2017; Lilly et al., 2016; Marchand et al., 2011; Uenishi et al., 2018; Vodyanik et al., 2010).

**Figure 3.**
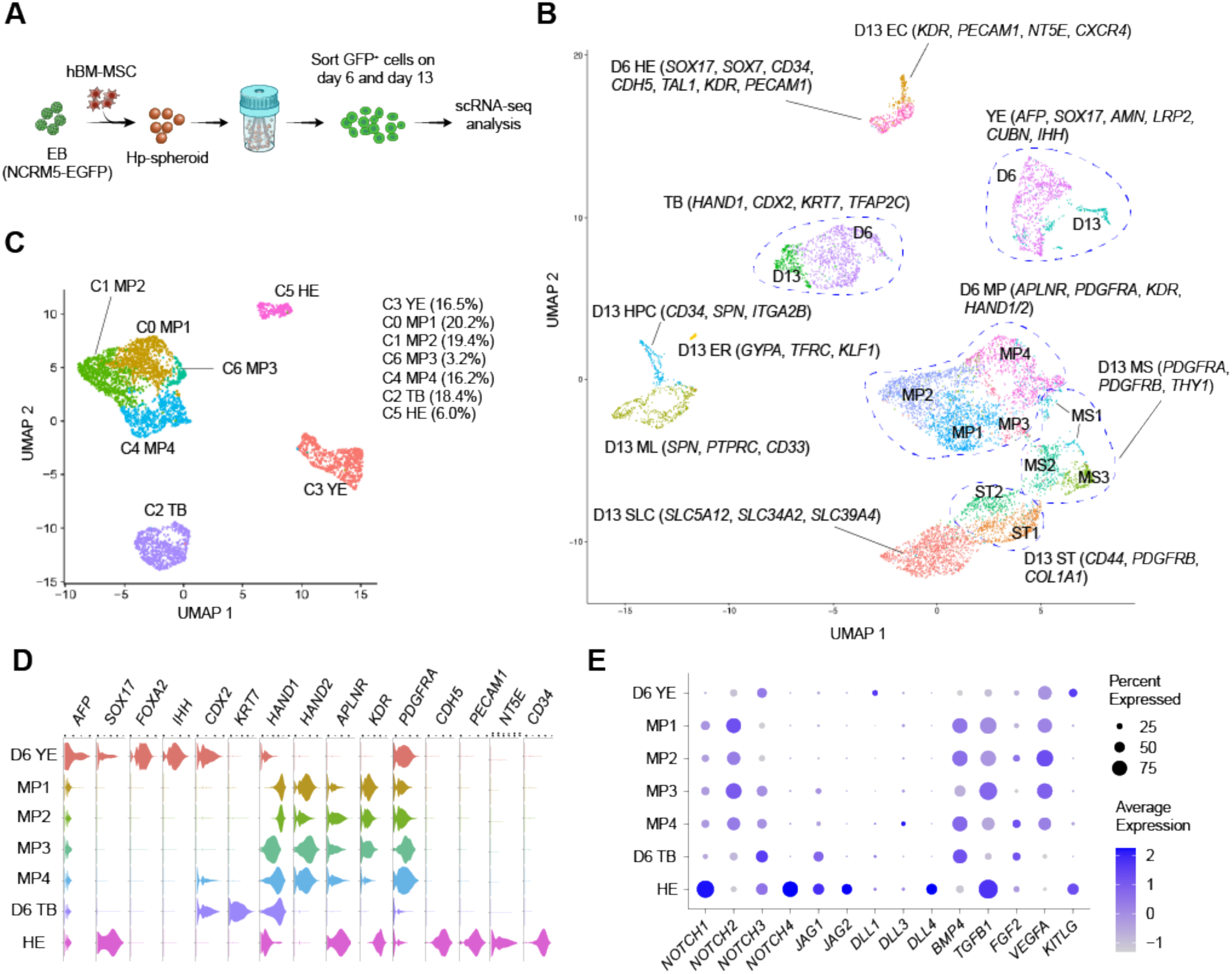
Single-cell RNAseq analysis reveals similarities between the day 6 organoid cell complement and the early human definitive yolk sac. (A) Schematic outline of scRNA-seq analysis of GFP+ cells isolated from day 6 and day 13 organoids. (B) UMAP plot for scRNA-seq analysis of 9,296 GFP^+^ cells isolated from both day 6 and day 13 organoids. Cluster annotations that were obtained when clustering day 6 and day 13 cells separately are visualized. (C) UMAP plot of 4895 GFP^+^ cells isolated from day 6 organoids. (D) Violin plots showing the expression of representative lineage marker genes for day 6 clusters. (E) Dot plot representing the gene expression of Notch signaling and key morphogens required for the hemato-vascular development in the yolk sac within clusters. Circles are coded by color (average gene expression level) and size (proportion of cells in clusters expressing a gene).

The human definitive yolk sac originates from mesodermal and endodermal cells. Consistent with this, we detected mesodermal and endodermal gene expression signatures in day 6 organoids (**Figures 3C and 3D**). For day 6 cells, 4 distinct populations were evident in the Uniform Manifold Approximation and Projection (UMAP) (**Figure 3C**). Cells within a single cluster showed elevated expression of both endodermal markers (*AFP*, *SOX17*, and *FOXA2*) as well as genes known to be expressed in the human definitive yolk sac (*AMN*, *LRP2*, *CUBN*, *TTR*, and *IHH*) (**Figures 3D and S4C**) (Cindrova-Davies et al., 2017) and this cluster was annotated as yolk sac endoderm (cluster YE). The main functions of the human yolk sac endoderm are absorption of nutrients from the exocoelomic cavity and shuttling of these nutrients to the developing embryo (Burton et al., 2001; Pereda and Motta, 1999; Zohn and Sarkar, 2010). Nutrient uptake is performed by endocytosis, therefore endocytic receptor complex associated genes (*CUBN*, *LRP2*, and *AMN*) are highly expressed in human yolk sac endoderm (Burke et al., 2013; Ross and Boroviak, 2020) and, consistent with this, in cells of cluster YE (**Figure S4C**). Moreover, genes required for transport of nutrients, such as the vitamin A transport gene *TTR*, the solute carrier (SLC) (*SLC39A5* and *SLC39A14*), and various apolipoprotein genes (*APOA1*, *APOA2*, *APOA4*, *APOB*, *APOE* and *APOM*) were highly expressed in YE cells (**Figure S4D**) (Cindrova-Davies et al., 2017; Ross and Boroviak, 2020). Mouse yolk sac endoderm secretes soluble factors, such as IHH (Indian Hedgehog) and VEGF (vascular endothelial growth factor), to regulate the commitment of mesoderm progenitors to HE cells (Goldie et al., 2008; M.A. Dyer, 2001). Similarly, high expression of *IHH* and *VEGFA* was detected in cells of cluster YE (**Figures 3D, 3F, and S4C**). These expression data demonstrated that endodermal cells generated in the organoids showed expression of several genes with functions in the mouse and human yolk sac endoderm.

Our kinetic flow cytometry analysis revealed that mesoderm progenitors peaked on day 6 (**Figure 2G**). Consistent with this finding, the most dominant population in day 6 organoids were mesoderm progenitors (*KDR*, *APLNR*, and *PDGFRA*; clusters MP) (**Figures 3D and S5A**). In accordance with previous reports, MP cells also expressed lateral plate/extraembryonic mesoderm genes (*HAND1/2* and *FOXF1*) (**Figures 3D and S5A**) (Vodyanik et al., 2010). Intriguingly, a subset of mesoderm progenitors appeared to be committed to HE cells as indicated by the expression of endothelial lineage genes (*CDH5*, *CD34*, *KDR*, and *PECAM1*; cluster HE) (**Figures 3D and S5C**). Gene expression profiles suggested that these HE cells were primed to undergo the segregation process to generate endothelial and hematopoietic cells since our flow cytometry analysis demonstrated that most CD144^+^ cells on day 6 were CD73^−^, while cells in cluster HE were already found to express *NT5E* (CD73) (**Figures 3D and S5C**). Additionally, HE cells also expressed *SOX17* and *SOX7*, which are key genes for the transition to hematopoietic cells (Lilly et al., 2016; Nakajima-Takagi et al., 2013) (**Figures 3D and S5C**). Recent studies demonstrated that Notch signaling plays a pivotal role to undergo the EHT process and to induce definitive hematopoiesis from hPSCs (Jang et al., 2015; Leung et al., 2018; Uenishi et al., 2018). Since organoid-derived HPCs possessed definitive hematopoiesis potential, we attempted to determine whether Notch signaling-associated genes are upregulated in HE cells. Notably, both Notch ligands (*JAG1*, *JAG2*, and *DLL4*) and receptors (*NOTCH1*, *NOTCH3*, and *NOTCH4*) were highly expressed in cluster HE (**Figure 3E**), suggesting that auto-activation of Notch signaling promotes the transition process and a role in the induction of definitive hematopoiesis. Despite the early loss of hBM-MSCs and the lack of extrinsic factors to support organoid development, the early hemato-vascular development was precisely regulated during organoid formation. We hypothesized that the critical morphogens required for this stage may be provided from hiPSC-derived cells in organoids. Genes encoding morphogens known to be essential for hemato-vascular development, such as BMP4 (*BMP4*), FGF2 (*FGF2*), VEGF (*VEGFA*), SCF (*KITLG*), and TGFβ-1 (*TGFB1*), were highly expressed in hiPSC-derived cells (Kennedy et al., 2012; Ledran et al., 2008; Woods et al., 2011; Yu et al., 2012) (**Figure 3E**). Our observation raised the possibility that these important morphogens and the signaling required for early embryonic development in the yolk sac can be provided by hiPSC-derived cells reciprocally. In addition to mesoderm progenitors and their derivatives, we identified a unique cell type within mesodermal lineage cells in day 6 organoids. Cells in this cluster expressed trophoblast-associated genes, such as *HAND1*, *CDX2*, *KRT7*, *TFAP2A/2C*, *GATA3*, *CDH1*, and *CXCL12*, and were annotated as trophoblast-like cells (cluster TB) (Dong et al., 2020; Kubaczka et al., 2015; Marchand et al., 2011; Okae et al., 2018) (**Figures 3D and S4E**). Interestingly, genes expressed in the late stage of trophoblast development, such as *CGA*, *CGB*, and *HLA-G*, were not enriched in TB cells (Dong et al., 2020; Xu et al., 2002).

scRNA-seq analysis of day 6 organoids revealed a cell complement reminiscent of the cellular components of the developing human definitive yolk sac, including mesoderm progenitors, HE cells, and yolk sac endoderm. Gene expression profiling revealed that signaling pathways required for early hemato-vascular development were recapitulated in the organoids.

### Various hematopoietic cell subsets are detected in day 13 organoids

Unsupervised clustering of day 13 cells revealed 12 transcriptionally distinct clusters (Figure 4A). Similar to day 6 organoids, we detected yolk sac endoderm cells (cluster YE) and trophoblast-like cells (cluster TB). Approximately 70% of cells in day 13 organoids were stroma-like cells (*CD44*, *PDGFRB*, and *COL1A1*). Remarkably, these stromal-like cells expressed *HAND1/2* and *FOXF1*, indicating that they are derivatives of mesoderm progenitors in an early mesoderm stage (Figures 4B and S5A). We classified these stromal-like cells into three different groups; early mesenchymal cell (*PDGFRA*, *PDGFRB*, and *THY1*; clusters MS), stromal cell (*CD44*, *PDGFRB*, *COL1A1*, and *DCN*; ST clusters), and SLC transport genes expressing stromal cell (cluster SLC) (Figures 4B and S5B). SLCs are transmembrane transporters that are widely expressed in the human yolk sac to import various nutrients from the exocoelomic cavity (Cindrova-Davies et al., 2017). Notably, various SLC family genes were enriched in the top 10 of the SLC cluster specific genes (Figure S5B; Table S2).

**Figure 4.**
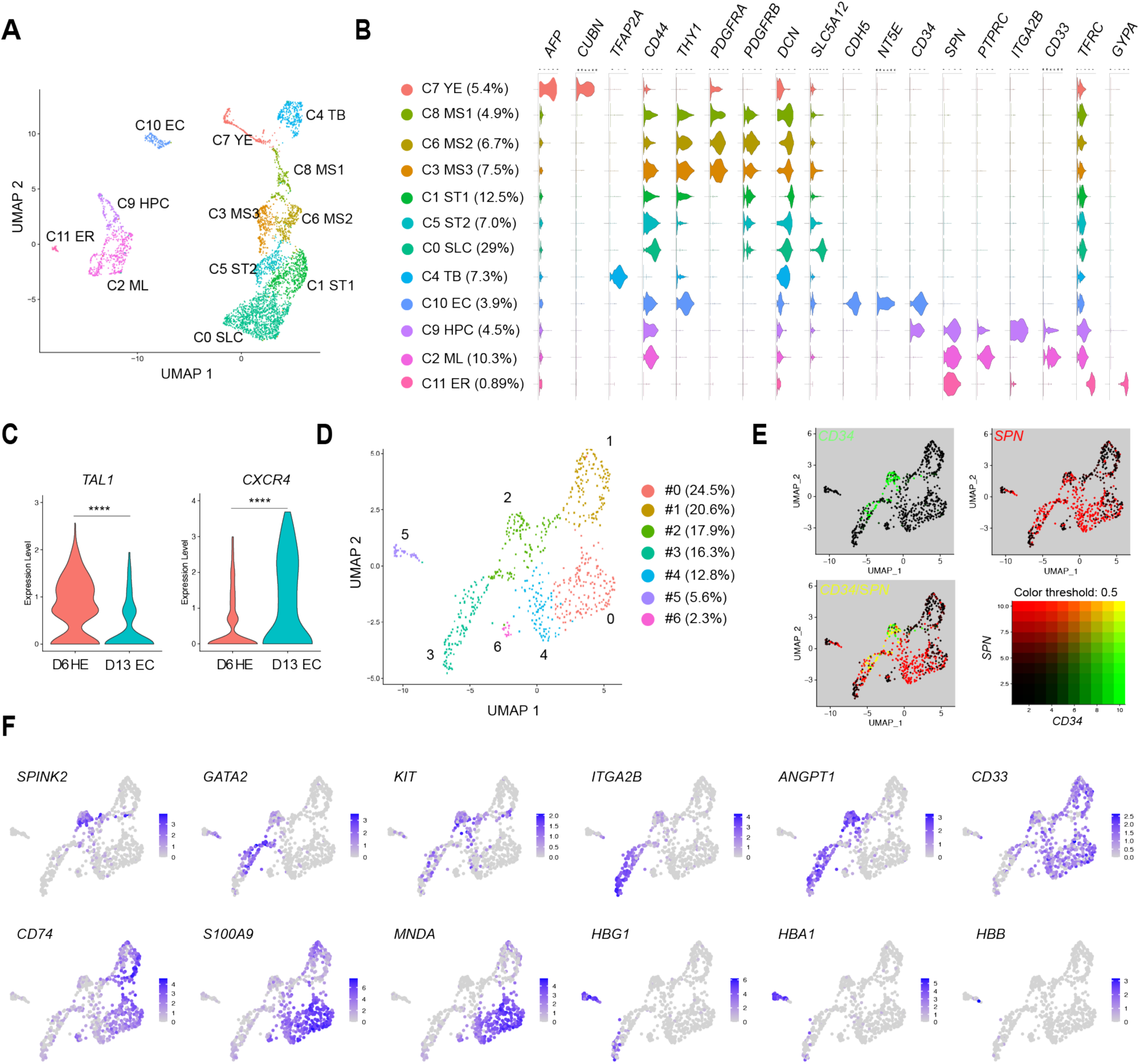
Various types of hematopoietic cells are present in day 13 organoids. (A) UMAP plot displaying scRNA-seq analysis of 4,401 GFP^+^ cells isolated from day 13 organoids. (B) Violin plots showing expression of representative lineage marker genes for day 13 clusters. (C) Comparison of the gene expression levels of *TAL1* and *CXCR4* between day 6 HE cells and day 13 EC cells. (****p < 0.0001) (D) UMAP plot displaying sub-clusters of hematopoietic lineage cells in day 13 organoids. (E) Co-expression of *CD34* and *SPN* within hematopoietic sub-clusters on UMAP. Cells in UMAP are colored according to the expression level of *CD34, SPN*, and both genes. An expression cutoff of 2.5 is used for visualization. (F) Feature plots of expression distribution for various hematopoietic lineage genes within hematopoietic sub-clusters. Expression levels for each cell are color-coded and overlaid onto the UMAP plot.

Consistent with the peak of HPC generation, we identified three *SPN* (CD43) expressing hematopoietic clusters and an endothelial cluster (*KDR*, *PECAM1*, *CD34*, and *NT5E*: EC cluster) in day 13 organoids, indicating that HE cells successfully accomplished the transition process and generated hematopoietic cells and more mature endothelial cells (**Figures 4B and S5C**). As reported in previous studies, the mature endothelial cells showed lower expression of *TAL1* and higher *CXCR4* expression when compared to HE cells (**Figure 4C**) (Choi et al., 2012; Ditadi et al., 2015). Hematopoietic lineage cells were tentatively classified into multipotent HPCs (expressing both *CD34* and *SPN*; cluster HPC), myeloid cells (*SPN*, *CD33* and, *PTPRC*; cluster ML), and erythroid cells (*GYPA* and *TFRC*; cluster ER) (**Figure 4B**). In the murine yolk sac, the first definitive hematopoietic cells emerge as erythro-myeloid progenitor (EMP), then followed by HPCs with lymphoid-myeloid potential (Palis et al., 1999; Palis and Yoder, 2001). These two types of definitive hematopoietic progenitor cells can also be observed during hPSC differentiation. Generally, hPSC-derived EMP cells can be detected by co-expression of CD41a (*ITGA2B*) and CD235a (*GYPA*), while multipotent HPCs do not express these lineage markers but express CD34 and CD43 (*SPN*) (Klimchenko et al., 2009; Slukvin, 2013; Vodyanik et al., 2006). Contrary to this, high expression of *ITGA2B* was detected in the HPC cluster and *GYPA* expression was observed only in the erythroid cluster (**Figure 4B**). Since hPSC-derived multipotent HPCs have been reported as a heterogenetic population (Fidanza et al., 2020), we sought to explore the heterogeneity within the HPC cluster. We performed a sub-cluster analysis for cells from the three hematopoietic clusters which uncovered seven hematopoietic subsets (**Figure 4D; Table S4**).

Co-expression of *CD34* and *SPN* was mainly found in sub-cluster 2 (**Figure 4E**) and cells in this cluster also expressed HSC specific genes, such as *SPINK2*, *KIT*, and *SOX4* (**Figures 4F and S6**) (Fidanza et al., 2020; Zeng et al., 2019). Previous reports demonstrated that early hematopoietic cells derived from HE cells express CD41a (*ITGA2B*) (Garcia-Alegria et al., 2018; Mitjavila-Garcia et al., 2002), which we detected in cells of sub-cluster 2 and 3 with stronger expression in subcluster 3 (**Figure 4F**). Notably, early hematopoietic specific genes, such as *GATA2*, *KIT*, *MEIS1*, and *ANGPT1*, were also expressed in sub-cluster 3 (**Figures 4F and S6**), suggesting early hematopoietic progenitor states for these cells (Azcoitia et al., 2005; Castaño et al., 2019; Sitnicka et al., 2003; Suri et al., 1996; Zeng et al., 2019). Interestingly, an early T lymphoid lineage gene *CD7* expression could be detected in sub-clusters 2 and 3, again supporting the definitive hematopoietic potential of the organoid (**Figure S6**). Myeloid lineage genes, such as *CD33*, *CD74*, *S100A9*, and *MNDA*, were expressed in sub-cluster 0, 1, 4 and 6 (**Figures 4F and S6**), suggesting myeloid lineage cell states. Sub-cluster 5 was positive for erythroid cell associated genes including *GYPA*, *KLF1*, and hemoglobin genes *HBA1*, *HBE1*, and *HBG* (**Figures 4F and S6**). The gene expression patterns of hemoglobin suggested that most erythroid lineage cells in day 13 organoids were primitive or fetal type because *HBB* (adult type-beta globin gene) expression was barely observed (**Figures 4F and S6**). Collectively, gene expression data indicated that day 13 organoids contain myeloid and erythroid lineage committed hematopoietic cells as well as hematopoietic progenitor cells in different states. In mice, various hematopoietic progenitor cells can be identified in the yolk sac prior to the emergence of the adult-type HSCs in the embryo proper (Yamane, 2018). Similarly, our yolk sac-like organoids induced a heterogeneous population of hematopoietic progenitors from hiPSCs.

Overall, our scRNA-seq analyses for day 6 and day 13 organoids captured cells with a human definitive yolk sac-like composition. These findings suggested that the organoid system can serve as an *in vitro* model to study human yolk sac endoderm functions and the interactions between endoderm and mesoderm during embryonic hematopoiesis.

### Stromal cells induced from hiPSCs can be used for the yolk sac-like organoid system

Our approach to generate yolk sac-like organoid is not only a useful study model to address the developmental biology of the human yolk sac, but also has a great potential to provide cost-effective bulk production of hiPSC-derived HPCs for therapeutic use. However, the *in vitro* expansion of hBM-MSCs induces cell senescence and reduces their stemness (Jiang et al., 2017). Moreover, isolation of hBM-MSCs is an invasive procedure requiring clinical justification. Accordingly, we sought to identify alternative cell types that are more easily accessible and can be readily expanded. Since Hp-spheroids can induce early stromal cells and continue their organogenesis after the loss of hBM-MSCs, we speculated that stromal cells generated from hiPSCs might be a potential alternative to hBM-MSCs. To address this, we generated hiPSC-derived stromal cells (iSTCs) from NCRM5 cells (GFP^−^ cells) according to our previous report (Takebe et al., 2017) (Figure S7A). Flow cytometry analysis demonstrated that during their differentiation, iSTCs increased the expression of typical stromal cell markers, such as CD44, CD105, and CD73. However, by day 20 of induction, the expression levels of CD105 and CD73 were much lower than CD44 (Figure S7B). This was consistent with our kinetic flow cytometry analysis of CD44 and CD73 during organogenesis (Figure S3B). Based on the expression pattern of cell surface markers, iSTCs were still in an immature state until day 20. Nevertheless, all types of iSTCs isolated from various induction points (day 8, day 14, and day 20) can support the generation of HPCs in organoids with similar proficiency to using hBM-MSCs (**Figure S7C**). Our data indicate that a wide range of stromal cells, from hBM-MSCs to early stromal cells, can be used for yolk sac-like organoid formation. Since patient-specific hiPSCs can produce an unlimited number of iSTCs, combining the use of bioreactors and iSTCs allows for establishing an industry-scaled yolk sac-like organoid system.

### HPCs generated in autologous yolk sac-like organoids give rise to various blood cell lineages

To further evaluate the potential of our yolk sac-like organoid system for clinical application of iPSC-based blood cell therapy, we sought to determine whether HPCs generated by our modified iSTCs induced organoid (iSTC-organoid) system could be differentiated to erythroid cells, macrophages, and T lymphocytes by current maturation protocols. To this end, we used a patient-specific hiPSC line, Mart1-iPSC, that was generated from a cytotoxic T cell possessing T cell receptor (TCR) specific for the melanoma epitope Mart1 (Vizcardo et al., 2013). Mart1-iPSC-derived HPCs were harvested on day 13 and subjected to further differentiation toward these targeted blood cell lineages (Figure 5A). Similar to NCRM5-EGFP cells, Mart1-iPSC cells also expressed HPC markers using the iSTC-organoid system (Figure 5B). For erythroid and macrophage differentiation, CD34^+^ cells were isolated with magnetic-activated cell sorting (MACS). After 15 days of erythroid differentiation, the cell pellets turned red (Figure 5C) and flow cytometry analysis of CD71 and glycophorin A (GPA) revealed that these cells contained CD71^+^GPA^+^ cells (erythroblasts) and CD71^−^GPA^+^ cells (mature erythroid cells) (Figure 5D). Globin gene expression patterns in Mart1-iPSC-derived erythroid cells were determined by quantitative RT-PCR. We found the dominant globin expression pattern in Mart1-iPSC-derived erythroid cells included α-globin and γ-globin, indicating most of them were fetal type erythroid cells, but not of the primitive type (Figure 5E). After culturing under macrophage differentiation conditions for 14 days, Mart1-iPSC-derived CD34^+^ cells generated cells with typical human macrophage cell markers (Figure 5F) and morphology (Figure 5G). Additionally, normal phagocyte activities including phagocytosis of zymosan particles and production of reactive oxygen species were observed in Mart1-iPSC-derived macrophages upon stimulation with phorbol 12-myristate 13-acetate (PMA) (Figures 5H and 5I). Finally, we confirmed that Mart1-iPSC-derived HPCs co-cultured with OP9-DLL1 cells for 21 days differentiated into mature CD4^+^CD8^+^ double positive T cells with a high-population of CD3^+^ T cells recognizing the Mart1-tetramer (**Figure 5J**). As reported in previous studies, antibody-driven TCR stimulation of these CD4^+^CD8^+^ double positive T cells induced CD8^+^ single positive T cells (**Figure 5K**). Function and antigen specificity of these Mart1-iPSC derived CD8^+^ single positive T cells were confirmed by cytokine release after co-culture with Mart1 peptide pulsed T2 cells (Maeda et al., 2016; Vizcardo et al., 2013) (**Figure 5L**).

**Figure 5.**
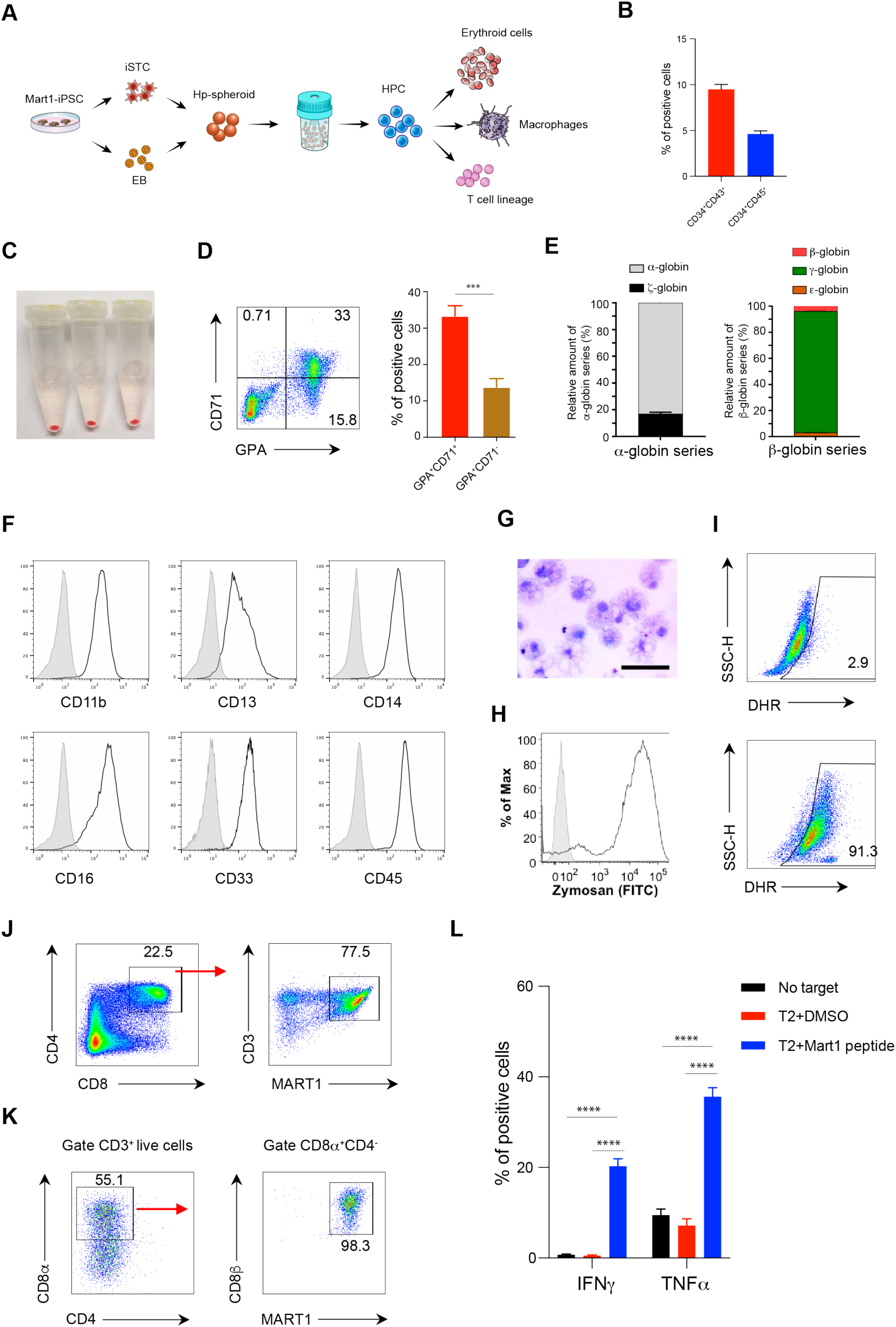
HPCs generated in iSTC-organoids can undergo maturation to erythroid cells, macrophages, or T lymphocytes using current cell maturation protocols. (A) Schematic outline illustrating the use of Mart1-iPSC for iSTC-organoids to generate erythroid cells, macrophages, and T lymphocytes. (B) Quantifications of the percentage of CD34^+^CD43^+^ and CD34^+^CD45^+^ cells in Mart1-iPSC derived iSTC-organoids on day 13. Values represent mean ± SD (n = 3) (C-E) Mart1-iPSC derived HPCs cultured in the erythroid differentiation condition for 15 days were harvested to characterize erythroid cell properties. The color of cell pellets changes to red (C), and typical erythroid lineage markers, such as CD71 and GPA, can be detected by flow cytometry analysis (D). Globin expression patterns are determined by relative RNA expression of α-, ζ-globin (α-globin series) and ε-, γ-, and β-globin (β-globin series) (E). Values represent mean ± SD (n = 3). (F-I) Macrophage differentiation from Mart1-iPSC derived CD34^+^ cells. (F) Macrophage surface markers are analyzed by flow cytometry (gray is isotype control). (G) Giemsa stain of macrophage cytospin. Scale bar = 50 um. (H) Phagocytosis assay of FITC-labeled Zymosan A particles (gray is isotype control). (I) DHR assay of reactive oxygen species production by Mart1-iPSC derived macrophages in response to stimulation with PMA (lower panel); unstimulated macrophage control is also shown (upper panel). (J–L) T lymphocytes generated from Mart1-iPSC derived HPCs. After 22 days of T cell differentiation, CD4^+^CD8^+^ double positive T cells can be detected by flow cytometry analysis, most of which are CD3^+^Mart1-tetramer^+^ cells (J). Mart1-iPSC derived CD4^+^CD8^+^ double positive T cells can be induced to CD8αβ single positive T cells by stimulation with Mart1 peptide primed T2 cells (K). Mart1-iPSC derived CD8^+^ single positive T cells release INFγ and TNFα in response to Mart1 peptide pulsed T2 cells (L). T cell responses without T2+Mart1 peptide (no target) or without Mart1 peptide (T2+DMSO) are also shown. Values represent mean ± SD (n = 3, ****p < 0.0001).

To exclude the possibility that this approach was only applicable to some hiPSC lines, we also analyzed a sickle-cell disease derived iPSC cell (SCD-iPSC) line (Uchida et al., 2017). Similar to NCRM5-EGFP cells, SCD-iPSC cells also generated HPCs with T lymphocyte potential (**Figure S7D**). Therefore, our data demonstrate that HPCs generated by our iSTC-organoid system are able to successfully produce erythroid, myeloid, and T lymphoid lineage cells.

### The iSTC-organoid system can be performed in a xeno-free condition

For the therapeutic use of iPSC-based cell therapies, production of cells under xeno-free conditions is desirable due to biosafety issues. Currently, there are many xeno-free media formulations available, and we screened several commercial xeno-free media designed for culturing MSCs or hematopoietic cells available at the time of this study, but none of them supported the iSTC-organoid system (data not shown). Therefore, we set out to develop a xeno-free medium for our iSTC-organoid system (Figure 6A). We hypothesized that high doses of growth factors/cytokines supplemented in available media disrupt the physiological diffusion of morphogens in the developing organoids and may disturb the organogenesis. Therefore, we used αMEM as a base medium and screened commercial FBS replacements. We found that αMEM containing 2.5% PLTGold (heparin-free human platelet lysate) and StemfitCo2 (an FBS-replacement kindly provided from Ajinomoto Co., Inc) supported the iSTC-organoid system in generating CD34^+^CD43^+^ cells in bioreactors similar to using FBS (Figures 6B and 6C). To assess the definitive hematopoietic potential of HPCs generated by this condition, we conducted T cell differentiation as described above. Similar to the conditions of FBS, HPCs derived from NCRM5-EGFP (non-T cell derived hiPSCs) under the xeno-free condition were induced to become T cell progenitors and CD4^+^CD8^+^ double positive T cells (**Figures 6D and 6E**). As reported previously, the differentiation of non-T cell derived iPSCs do not show the precocious CD3 expression observed in TCR pre-rearranged iPSCs (Vizcardo et al., 2018). During T cell development, CD3 expression is concomitant with TCR expression after TCR gene rearrangements. Therefore, NCRM5-EGFP hiPSC showed a spike peak in the production of CD3^+^ cells in CD4^+^CD8^+^ double positive T cells around day 30 (**Figures 6F and 6G**). Moreover, there was no significant difference in T cell differentiation of HPC generated in either xeno-free medium or FBS containing medium (**Figure 6H**). Measuring TCR diversity in CD3^+^ cells isolated from CD4^+^CD8^+^ double positive T cells indicated successful TCRβ rearrangement in NCRM5-EGFP derived T lineage cells (**Figure 6I**). Taken together, the iSTC-organoid system applied in a xeno-free condition performed well in generating HPCs with definitive hematopoiesis. Our data suggested that the iSTC-organoid system can provide a simple, robust, and scalable approach to generate hiPSC-derived HPCs in a xeno-free condition.

**Figure 6.**
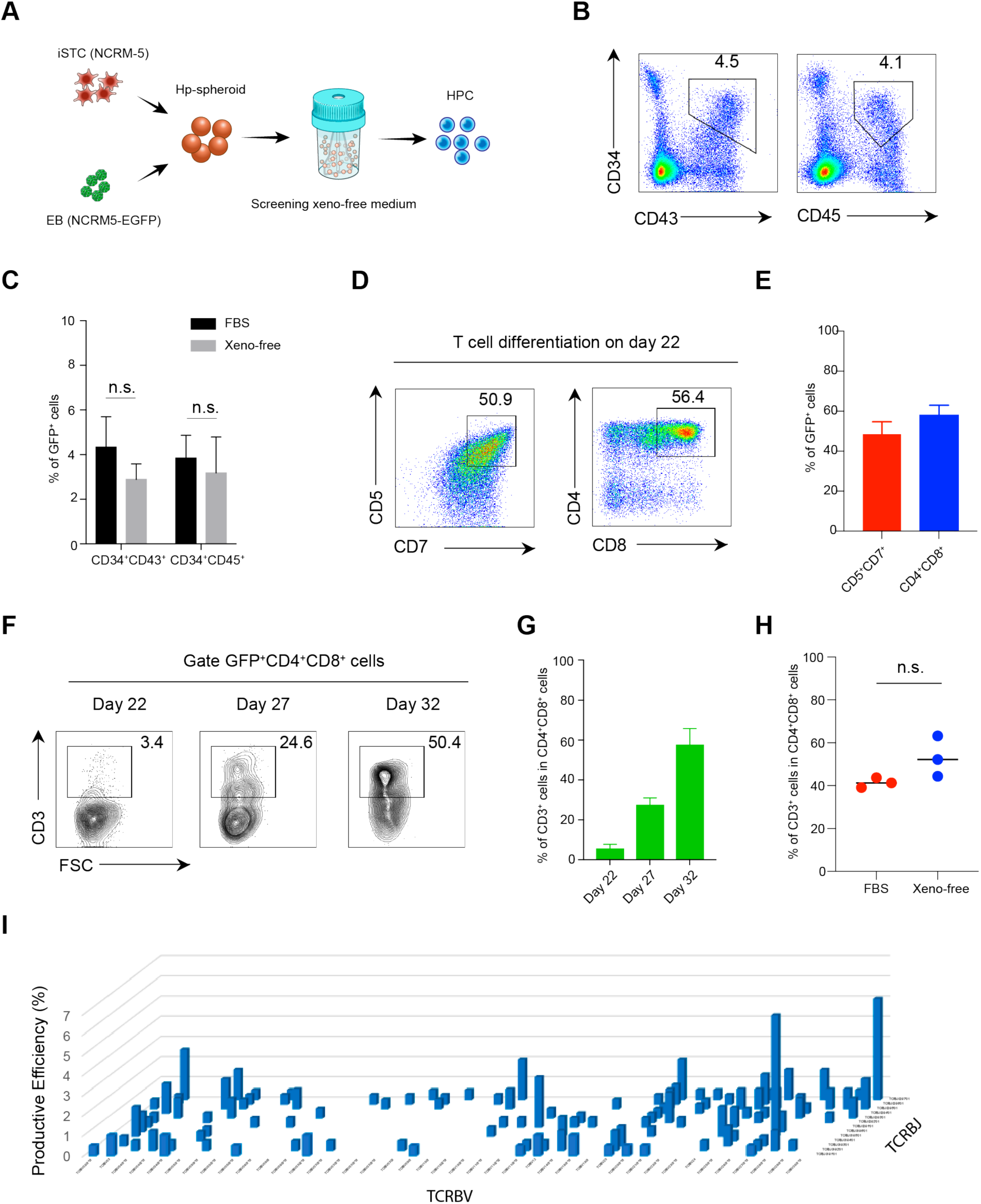
The iSTC-organoid system can be performed in a xeno-free condition. (A) Schematic outline of screening a xeno-free condition for the iSTC-organoid system. (B) Representative flow cytometry analysis of CD34^+^ and CD43^+^ cells in iSTC-organoids cultured in the xeno-free medium for 13 days. GFP^+^ cells are gated for analysis. (C) Comparison of the frequency of CD34^+^CD43^+^ cells and CD34^+^CD45^+^ cells in iSTC-organoids between culturing in FBS-based medium versus xeno-free medium. GFP^+^ cells are gated for analysis. Values represent mean ± SD (n = 3, n.s., not significant). (D) Representative flow cytometry analysis of T lineage cell markers, including CD5, CD7, CD4, and CD8, from differentiation of HPCs generated in xeno-free condition. Cells were harvested at day 22 of T cell differentiation. GFP^+^ cells are gated for analysis. (E) Quantifications of flow cytometry analysis in (D). GFP^+^ cells are gated for analysis. Values represent mean ± SD (n = 3). (F) Representative flow cytometry analysis of CD3^+^ cells in CD4^+^CD8^+^ cells during the T cell differentiation. GFP^+^ cells are gated for analysis. (G) Quantifications of flow cytometry in (F). Values represent mean ± SD (n = 3). (H) Quantifications of the frequency of CD3^+^ cells in CD4^+^CD8^+^ cells on day 32 of T cell differentiation. GFP^+^ cells are gated for analysis. Horizontal bars represent mean value (n = 3, n.s., not significant). (I) TCR sequence analysis showed the diversity of rearranged TCR genes in NCRM5-EGFP derived CD4^+^CD8^+^CD3^+^ T cells isolated from day 32 of the T cell differentiation.

## DISCUSSION

Embryonic development is a very complex and intricate process that requires cells to act in a coordinated manner. Each cell in the embryo receives signals from neighboring cells in the form of morphogens or direct cell-cell contacts to regulate its fate. Here, we demonstrate two important new findings.

Firstly, hiPSCs spontaneously form yolk sac-like organoids in the absence of any extrinsic factors when they are co-cultured with stromal cells as Hp-spheroids. Despite the early loss of co-cultured stromal cells, hiPSCs continued to develop into yolk sac-like organoids and generate multipotential HPCs. For this to happen it is possible that hiPSCs (EBs) and stromal cells are separated on different sides in the Hp-spheroid, resulting in a gradient of morphogens secreted by the stromal cells, creating a polarity in the Hp-spheroids that resembles the initial polarity in embryonic tissue (Ashe and Briscoe, 2006). For example, the gradient of BMP4 has a great impact on mesodermal tissue formation and early hematopoietic development during embryogenesis (Caroline J. Marshall, 2000; Sadlon et al., 2004). However, for hiPSC differentiation *in vitro*, it is difficult to experimentally extrapolate the use of artificial morphogens to faithfully mimic the natural gradient of morphogens present in embryonic tissues.

Our organoid system is a highly reproducible approach to generate definitive HPCs across experiments and different hiPSC cell lines, even though no stage-specific exogenous factors were used to control hiPSC differentiation. Importantly, our organoid induced hematopoiesis followed the hemato-vascular development observed in yolk sac mesoderm. A number of current approaches to generate hiPSC-derived organoids require exogenous factors to regulate their differentiation and morphogenesis until hiPSC-derived organ-specific stem cells initiate self-assembly and organogenesis (McCauley and Wells, 2017). However, embryonic tissue formation is regulated by crosstalk and signaling between multiple germ layers, therefore the use of artificial morphogens may mask these interactions and limit the observation of innate embryogenesis *in vitro*. Without the use of a complicated cocktail of factors, our yolk sac-like organoid system provides a simple and more physiological study model for the investigation of the developmental biology of the human definitive yolk sac.

Dissection of day 6 and day 13 organoids with scRNA-seq analysis revealed that the yolk sac-like organoids are composed of three groups of cells; mesoderm progenitors, trophoblast-like cells, and yolk sac endodermal cells. The mesoderm progenitors eventually give rise to hemato-vascular and stromal lineages cells during the organoid development. These mesoderm progenitors and their derivatives as well as yolk sac endodermal cells can be observed in the process of human definitive yolk sac development. It has been found that human yolk sac endoderm takes up nutrients by endocytosis and shuttles them to the embryo through blood vessels or yolk sac cavity (Burton et al., 2001; Pereda and Motta, 1999; Ross and Boroviak, 2020; Zohn and Sarkar, 2010). In line with this, endodermal cells in yolk sac like organoids expressed genes required for megalin and cubilin mediated endocytosis and various apolipoprotein genes to transport cholesterol, lipids, and lipid soluble vitamins. Additionally, similar to mouse yolk sac endoderm, they also expressed *IHH* and *VEGFA*, which promote the specification of HE cells from mesoderm progenitors. Our hiPSC-derived endodermal cells are therefore intriguingly similar to human and mouse yolk sac endoderm cells. Our observations suggest that these yolk sac-like organoids can be used to study the molecular biology of human yolk sac endoderm and its functions.

Secondly, our approach to generate yolk sac-like organoids offers several advantages for clinical application in the context of hiPSC-based blood cell therapy. scRNA-seq analysis of day 13 organoids demonstrated that our organoid system induces various types of hematopoietic cells, including myeloid, erythroid lineage cells and multipotential HPCs. Importantly, organoid-derived HPCs can be differentiated into erythroid cells, macrophages, and T lymphocytes with current maturation protocols, indicating that our organoid system is adaptable to most of such maturation protocols. Since the expansion of hiPSC-derived HPCs is still challenging, the bulk production of HPCs is an important step toward the therapeutic use of hiPSC-derived blood cells. Currently, generation of hiPSC-derived HPCs in a xeno-free condition requires a complicated cocktail of cytokines/growth factors, and severely limits the establishment of a cost-effective bulk production of HPCs. Our organoid system overcomes these restrictions and does not require GMP (good manufacturing practices) grade cytokines and growth factors. The combination of iSTCs and bioreactors allows an industrial scale production of HPCs from patient-specific hiPSCs. Our results demonstrated that iSTCs isolated from day 8 of induction can be used in the organoid system just as effectively as hBM-MSCs, suggesting that a large number of stromal cells required for a clinical-scale organoid system can be prepared easily and quickly from patient-specific iPSCs. Notably, in contrast to the classical OP9 co-culture system, our system uses pre-aliquoted iSTCs directly for Hp-spheroid formation, allowing us to save time and manpower to prepare stromal cells.

In conclusion, we developed a simple, scalable system to generate yolk sac-like organoids that are structurally and functionally similar to the human definitive yolk sac, without using any exogenous factors. Our yolk sac-like organoid system can be used as a model to study the molecular biology of human definitive yolk sac, including the functions of yolk sac endoderm and the hemato-vascular development in the human yolk sac. Importantly, this system will also pave a new avenue to the clinical application of hiPSC-based blood cell therapy.

## Supporting information

Sup analysis1

Sup analysis2

## ACKNOWLEDGEMENTS

For technical support, we thank Zhiya Yu, Rafiqul Islam, Ken-ichi Hanada, Marta Bosch-Marce, Minh Tran, and Chengyu Liu. We thank Arnold Mixon and Shawn Farid for flow cytometry support; and Eric Tran, Rigel Kishton, Madhusudhanan Sukumar, Tori Yamamoto, Douglas Palmer, Ping-Hsien Lee, Devikala Gurusamy, Amanda Henning, and Sherif Badr for critical feedback. We thank Hiroshi Kawamoto and Kyoko Masuda for kindly providing the OP9/DLL1 cell line. We thank Erina He and Maria Romanova for graphic support; Celina Juliano for access to computational resources. Finally, we thank Francis Flomerfelt, for advice, feedback, and support. This research was supported by the Intramural Research Programs of the NCI (ZIA BC010763), NIH/NCI (K08CA197966), NIAID (Z01 AI000644), and NINDS (ZIA NS003034). This work was also supported by the Tiens Charitable Foundation, the NIH Center for Regenerative Medicine, the Milstein Family Foundation, and the Melanoma Research Alliance.

## AUTHOR CONTRIBUTIONS

N.T. and R.V. conceived the project, designed the experiments, collected and analyzed the data, and wrote the manuscript. S.S. conducted bioinformatics analysis and edited manuscript. J.J.H. and N.U. performed erythroid cell differentiation with expert advice from J.F.T.. T.M., and M.L.G. performed T cell differentiation. C.L.S., U.C., J.B. and S.K. performed macrophage differentiation with expert advice from H.L.M.. Y.H., and R.I. performed imaging analysis with expert advice from D.S.G. and M.J.K.. N.H., S.K.V., Y.Y., M.K., T.T., J.Z., D.F.S., and P.G.R. helped to interpret experimental results and provided advice regarding the research strategy. N.P.R. supervised the project.

