## Supplemental information for "Self-organized yolk sac-like organoids allow for scalable generation of multipotent hematopoietic progenitor cells from human induced pluripotent stem cells"

#### METHOD DETAILS

##### Human iPSC maintenance

NCRM5 and NCRM5-AAVS1-CAG-EGFP (NCRM5 iPSCs with CAG-EGFP integrated at Chr.19 AAVS1 safe harbor locus) were established by the iPSC core facility of the National Heart, Lung and Blood Institute in NIH. Mart1-iPSC were kindly provided by Riken BBC, in Japan. SCD-iPSC line was established from bone marrow stromal cells isolated from a sickle cell disease patient (Uchida et al., 2017). All hiPSCs were cultured on Matrigel (Corning) or iMatrix-511 (Nippi) coated dishes in xeno-free hiPSC medium Essential 8 (Invitrogen) or StemFit (Ajinomoto Co. Inc). They were routinely passaged as small clumps/single cells using 0.5 mM EDTA in phosphate buffered saline (PBS) with the split ratio of 1:6 to 1:10 every 3 to 4 days after reaching 65% to 80% confluence. After EDTA treatment, hiPSCs were transferred to new Matrigel or iMatrix-511 coated dishes in hiPSC medium supplemented with ROCK inhibitor Y-27632 (10 μM, R&D Systems Inc.). Next day, the medium was changed to hiPSC medium without ROCK inhibitor.

##### Preparation of hBM-MSCs for co-culturing with hiPSCs

Healthy donor derived hBM-MSCs were kindly provided from the Cell Processing Section, Department of Transfusion Medicine in NIH. Establishment of hBM-MSCs is described in our previous study (Sabatino et al., 2012). hBM-MSCs in cryobags (100 x $10^6$ cells/bag) were prealiquoted into cryotubes (5 x $10^6$ cells/tube) with CELLBANKER (AMSBIO) for Hp-spheroid system.

**EBs differentiation to HPCs with a cytokine/growth factor cocktail**

Semi-confluent hiPSC were dissociated with 0.5 mM EDTA in PBS and transferred to Aggrewell 400 (6 well plate type, 1 x $10^6$ cells/ well, STEMCELL Technologies) for EB formation (100-150 cells/EB) in hiPSC medium supplemented with ROCK inhibitor Y-27632 (10 μM) according to the manufacturer's protocol. To compare with the Hp-spheroid system, the differentiation of EBs with exogenous factors was performed according to the previous publication with some modifications (Kennedy et al., 2012). In brief, EBs were transferred from Aggrewell 400 plates to ultra-low attachment 3D culture plates (96 well U bottom, S-BIO) and were cultured in αMEM medium (Invitrogen) with 20% fetal bovine serum (FBS) and exogenous factors. Human cytokines and growth factors were added as follows: bone morphogenic protein 4 (BMP4, 10 ng/ml, R&D Systems) and fibroblast growth factor 2 (FGF2, 20 ng/ml, R&D Systems) from day 0 to day 3, FGF2 (20 ng/ml) from day 3 to day 5, FGF2 (20 ng/ml), vascular endothelial growth factor (VEGF, 10 ng/ml, R&D Systems), interleukin 3 (IL3, 10 ng/ml, R&D Systems), interleukin 6 (IL6, 10 ng/ml, R&D Systems), Flt3-Ligand (Flt3-L, 10 ng/ml, R&D Systems) and stem cell factor (SCF, 100 ng/ml, R&D Systems) from day 5 to day 8, VEGF (10 ng/ml), IL3 (10 ng/ml), IL6 (10 ng/ml), Flt3-L (10 ng/ml), SCF (100 ng/ml), and thrombopoietin (TPO, 10 ng/ml, R&D Systems) from day 8 to day 13. Medium was changed every day.

**HPCs generation from hiPSCs in 2D**

The classic OP9 co-culture system in 2D for HPC generation was performed according to previous publication with a slight modification (Vizcardo et al., 2013). In brief, OP9 cells were cultured on gelatin coated 10 cm dishes for 7-8 days before co-culturing with hiPSCs. To place similar size of hiPSC clumps on OP9 dishes, we harvested hiPSC and formed EBs (2.5 x $10^3$ cells/EB) in Aggrewell 800 plates (24 well plate type, STEMCELL Technologies) in hiPSC medium supplemented with ROCK inhibitor Y-27632 (10 μM) in advance. Next day, we transferred EBs onto gelatinized OP9 dishes in OP9 medium consisting of αMEM with 20% FBS. On day 13 of the differentiation, hiPSC-derived sac-like structures were enzymatically dissociated for further analysis. To prepare gelatinized hBM-MSC dishes, we transferred hBM-MSCs onto gelatin-coated 10 cm dishes within one passage after thawing and cultured them in OP9 medium for 7-8 days before co-culture with hiPSCs.

**Generation of yolk sac-like organoids from hiPSCs with Hp-spheroids system in bioreactors**

To form Hp-spheroids, small EBs (100-150 cells/EB) were mixed with thawed stromal cells (hBM-MSCs or OP9 cells or iSTCs) to make cell suspension in αMEM with 20% FBS. Cell suspension was transferred to Aggrewell 800 plates and then was immediately settled into microwells of Aggrewell 800 by centrifuge (800rpm, 3min). Next day, Hp-spheroids were transferred to bioreactors (ABLE Bioreactor Magnetic Stir System 30 mL, REPROCELL), stirring at 65 rpm, in αMEM with 20% FBS. Medium was changed for every 2-3 days. Generally, we used 4 x $10^6$ hiPSCs for EBs formation and mixed with 5 x $10^6$ thawed stromal cells to form 2400-2800 Hp-spheroids for one bioreactor. We used αMEM containing 2.5% PLTGold (Mill Creek Life Sciences) and 1x StemFitC02 (kindly provided by Ajinomoto Co. Inc) as xeno-free medium for the iSTC-organoid system. EBs and thawed iSTCs were mixed in the xeno-free medium supplemented with ROCK Inhibitor Y-27632 (10 μM) for Hp-spheroids formation. The next day, Hp-spheroids were transferred to bioreactors in the xeno-free medium without ROCK inhibitor.

**Dissociation of yolk sac-like organoids**

To dissociate organoids, we washed with Hank's Balanced Salt Solution (HBSS) once and then treated with HBSS containing Liberase TM (Roche, 250 μg/ml) and DNase I (Roche, 500 μg/ml) at 37°C for 15-20 minutes, followed by pipetting for 1 minute with a P1000 disposable plastic tip to break organoids. Cells were then washed with PBS once and treated with PBS containing 10% TrypLE Express (Gibco) and DNase I (500 μg/ml) at 37°C for 10-15 minutes, followed by pipetting for 1 minute with a P1000 disposable plastic tip to make a single cell suspension.

**Induction of stromal cells from hiPSCs**

Generation of iSTCs was performed according to our previous study with a slight modification (Takebe et al., 2017) as summarized in **Figure S7A**. In brief, hiPSCs were plated on iMatrix-511 coated dishes in hiPSC medium supplemented with ROCK inhibitor Y-27632 (10 μM) before differentiation. We used DMEM/F12 (Gibco) containing 1x StemFit for Differentiation (AS401, Ajinomoto Co. Inc) and 1% Glutamax as a basal medium. For mesoderm induction stage, hiPSCs were cultured in the basal medium supplemented with CHIR99021 (8 μM, Tocris Bioscience) and BMP4 (25 ng/ml). Medium was changed every day until day 3 of induction. On day 4, medium was changed to stromal cell induction medium consisting of the basal medium supplemented with Activin A (2 ng/ml, Peprotech Inc.) and PDGF-BB (10 ng/ml, R&D Systems). From day 8, the medium was changed to stromal cell maturation medium consisting of the basal medium supplemented with PDGF-BB (10 ng/ml). After the stromal cell induction stage, the medium was changed every 2 days.

**Hematopoietic colony-forming unit (CFU) assay**

HPCs were harvested on day 13 from organoids and then $CD34^+CD43^+$ cells were sorted by a FACSAria II (BD Biosciences) for hematopoietic CFU assays. 1,500 $CD34^+CD43^+$ cells were seeded per 35mm-dish in MethoCult H4434 (STEMCELL Technologies) methylcellulose-based medium with recombinant cytokines for CFU assay according to the manufacturer's protocol, and colonies were identified and counted after 14 days of culture.

**Erythroid cell differentiation from organoid-derived HPCs**

Erythroid cell differentiation from organoid-derived HPCs was performed as described in our previous publications with a slight modification (Fujita et al., 2016; Haro-Mora et al., 2020). In brief, $CD34^+$ cells isolated from organoids on day 13 were enriched by magnetic-activated cell sorting (MACS) (Miltenyi Biotec) and were cultured on irradiated OP9 feeder cells for 2 days in Iscove's Modified Dulbecco's Medium (IMDM) (Sigma-Aldrich) containing VEGF (20 ng/ml, Peprotech), SCF (50 ng/ml), Flt3-L (50 ng/ml), TPO (50 ng/ml), IL3 (5 mg/ml), BMP4 (10 ng/ml), erythropoietin (EPO, 5 U/mL, AMGEN), and 15% FBS. Then, the suspension cells were collected and transferred onto fresh irradiated OP9 dishes for 5 days in IMDM medium containing SCF (10 ng/ml), IL3 (1 ng/ml), EPO (2 U/ml), dexamethasone (1 mM, VETone), estradiol (1 mM, Pfizer), and 20% FBS. For erythroid maturation, the medium was replaced with IMDM media containing EPO (2 U/ml), insulin (10 ng/ml, Lilly), transferrin (0.56 mg/ml, Sigma-Aldrich), 2% bovine serum albumin (BSA) (Roche), 2mM L-glutamine, and 20% FBS. The cells were cultured in the maturation medium for 8 days. hiPSC-derived erythroid cells on day 15 of differentiation were harvested for flow cytometry analysis and evaluation of globin-expression patterns by quantitative PCR assay and RP-HPLC (Fujita et al., 2016).

**Macrophage differentiation from organoid-derived HPCs**

Macrophage differentiation from organoid-derived HPCs was performed according to our previous report with minor modifications (Merling et al., 2015). In brief, $CD34^+$ cells isolated from organoids on day 13 were enriched by MACS and cultured for 14 days in macrophage differentiation medium consisting of IMDM medium containing 10% FBS and macrophage colony stimulating factor (M-CSF, 100 ng/ml, Peprotech). Medium was changed two times per week, with suspended cells centrifuged and re-plated until the majority of the cells became adherent. Macrophages were cultured with interferon γ (IFN-γ, 65 U/ml, Actimmune) for 3 days prior to functional assays. Cells were analyzed by cytospin Giemsa stain for macrophage morphology, and color images of stained macrophages were acquired using an EVOS XL Core system (Thermo Fisher Scientific); whole image adjustments of brightness, color balance, and contrast were performed using Adobe Photoshop software (Adobe) without additional image processing. Phagocytosis ability of hiPSC-derived macrophages was evaluated as previously described (Brault et al., 2019). Briefly, macrophages were incubated for 30 minutes at 37°C while shaking (300 rpm) with zymosan A BioParticles isolated from Saccharomyces cerevisiae (Thermo Fisher Scientific) at a multiplicity of infection (MOI) of 10:1. The suspension was then transferred in a FACS tube on ice and the same volume of 2 mg/ml trypan blue was added to stop the reaction and quench the fluorescence of membrane-bound not-internalized particles. Flow cytometry analysis was performed by FACSCanto flow cytometer (BD Biosciences). Dihydrorhodamine-123 (DHR) assay of reactive oxygen species production was performed as previously described (Merling et al., 2015). Briefly, cells were incubated at 37°C for 5 minutes in 400 µL of HBSS containing 130 µM of DHR (Molecular Probes, Thermo Fisher Scientific) and 500 U of catalase (Sigma-Aldrich). Cells were then stimulated for reactive oxygen species production by addition of 100 µL of 400 ng/mL phorbol 12-myristate 13-acetate (PMA, Sigma-Aldrich) in HBSS containing calcium and magnesium (Thermo Fisher Scientific), and incubated at 37°C for 14 minutes, after which flow cytometry was performed using a FACSCalibur system (BD Biosciences).

**T cell differentiation from organoid-derived HPCs**

T cell differentiation from organoid-derived HPCs was described in our previous reports (Maeda et al., 2016; Vizcardo et al., 2013). In brief, organoids were dissociated, and single cells were transferred onto OP9/DLL1 dishes with T cell differentiation medium consisting of OP9 medium supplemented with interleukin 7 (IL7, 5 ng/mL, R&D Systems), Flt3-L (5 ng/mL), and SCF (10 ng/mL). After 3 days, semi adherent cells were transferred to new OP9/DLL1 dishes, followed passaging semi-adherent onto new OP9/DLL1 dishes every 5-7 days. For cytokine release analysis of Mart1-iPSC-derived T cells, we harvested floating cells from OP9/DLL1 dishes on day 22 of the T cell differentiation and enriched $CD4^+CD8^+$ double positive T cells with $CD4^+$ MACS (Miltenyi Biotec) (Maeda et al., 2016). $CD4^+CD8^+$ double positive T cells were stimulated by Mart-1 peptide pulsed T2 cells to be induced to $CD8^+$ single positive T cells. After one week of post stimulation, Mart-1-iPSC-derived $CD8^+$ single positive T cells were co-cultured with Mart-1 peptide or DMSO pulsed T2 cells for cytokines release assay. TCR-Vβ deep sequencing was performed by ImmunoSEQ (Adaptive Biotechnologies) on genomic DNA isolated from NCRM5-EGFP-derived ($GFP^+$) $CD4^+CD8^+CD3^+$ cells on day 32 of T cell differentiation.

**Flow cytometry Analysis**

To analyze cell surface markers, cells were stained with antibodies against cell surface antigens and Propidium Iodide (PI) in FACS buffer (PBS with 2% FBS and 0.1% sodium azide). To determine intracellular cytokine expression, Mart1-iPSC-derived $CD8^+$ single positive cells were co-cultured with Mart-1 peptide or DMSO pulsed T2 cells for 6 hours in the presence of GolgiStop (BD Cytofix/Cytoperm). After stimulation, cells were stained for surface markers and a fixable live/dead stain (Invitrogen). Cells stained for viability and cell surface markers were fixed and permeabilized with Fixation/Permeabilization Solution Kit (BD Cytofix/Cytoperm), and then they were stained with intracellular anti-cytokine antibodies. Stained cells were washed three times by the wash buffer and analyzed by a LSRFortessa (BD Biosciences). All flow cytometry data was analyzed using FlowJo 10.6.1 software (TreeStar).

**Histology**

Yolk sac-like organoids were fixed in 4% paraformaldehyde for 15 mins at room temperature. The fixed organoids were paraffin embedded and sliced into 20 μm thick sections for hematoxylin and eosin (H&E) staining (performed in Histoserv) or immunostaining.

**Immunostaining and confocal imaging**

For analysis of AFP and PDGFRβ, we used whole mount immunostaining. Organoids harvested on day 13 were permeabilized using PBS supplemented with 0.5% Triton X-100 for 15 minutes at room temperature. Blocking was performed for 1 hour at room temperature using a blocking buffer (PBS supplemented with 1% BSA and 0.1% Triton X-100). Subsequently, samples were incubated with primary antibodies overnight at 4°C. Samples were then washed three times with PBS and followed by 1-hour incubation with secondary antibodies at room temperature. Samples were then washed three times with PBS. Confocal images were collected with a 20x plan-apochromat (N.A. 0.75) objective lens using a Nikon Ti2 microscope (Nikon) equipped with a Yokogawa CSU-W1 spinning disk confocal unit and Photometrics BSI sCMOS camera. For analysis of CD34 and CD43, we used section immunostaining. The sections were deparaffinized with xylene followed by different concentrations of ethanol. The immunofluorescence staining procedures were performed according to our previous publication (Isonaka et al., 2018). In brief, the sections were treated with glycine and then with 1% BSA and 10% normal donkey serum in PBS. Afterward, slides were incubated with the primary antibody overnight at 4 °C. On the next day, slides were thoroughly washed in PBS, incubated with secondary antibodies for 1 hour, and then thoroughly washed again. Coverslips were mounted on slides with ProLong Gold antifade reagent (Thermo Scientific), and confocal images were captured using a Zeiss LSM 880 confocal laser scanning microscope (Carl Zeiss).

**RNA sequencing and analysis**

RNA-seq was performed on day 0 iPSC (NCRM5-EGFP) and NCRM5-EGFP derived cells isolated from organoids on day 6 and day 13 with three biological replicates for each time point. After dissociation of organoids, $GFP^{+}$ cells were sorted by a Sony FX500 (Sony) prior to total RNA extraction using the RNeasy Plus Mini Kit (Qiagen). Libraries were pooled and sequenced in two lanes of a HiSeq4000 (Illumina). Raw reads were trimmed for adapters and low-quality bases using Cutadapt 1.18 (Martin, 2011) (-j 8 -b file:adapters.fa -B file:adapters.fa --trim-n -m 20 -o trimmed_R1.fq -p trimmed_R2.fq input_R1.fq input_R2.fq) prior to alignment to the reference genome (Human genome – hg38, annotation Gencode_v24) using STAR 2.6.1 (Dobin et al., 2013). Gene expression quantification analysis was performed using RSEM 1.2.31 (Li and Dewey, 2011). Differential gene expression analysis was performed using edgeR 3.28.1 (Robinson et al., 2010). The analyses are available as a supplementary file (Supplemental analysis 01) and in the accompanying GitHub repository.

**Gene set enrichment analysis**

Gene Set Enrichment Analysis was performed using the GSEAPreranked module within GSEA 4.0.3 (Mootha et al., 2003; Subramanian et al., 2005). Enrichment was tested using a custom yolk sac gene set which comprised the top 200 yolk sac genes (sorted by RPKM) as identified by Cindrova-Davies et al. (2017, supplemental data set S1). 196 of these genes were expressed in our data set and were considered in the enrichment analysis (Table S1).

**Single-cell RNA-seq library preparation and sequencing**

Day 6 and day 13 organoids were dissociated into single cells (see above) and GFP$^{+}$ cells were sorted by a Sony FX500. Cell suspensions from each day were labeled using oligonucleotide-coupled antibodies (Biolegend, TotalSeq-A0251 (barcode: GTCAACTCTTTAGCG) and TotalSeq-A0252 (barcode: TGATGGCCTATTGGG)) to allow for sample multiplexing on one capture lane and cells were mixed at 1:1 ratio. The sample concentration and viability were assessed using the LunaFL fluorescent cell counter (Logos Biosystems). A total recovery of 12,000 cells was targeted. Cell loading, library preparation and quality control were performed as described in the 10x Genomics 3' user guide (v3.0). Three sequencing runs were performed on an Illumina NextSeq 500 (28 X 8 X 0 X 98 base pair (bp) read configuration). The standard 10x

Genomics cellranger pipeline (version 3.1.0) was used to convert raw sequencing data to Fastq format. scRNA-seq reads were aligned to the GRCh38 human reference genome and gene expression counts and feature barcode counts were determined using Cell Ranger.

**Single-cell data analysis**

We used the R package Seurat for cluster analyses and exploration of the data set (Butler et al., 2018; Stuart et al., 2019). The complete analysis is available as a supplemental file (Supplemental analysis 02) and in the accompanying GitHub repository as R markdown documents and knitted PDFs. In the analyses we considered cells with nFeature_RNA (genes) >1000 and <8000 and nCount_RNA (Unique molecular identifiers (UMI)) >2000 and <80000. Cells with mitochondrial read counts > 20% were excluded. This filtering left 4895 day 6 cells and 4401 day 13 cells for analyses with a mean gene count of 5292 and a mean UMI count of 30,069. The data were normalized and scaled, and genes that varied more than expected for their expression level were identified. Principal components (PCs) were calculated on these variable genes which were then used in graph-based clustering followed by UMAP dimensionality reduction. Principal components (PC) 1:30 were considered in the clustering of cells from both days. Principal components 1:21 or 1:25 were used in separate clusterings of D6 cells or D13 cells respectively. The cluster annotations from both the day 6 and the day 13 clusterings were mapped to the resulting UMAP after cells from both days were clustered together. Sub-cluster analysis for the subset of hematopoietic cells considered PCs 1:40. Biomarkers for the day 6, day 13 and the hematopoietic cell clusterings were identified using Seurat's FindAllMarkers (Stuart et al., 2019). We annotated cluster identities using previously characterized genes.

**Statistical analysis**

All data with error bars are presented as mean SD. Difference was assessed using unpaired t-test using GraphPad Prism 9.1.0 (GraphPad software). Values of $P < 0.05$ were considered significant.

**Data availability**

The raw and processed data reported in this publication are archived at NCBI GEO (accession number GSE157140). Analysis code is available as supplemental files (Supplemental analysis 01 and 02) in an accompanying GitHub repository.

**(Supplemental references)**

RSEM accurate transcript quantification from RNA-Seq data with or without a reference genome.

Brault, J., Vigne, B., and Stasia, M.J. (2019). Ex Vivo Models of Chronic Granulomatous Disease. Methods Mol Biol *1982*, 587-622.

Butler, A., Hoffman, P., Smibert, P., Papalexi, E., and Satija, R. (2018). Integrating single-cell transcriptomic data across different conditions, technologies, and species. Nat Biotechnol *36*, 411-420.

Cindrova-Davies, T., Jauniaux, E., Elliot, M.G., Gong, S., Burton, G.J., and Charnock-Jones, D.S. (2017). RNA-seq reveals conservation of function among the yolk sacs of human, mouse, and chicken. Proc Natl Acad Sci U S A *114*, E4753-E4761.

Dobin, A., Davis, C.A., Schlesinger, F., Drenkow, J., Zaleski, C., Jha, S., Batut, P., Chaisson, M., and Gingeras, T.R. (2013). STAR: ultrafast universal RNA-seq aligner. Bioinformatics *29*, 15-21.

Fujita, A., Uchida, N., Haro-Mora, J.J., Winkler, T., and Tisdale, J. (2016). beta-Globin-Expressing Definitive Erythroid Progenitor Cells Generated from Embryonic and Induced Pluripotent Stem Cell-Derived Sacs. Stem Cells *34*, 1541-1552.

Haro-Mora, J.J., Uchida, N., Demirci, S., Wang, Q., Zou, J., and Tisdale, J.F. (2020). Biallelic correction of sickle cell disease-derived induced pluripotent stem cells (iPSCs) confirmed at the protein level through serum-free iPS-sac/erythroid differentiation. Stem Cells Transl Med *9*, 590-602.

Isonaka, R., Sullivan, P., Jinsmaa, Y., Corrales, A., and Goldstein, D.S. (2018). Spectrum of abnormalities of sympathetic tyrosine hydroxylase and alpha-synuclein in chronic autonomic failure. Clin Auton Res *28*, 223-230.

Kennedy, M., Awong, G., Sturgeon, C.M., Ditadi, A., LaMotte-Mohs, R., Zuniga-Pflucker, J.C., and Keller, G. (2012). T lymphocyte potential marks the emergence of definitive hematopoietic progenitors in human pluripotent stem cell differentiation cultures. Cell Rep *2*, 1722-1735.

Li, B., and Dewey, C.N. (2011). RSEM: accurate transcript quantification from RNA-Seq data with or without a reference genome. BMC Bioinformatics *12*, 323.

Maeda, T., Nagano, S., Ichise, H., Kataoka, K., Yamada, D., Ogawa, S., Koseki, H., Kitawaki, T., Kadowaki, N., Takaori-Kondo, A., *et al.* (2016). Regeneration of CD8alphabeta T Cells from T-cell-Derived iPSC Imparts Potent Tumor Antigen-Specific Cytotoxicity. Cancer Res *76*, 6839-6850.

Martin, M. (2011). Cutadapt removes adapter sequences from high-throughput sequencing reads. 2011 *17*, 3.

Merling, R.K., Sweeney, C.L., Chu, J., Bodansky, A., Choi, U., Priel, D.L., Kuhns, D.B., Wang, H., Vasilevsky, S., De Ravin, S.S., *et al.* (2015). An AAVS1-targeted minigene platform for correction of iPSCs from all five types of chronic granulomatous disease. Mol Ther *23*, 147-157.

Mootha, V.K., Lindgren, C.M., Eriksson, K.F., Subramanian, A., Sihag, S., Lehar, J., Puigserver, P., Carlsson, E., Ridderstråle, M., Laurila, E., *et al.* (2003). PGC-1alpha-responsive genes involved in oxidative phosphorylation are coordinately downregulated in human diabetes. Nat Genet *34*, 267-273.

Robinson, M.D., McCarthy, D.J., and Smyth, G.K. (2010). edgeR: a Bioconductor package for differential expression analysis of digital gene expression data. Bioinformatics *26*, 139-140.

Sabatino, M., Ren, J., David-Ocampo, V., England, L., McGann, M., Tran, M., Kuznetsov, S.A., Khuu, H., Balakumaran, A., Klein, H.G., *et al.* (2012). The establishment of a bank of stored clinical bone marrow stromal cell products. J Transl Med *10*, 23.

Stuart, T., Butler, A., Hoffman, P., Hafemeister, C., Papalexi, E., Mauck, W.M., 3rd, Hao, Y., Stoeckius, M., Smibert, P., and Satija, R. (2019). Comprehensive Integration of Single-Cell Data. Cell *177*, 1888-1902.e1821.

Subramanian, A., Tamayo, P., Mootha, V.K., Mukherjee, S., Ebert, B.L., Gillette, M.A., Paulovich, A., Pomeroy, S.L., Golub, T.R., Lander, E.S., *et al.* (2005). Gene set enrichment analysis: a knowledge-based approach for interpreting genome-wide expression profiles. Proc Natl Acad Sci U S A *102*, 15545-15550.

Takebe, T., Sekine, K., Kimura, M., Yoshizawa, E., Ayano, S., Koido, M., Funayama, S., Nakanishi, N., Hisai, T., Kobayashi, T., *et al.* (2017). Massive and Reproducible Production of Liver Buds Entirely from Human Pluripotent Stem Cells. Cell Rep *21*, 2661-2670.

Uchida, N., Haro-Mora, J.J., Fujita, A., Lee, D.Y., Winkler, T., Hsieh, M.M., and Tisdale, J.F. (2017). Efficient Generation of beta-Globin-Expressing Erythroid Cells Using Stromal Cell-Derived Induced Pluripotent Stem Cells from Patients with Sickle Cell Disease. Stem Cells *35*, 586-596.

Vizcardo, R., Masuda, K., Yamada, D., Ikawa, T., Shimizu, K., Fujii, S., Koseki, H., and Kawamoto, H. (2013). Regeneration of human tumor antigen-specific T cells from iPSCs derived from mature CD8(+) T cells. Cell Stem Cell *12*, 31-36.

### Supplementary Figure 1

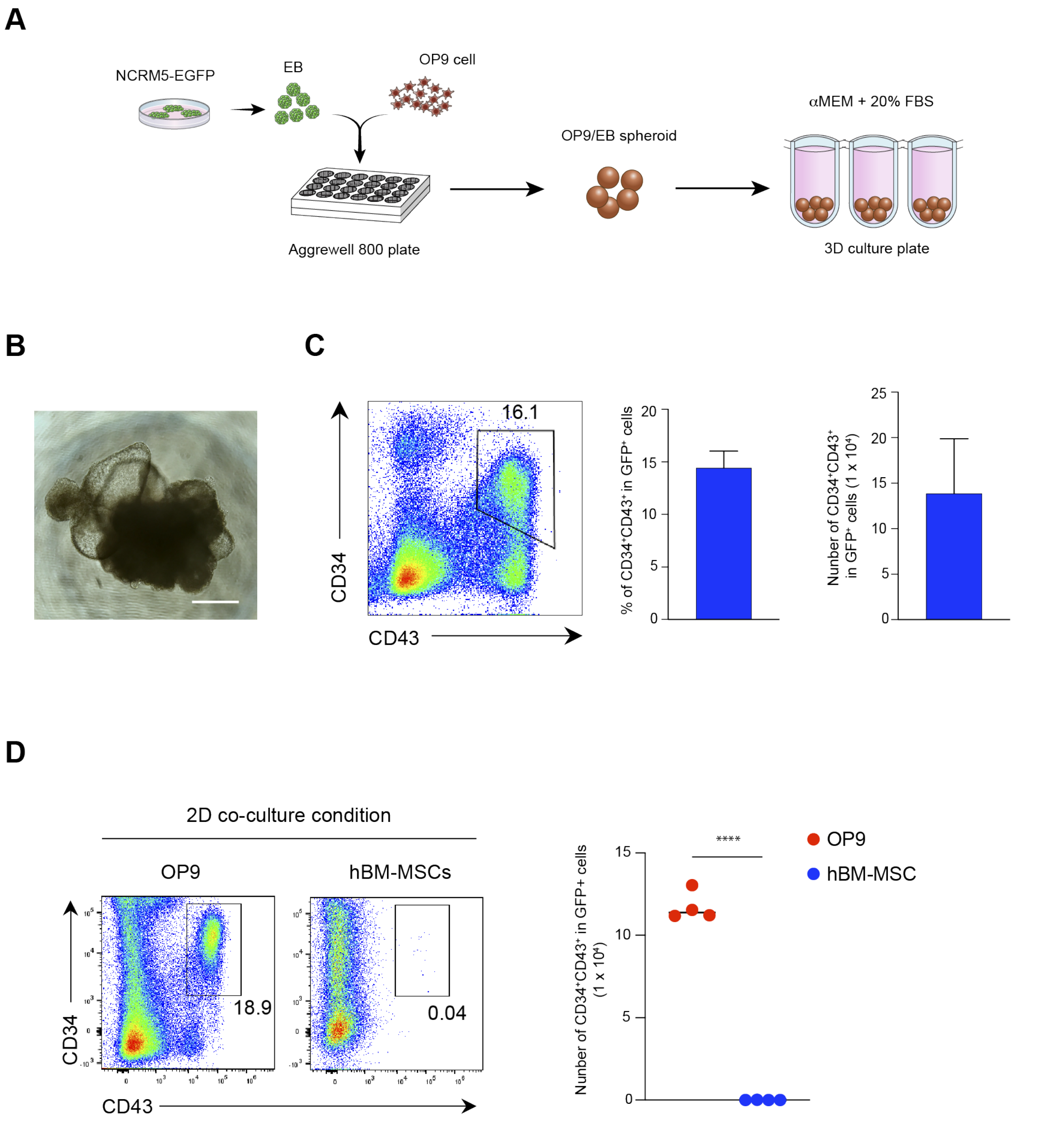

**Figure S1. OP9/EB co-cultured spheroids can induce hematopoiesis from hiPSC similar to the classical hiPSC/OP9 co-culture system in 2D.**

(A) Schematic of the OP9/EB co-cultured spheroid system. (B) Representative microscopy image of OP9/EB co-cultured spheroids on day 13. Scale bar = 500µm. (C) Representative flow cytometry analysis of $CD34^+$ and $CD43^+$ cells in $GFP^+$ cells isolated from OP9/EB co-cultured spheroids on day 13 (left) and bar graphs showing quantification of flow cytometry. The cell number represents $GFP^+CD34^+CD43^+$ cells generated from 1 x $10^6$ hiPSCs. Values represent mean ± SD (n = 3). (D) Representative flow cytometry analysis of $CD34^+$ and $CD43^+$ cells in $GFP^+$ cells isolated from hiPSCs co-cultured on OP9 cells or on hBM-MSCs in a 2D condition for 13 days (left). Quantifications of flow cytometry analysis are shown in the right graph. The cell number represents $GFP^+CD34^+CD43^+$ cells generated from 1 x $10^6$ hiPSCs. Horizontal bars represent mean value (n = 4, ****p < 0.0001).

Supplementary Figure 2

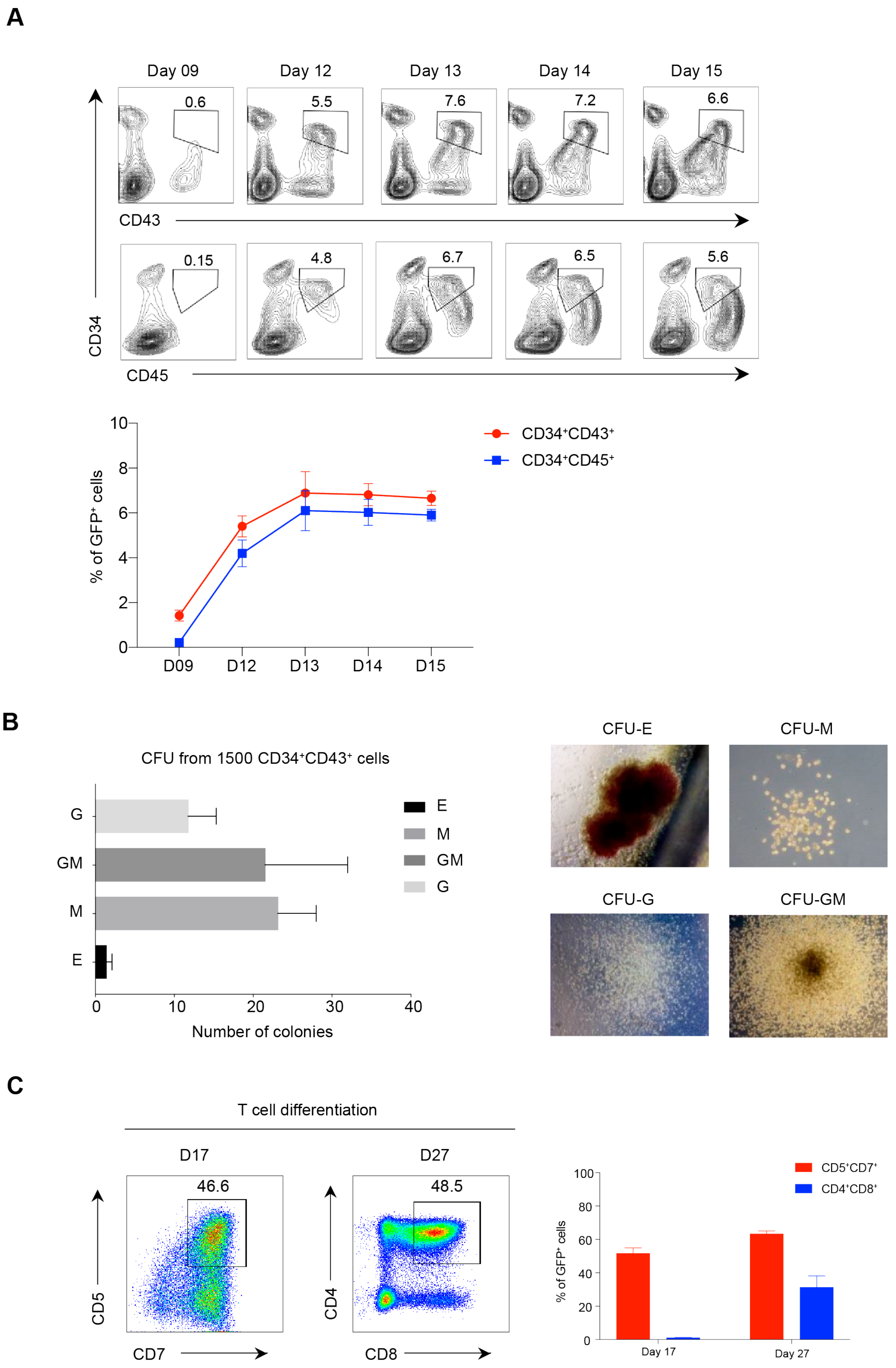

**Figure S2. hiPSC-derived HPCs generated by Hp-spheroids in bioreactors are multipotent.** (A) Kinetic flow cytometry analysis of $CD34^+CD43^+$ and $CD34^+CD45^+$ cells in Hp-spheroids cultured in bioreactors. $GFP^+$ cells are gated for analysis. Representative flow cytometry plots are shown in the upper panel and the quantifications of flow cytometry analysis are shown in the lower graph. Values represent mean ± SD (n = 3). (B) $GFP^+CD34^+CD43^+$ cells were sorted for the hematopoietic CFU assay on day 13. After 14 days of culturing in methylcellulose medium with cytokines, colonies derived from different types of progenitor cells were classified and counted based on morphology. Values represent mean ± SD (n = 3). E; erythroid, M; macrophage, G; granulocyte, GM; granulocyte and macrophage. (C) NCRM5-EGFP derived HPCs can be induced into $CD5^+CD7^+$ T progenitor cells and $CD4^+CD8^+$ T cells on OP9/DLL1 in the presence of cytokines. Representative flow cytometry plots are shown in the left panels. $GFP^+$ cells are gated for analysis. Quantifications of flow cytometry analysis are shown in the right graphs. The percentage of $CD4^+CD8^+$ T cells dramatically increase from day 17 to day 27 of T cell differentiation. Values represent mean ± SD (n = 4).

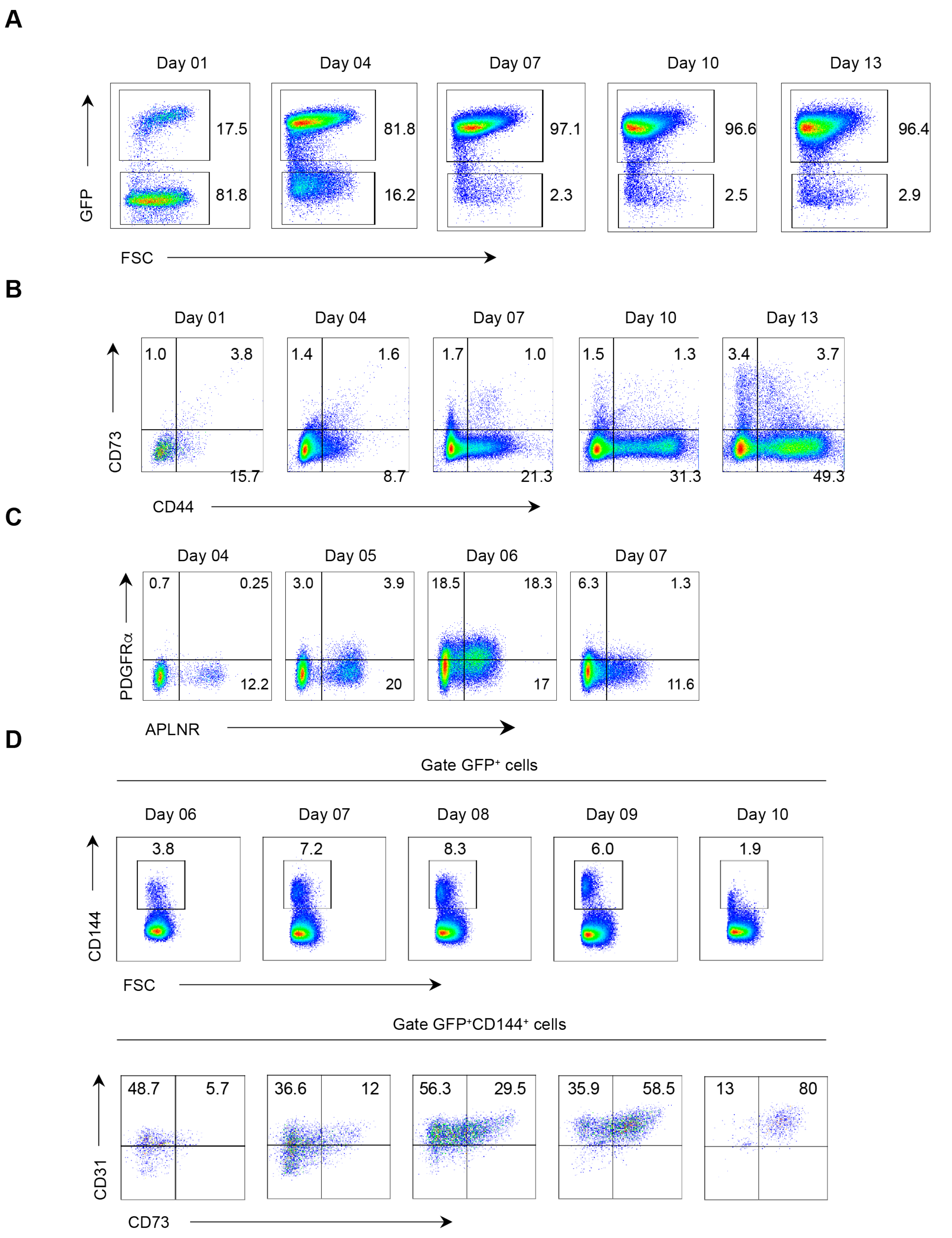
A
Day 01
Day 04
Day 07
Day 10
Day 13
17.5
81.8
81.8
16.2
97.1
2.3
96.6
2.5
96.4
2.9
GFP
FSC
B
Day 01
Day 04
Day 07
Day 10
Day 13
1.0
3.8
15.7
1.4
1.6
8.7
1.7
1.0
21.3
1.5
1.3
31.3
3.4
3.7
49.3
CD73
CD44
C
Day 04
Day 05
Day 06
Day 07
0.7
0.25
12.2
3.0
3.9
20
18.5
18.3
17
6.3
1.3
11.6
PDGFRα
APLNR
D
Gate GFP⁺ cells
Day 06
Day 07
Day 08
Day 09
Day 10
3.8
7.2
8.3
6.0
1.9
CD144
FSC
Gate GFP⁺CD144⁺ cells
48.7
5.7
36.6
12
56.3
29.5
35.9
58.5
13
80
CD31
CD73

**Figure S3. Hematopoietic development is recapitulated during Hp-spheroid development.** (A) Kinetic flow cytometric analysis of GFP$^{+}$ and GFP$^{-}$ cells in Hp-spheroids during their growth in a bioreactor. (B) Kinetic analysis of flow cytometry for CD44 and CD73 in GFP$^{+}$ cells isolated from Hp-spheroids. (C) Representative flow cytometry analysis of APLNR and PDGFRα in GFP$^{+}$ cells isolated from Hp-spheroids from day 4 to day 7. (D) Representative flow cytometry analysis of CD144, CD31, and CD73 in GFP$^{+}$ cells isolated from Hp-spheroids from day 6 to day 10. GFP$^{+}$CD144$^{+}$ cells are gated to analyze CD31 and CD73.

Supplementary Figure 4

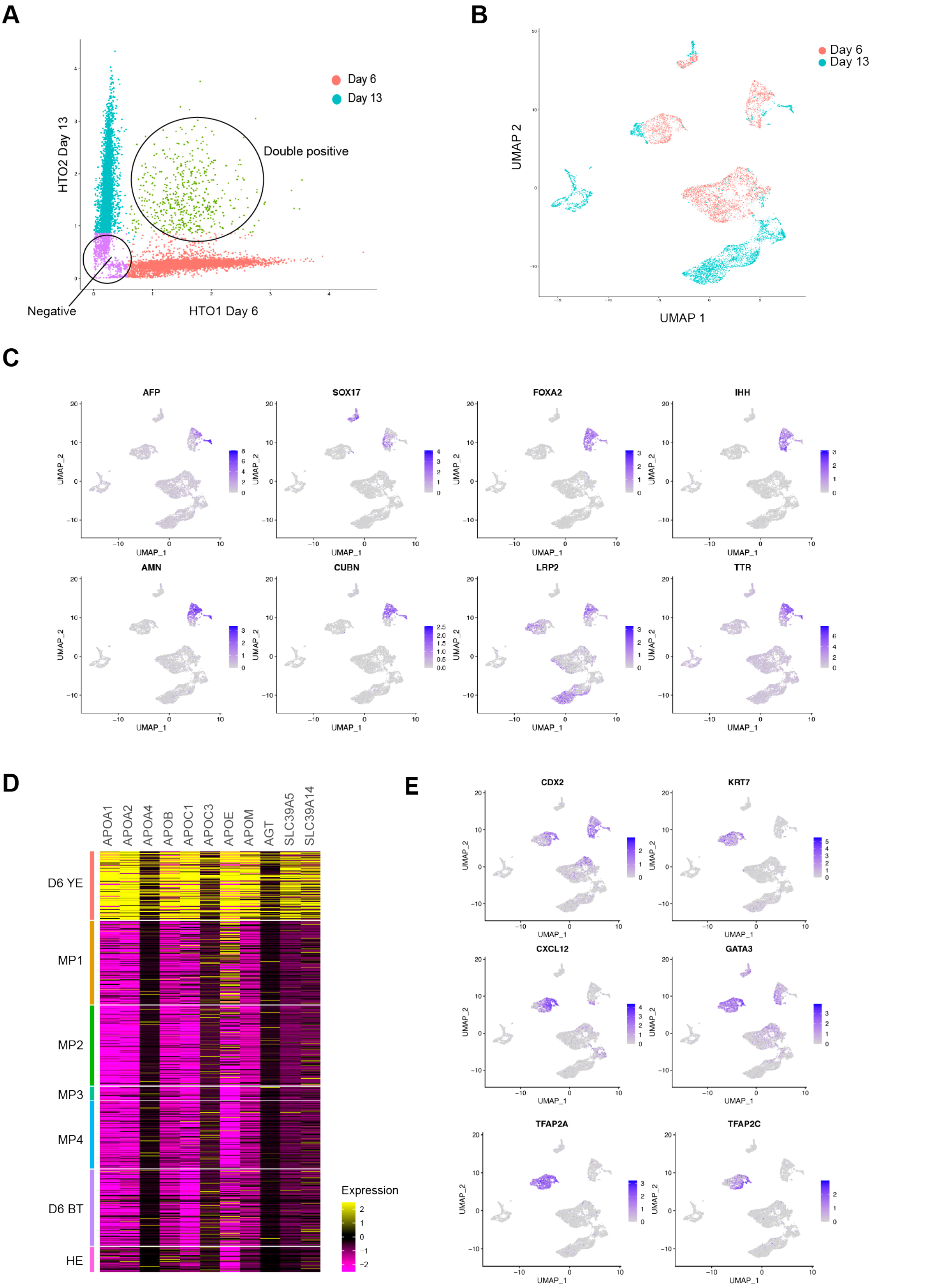

**Figure S4. The scRNA-seq analysis for GFP$^+$ cells isolated from day 6 and day 13 organoids.** (A) Visualization of cells classified as singlets, doublets and negative/ambiguous cells after demultiplexing. (B) UMAP representation for the combined clustering of day 6 (red dots) and day 13 (blue dots) cells with cells labeled by origin. (C) UMAP representation with gene expression for yolk sac endoderm specific genes used in cluster annotations. (D) Heat map showing the expression of genes encoding apolipoproteins and selected SLC transporter genes in day 6 clusters. (E) UMAP representation with gene expression for trophoblast specific genes. The annotated plot is shown in **Figure 3B** (C and E).

Supplementary Figure 5

**A**

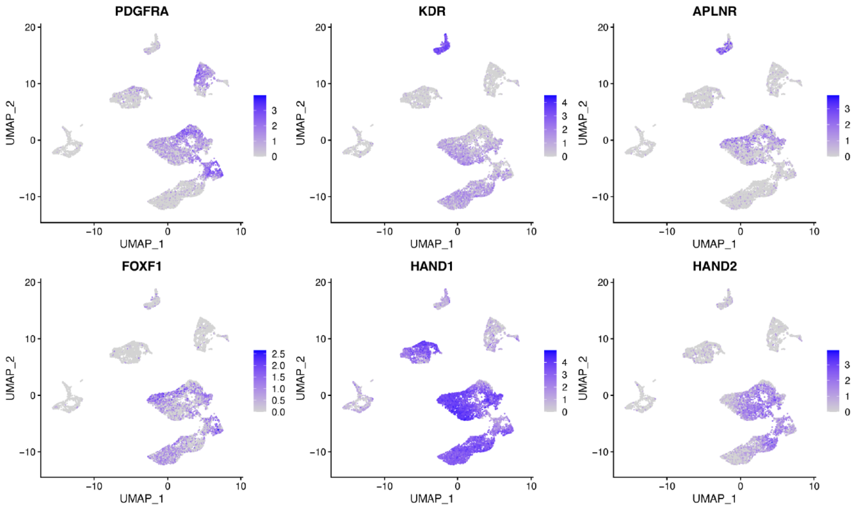


**B**

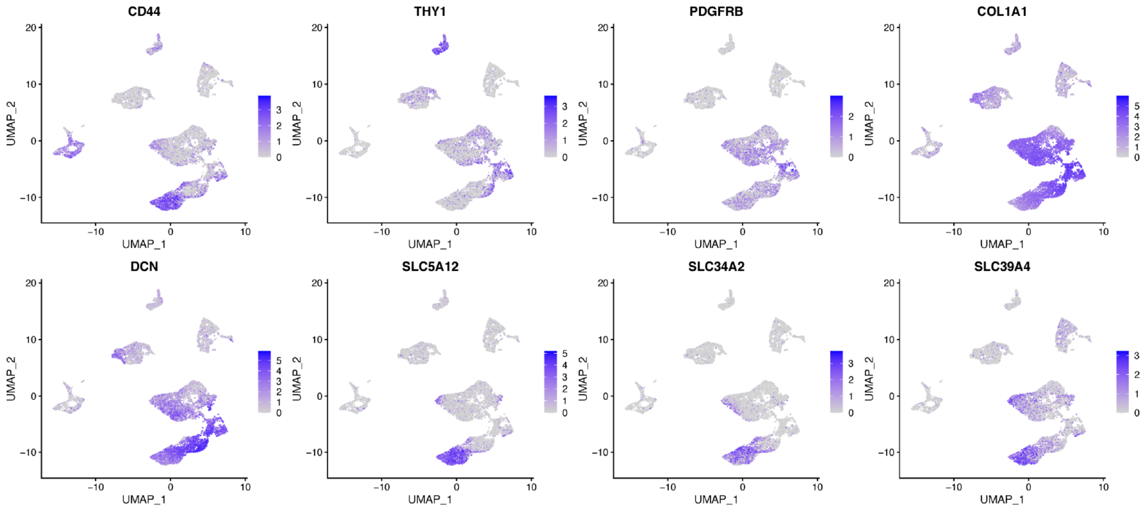


**C**

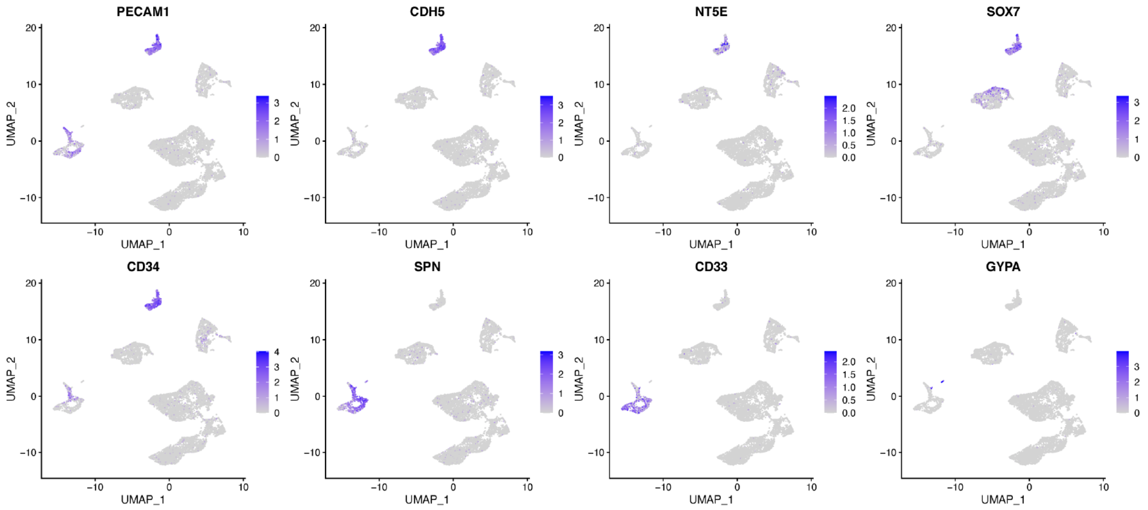

**Figure S5. Gene expression plots of genes specific to mesodermal progenitor cells and their derivatives.**

(A-C) UMAP representation with gene expression visualized for mesoderm progenitor (A), stromal/mesenchymal (B), and hemato-vascular (C) specific genes. The annotated plot is shown in **Figure 3B**.

Supplementary Figure 6

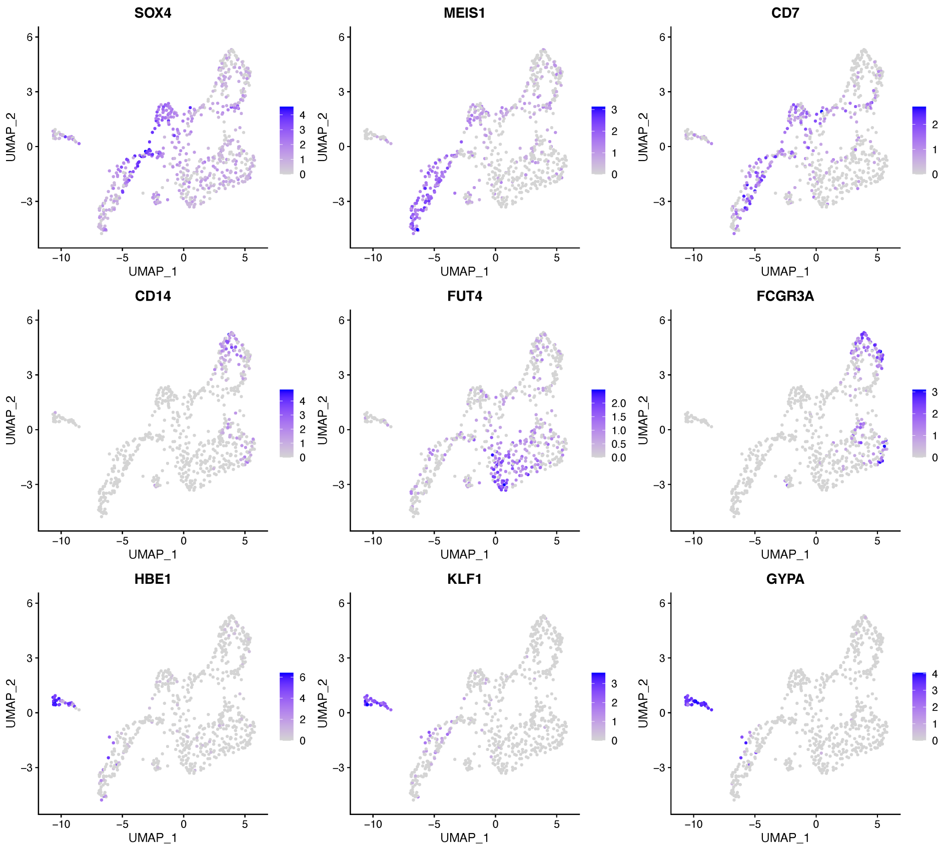

**Figure S6. Hematopoietic lineage gene expression in day 13 organoids.**

UMAP representation for day 13 hematogenic cells with gene expression visualized for various hematopoietic marker genes

Supplementary Figure 7

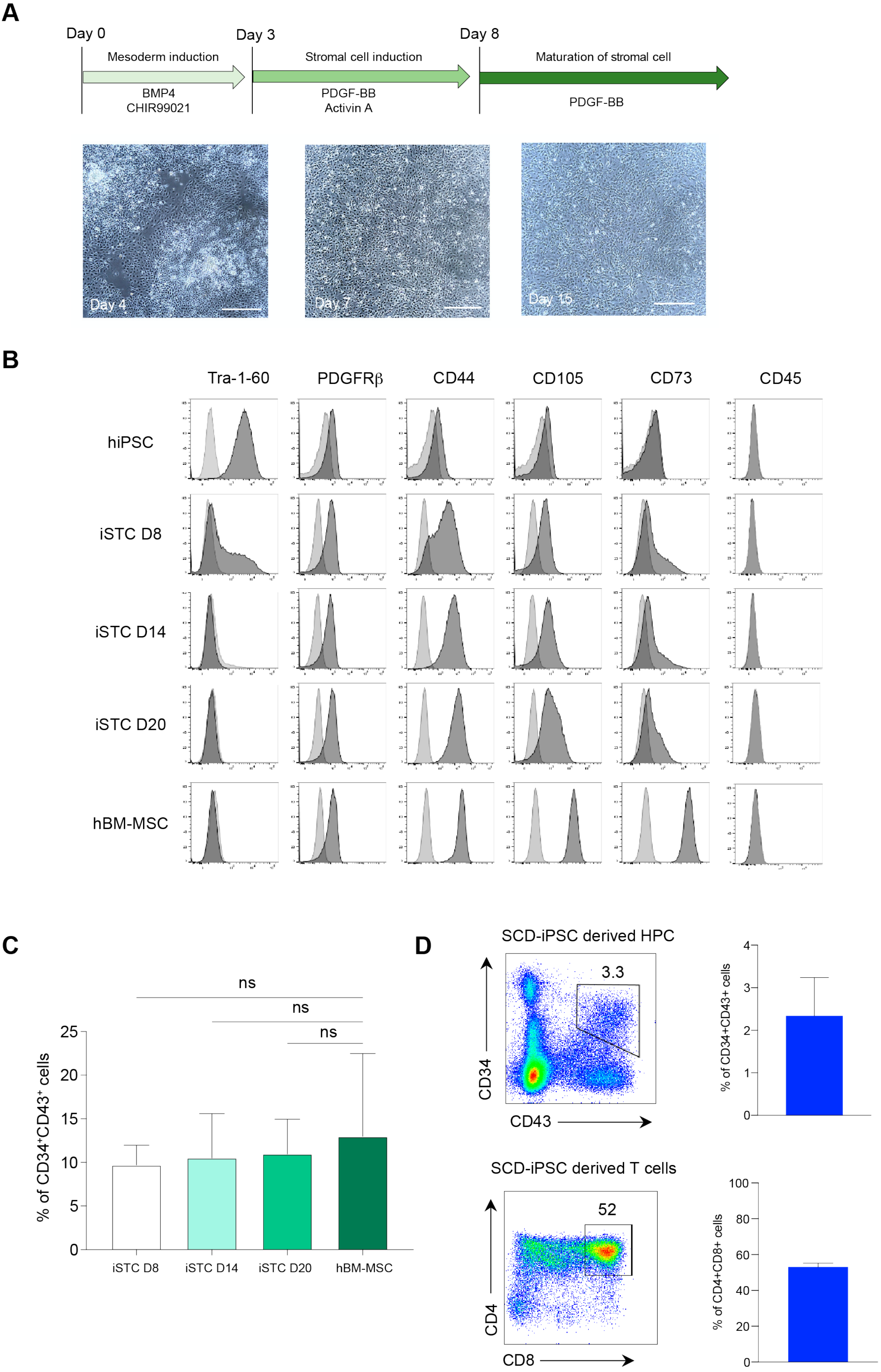

**Figure S7. hiPSC-derived stromal cells (iSTCs) can be used for the yolk-like organoid system.** (A) Schematic of induction of stromal cells from hiPSCs. Lower pictures show the morphologies of iSTCs at different time points of the differentiation. Scale bar = 500μm (B) Flow cytometry analysis of hBM-MSCs and iSTCs isolated from day 8 (D8), day 14 (D14), and day 20 (D20) of differentiation. During differentiation, Tra-1-60$^+$ cells (a pluripotent marker) decrease and typical stromal cell markers start to express, including PDGFRβ, CD44, CD105, and CD73. CD45 is a negative marker for hBM-MSCs and iSTCs. (C) NCRM5-EGFP were co-cultured with hBM-MSCs or iSTCs isolated from D8, D14, and D20 to form Hp-spheroids. These spheroids were cultured in bioreactors for 13 days and the percentage of CD34$^+$CD43$^+$ cells in GFP$^+$ cells isolated from each type of Hp-spheroids was analyzed by flow cytometry. Values represent mean ± SD (n = 3). (D) SCD-iPSC cells can generate CD34$^+$CD43$^+$ cells by the iSTC-organoid system for 13 days. SCD-iPSC derived HPCs can be induced to CD4$^+$CD8$^+$ T lineage cells by OP9/DLL1 co-culture system for 27 days. Representative flow cytometry plots are shown in the left and quantifications of flow cytometry analysis are shown in the right graphs. Values represent mean ± SD (n = 3).
