## Supplementary material for "Self-organized yolk sac-like organoids allow for scalable generation of multipotent hematopoietic progenitor cells from human induced pluripotent stem cells": Sup analysis1

### Tamaoki et al. - Bulk RNAseq analysis

2021-03-15 22:25:29

The document contains a differential gene expression analysis for bulk RNAseq data for iPSC (NCRM5) controls and NCRM5-EGFP derived cells isolated from hi-spheroids on day 6 (D06) and day 13 (D13). The following contrasts were made: iPSC vs D06, D06 vs D13 and iPSC vs D13. Ranked gene lists were created for gene set enrichment analysis (GSEA). To replicate these analyses RSEM counts (file bulk_count_mtx.txt) can be downloaded from GEO (accession GSE157140).

```
library(edgeR)
library(RColorBrewer)
library(gplots)
library(knitr)
opts_chunk$set(tidy.opts=list(width.cutoff=60),tidy=TRUE)
```

```
# load raw counts
c <- read.table("../bulk/RawCountFile_rsemgenes.txt", header = TRUE)

# original column names 'gene_id' '1_D6_7_23' '2_D13_7_23'
# '3_Nl5_GFP_1' '4_Nl5_GFP_2' '5_Nl5_GFP_3' '6_D6_2' '7_D6_3'
# '8_D13_2' '9_D13_3' samples 1_D6_7_23 and 2_D13_7_23 were
# treated as replicate 1

# simplified naming sheme
colnames(c) <- c("id", "D6_1", "D13_1", "NCRM5_1", "NCRM5_2",
    "NCRM5_3", "D6_2", "D6_3", "D13_2", "D13_3")

# change colum order
cd <- c[, c(1, 4, 5, 6, 2, 7, 8, 3, 9, 10)]

write.csv(cd, "RawCountFile_rsemgenes_reordered.csv")

# for GEO
write.table(cd, "bulk_count_mtx.txt", sep = "\t", row.names = F)
# read GEO file, GEO accession number GSE157140
cd <- read.table("bulk_count_mtx.txt", sep = "\t", header = T)

# group assignement
Group <- factor(c("NCRM5", "NCRM5", "NCRM5", "D6", "D6", "D6",
    "D13", "D13", "D13"))

# create experimental design data frame defining treatment
# type for each sample
df <- data.frame(Sample = colnames(cd[, c(2:10)]), Group)

# generate DGEList object containing raw counts, the
# treatment type for each column, and the gene annotations
y <- DGEList(counts = cd[, c(2:10)], group = df$Group, genes = cd[,
    1])
```

```r
# dim(y) 60554 genes

# calculate library size for each sample and store within
# DGElist
y$samples$lib.size <- colSums(y$counts)

# multiple genes did receive no or very few counts exclude
# transcripts that do not have at least two samples with more
# than one count per million
keep <- rowSums(cpm(y) > 1) >= 2
y <- y[keep, , keep.lib.sizes = FALSE]

# number of genes with expression data remaining after count
# filtering dim(y) 17605 genes

# calculate normalization factors for each sample
y <- calcNormFactors(y)

# review library sizes and normalization factors
y$samples
```

```
##         group lib.size norm.factors
## NCRM5_1 NCRM5 16462919    1.0470780
## NCRM5_2 NCRM5 26079512    1.0356643
## NCRM5_3 NCRM5 24342832    1.0526793
## D6_1       D6 20610539    0.8965679
## D6_2       D6 27795387    1.0527813
## D6_3       D6 28653055    1.0820409
## D13_1     D13 25762012    0.8656076
## D13_2     D13 30960249    0.9940958
## D13_3     D13 32160396    0.9967623
```

```r
design <- model.matrix(~0 + Group)
colnames(design) <- levels(Group)

# estimate dispersions
y <- estimateDisp(y, design)
```

Genes that do not have at least two treatments with more than one count per million were excluded. This leaves us with 17605 genes with expression data. The function plotMDS produces a plot in which distances between samples correspond to leading biological coefficient of variation (BCV) between samples. The plot indicates good replication.

```r
plotMDS(y, method = "bcv", cex = 0.7)
```

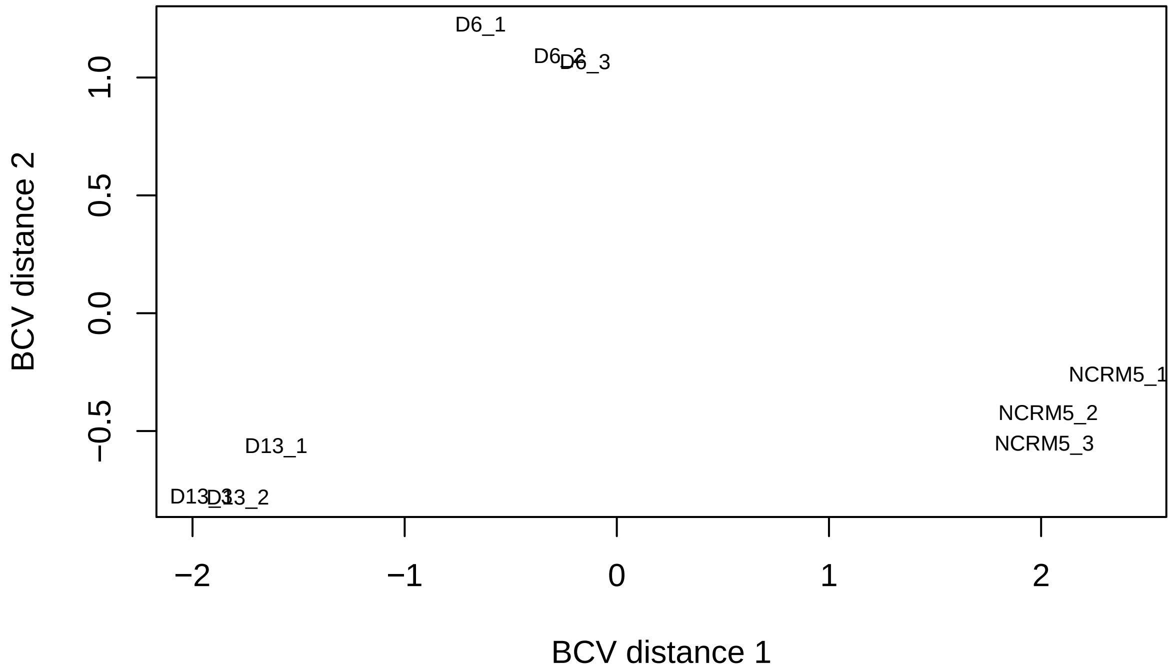


##### Differential gene expression analysis - NCRM5 (iPSC) vs D06

```
contrasts <- makeContrasts(NCRM5vsD06 = NCRM5 - D6, NCRM5vsD13 = NCRM5 -
    D13, D06vsD13 = D6 - D13, levels = design)

# fit linear model
fit <- glmQLFit(y, design)

# identify genes that change expression - NCRM5 vs D06
qlf0_6 <- glmQLFTest(fit, contrast = contrasts[, "NCRM5vsD06"])
```

Top differentially expressed genes. Sorted by p-val. Negative Fold Change indicates upregulation at D06.

```
# top differentially expressed genes
topTags(qlf0_6, n = 20, adjust.method = "BH", sort.by = "PValue",
    p.value = 1)
```

```
## Coefficient:  -1*D6 1*NCRM5
##                            genes      logFC    logCPM         F       PValue
## 5607    ENSG00000124942.13_AHNAK  -6.917159 10.892689 2182.1967 7.659211e-12
## 14100   ENSG00000176438.12_SYNE3  -8.007492  4.426349 1949.0138 1.256250e-11
## 8915    ENSG00000147041.11_SYTL5  -6.484245  6.060363 1536.5954 3.555017e-11
## 4518    ENSG00000116016.13_EPAS1  -7.936412  7.504246 1463.1889 4.403424e-11
## 13106     ENSG00000171564.11_FGB -15.804454  9.043693 1368.9908 5.889877e-11
## 222   ENSG00000008513.14_ST3GAL1  -4.572578  6.939891 1243.5405 8.963125e-11
## 8488      ENSG00000143878.9_RHOB  -3.002674  6.617140 1227.8605 9.473862e-11
## 1635     ENSG00000084674.13_APOB  -8.230298  8.586378 1226.3151 9.526110e-11
## 8505    ENSG00000143995.19_MEIS1 -10.111445  5.909603 1164.4445 1.194220e-10
## 6844    ENSG00000134138.19_MEIS2 -10.449228  5.583449 1150.6195 1.258137e-10
## 11435    ENSG00000164741.14_DLC1  -4.194262  5.993891 1091.0562 1.586638e-10
## 3550     ENSG00000108557.17_RAI1  -5.018838  4.400158 1036.1027 1.987907e-10
## 16254   ENSG00000186340.14_THBS2   5.325065  2.874474 1008.6894 2.234530e-10
```

```
## 1684  ENSG00000086062.12_B4GALT1  -3.582526  6.389787  992.9945 2.392684e-10
## 1695   ENSG00000086475.14_SEPHS1   3.177033  8.317091  945.9299 2.956784e-10
## 15776   ENSG00000184500.14_PROS1  -4.105977  5.432141  943.6249 2.988394e-10
## 11247   ENSG00000164093.15_PITX2 -11.015076  6.907783  942.8526 2.999077e-10
## 6945     ENSG00000134686.16_PHC2  -3.393878  5.533204  914.8946 3.419472e-10
## 9059      ENSG00000148180.17_GSN  -2.971919  7.519266  903.5332 3.610817e-10
## 9001    ENSG00000147601.13_TERF1   3.338833  8.093293  886.3556 3.925754e-10
##                FDR
## 5607  1.105814e-07
## 14100 1.105814e-07
## 8915  1.938057e-07
## 4518  1.938057e-07
## 13106 2.073826e-07
## 222   2.096340e-07
## 8488  2.096340e-07
## 1635  2.096340e-07
## 8505  2.214950e-07
## 6844  2.214950e-07
## 11435 2.539342e-07
## 3550  2.916425e-07
## 16254 3.008800e-07
## 1684  3.008800e-07
## 1695  3.105809e-07
## 15776 3.105809e-07
## 11247 3.105809e-07
## 6945  3.344433e-07
## 9059  3.345707e-07
## 9001  3.455645e-07
```

```r
# set p-value cutoff
p.value = 0.01

# the total number of differentially expressed genes at
# selected adjusted p-value cutoff
de <- decideTestsDGE(qlf0_6, adjust.method = "BH", p.value)
summary(de)
```

```
##        -1*D6 1*NCRM5
## Down            4226
## NotSig          9816
## Up              3563
```

```r
# highlight DGE genes on a plot of logfold-change versus
# log-counts-per-millions
detags <- rownames(qlf0_6)[as.logical(de)]
plotSmear(qlf0_6, de.tags = detags, main = "iPSC (positive FC) vs D06")
abline(h = c(-1, 1), col = "blue")
```

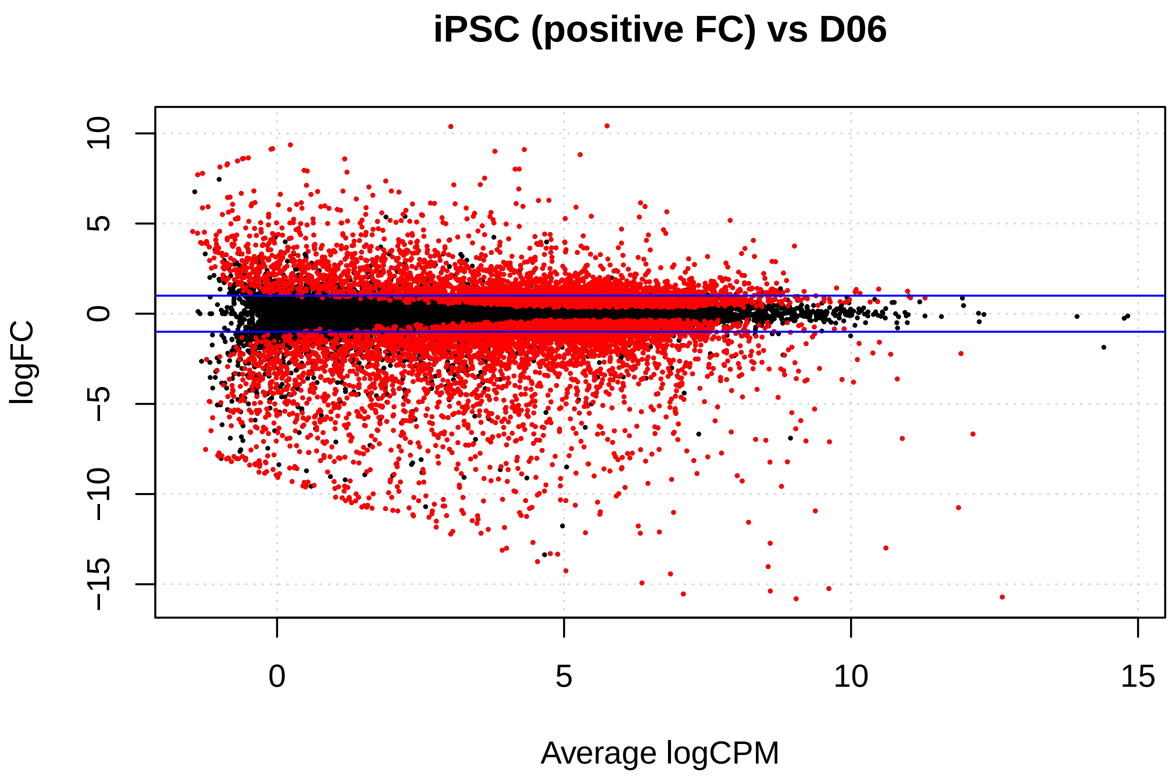


```
# results table
res0_6 <- qlf0_6$table
# restore gene column
res0_6 <- cbind(res0_6, qlf0_6$genes)

# add column of adjusted p values
res0_6 <- cbind(res0_6, Padj = p.adjust(res0_6$PValue, method = "BH"))

# change column order
res0_6 <- res0_6[, c(5, 1:4, 6)]
res0_6 <- res0_6[order(res0_6$Padj), ]

# output results table
write.csv(res0_6, "iPSCvsD06.csv", row.names = FALSE)
```

##### Differential gene expression analysis - D06 vs D13

```
# identify genes that change expression - D06 vs D13
qlf6_13 <- glmQLFTest(fit, contrast = contrasts[, "D06vsD13"])
```

Top differentially expressed genes. Sorted by p-val. Negative Fold Change indicates upregulation at D13.

```
# top differentially expressed genes
topTags(qlf6_13, n = 20, adjust.method = "BH", sort.by = "PValue",
    p.value = 1)
```

```
## Coefficient:  -1*D13 1*D6
##                                  genes      logFC   logCPM        F
## 15575        ENSG00000183688.4_FAM101B  -3.434282 5.250650 719.0722
## 95               ENSG00000005381.7_MPO -13.759924 8.001422 713.3475
```

```
## 17135          ENSG00000196415.9_PRTN3 -12.643251 5.226596 695.1174
## 3306          ENSG00000106404.13_CLDN15  -3.704526 4.223673 565.2144
## 2855            ENSG00000103811.15_CTSH  -2.607032 5.054734 538.9206
## 10602          ENSG00000161270.19_NPHS1  -5.598756 2.210935 535.0982
## 1876              ENSG00000090382.6_LYZ -10.562195 9.364657 510.6176
## 5320             ENSG00000122862.4_SRGN -10.355073 6.876388 424.2697
## 4916           ENSG00000119535.17_CSF3R  -4.705314 3.675219 398.5338
## 1829             ENSG00000089199.9_CHGB  -3.725746 3.968848 394.3350
## 10436          ENSG00000160179.18_ABCG1  -4.378521 4.431233 387.6452
## 57947 ENSG00000278730.1_RP11-147L13.11  -3.350184 5.550657 349.9787
## 8640           ENSG00000144891.17_AGTR1  -6.556318 4.730177 343.9400
## 574            ENSG00000041353.9_RAB27B  -4.913247 4.066607 342.1507
## 20165        ENSG00000205038.11_PKHD1L1  -5.472521 3.810385 322.8502
## 6797            ENSG00000133800.8_LYVE1 -10.317552 2.923338 318.5192
## 10957         ENSG00000163220.10_S100A9 -11.967340 6.506949 306.6978
## 9441           ENSG00000151655.17_ITIH2  -8.857430 1.507921 306.3766
## 5812             ENSG00000126264.9_HCST  -5.538368 1.626093 305.5739
## 8915           ENSG00000147041.11_SYTL5   2.269550 6.060363 299.2294
##             PValue          FDR
## 15575 9.755085e-10 6.632860e-06
## 95    1.010000e-09 6.632860e-06
## 17135 1.130280e-09 6.632860e-06
## 3306  2.774138e-09 1.031911e-05
## 2855  3.410112e-09 1.031911e-05
## 10602 3.516879e-09 1.031911e-05
## 1876  4.307334e-09 1.083294e-05
## 5320  9.591361e-09 2.110699e-05
## 4916  1.256217e-08 2.265364e-05
## 1829  1.314876e-08 2.265364e-05
## 10436 1.415451e-08 2.265364e-05
## 57947 2.197306e-08 3.045152e-05
## 8640  2.367935e-08 3.045152e-05
## 574   2.421592e-08 3.045152e-05
## 20165 3.107025e-08 3.622579e-05
## 6797  3.292318e-08 3.622579e-05
## 10957 3.871716e-08 3.644323e-05
## 9441  3.889136e-08 3.644323e-05
## 5812  3.933095e-08 3.644323e-05
## 8915  4.302870e-08 3.682662e-05
```

```r
# set p-value cutoff
p.value = 0.01

# the total number of differentially expressed genes at
# selected adjusted p-value cutoff
de <- decideTestsDGE(qlf6_13, adjust.method = "BH", p.value)
summary(de)
```

```
##        -1*D13 1*D6
## Down          2408
## NotSig       12758
## Up            2439
```

```r
# highlighted DGE genes on a plot of logfold-change versus
# log-counts-per-millions
```

```
detags <- rownames(qlf6_13)[as.logical(de)]
plotSmear(qlf6_13, de.tags = detags, main = "D06 (positive FC) vs D13")
abline(h = c(-1, 1), col = "blue")
```

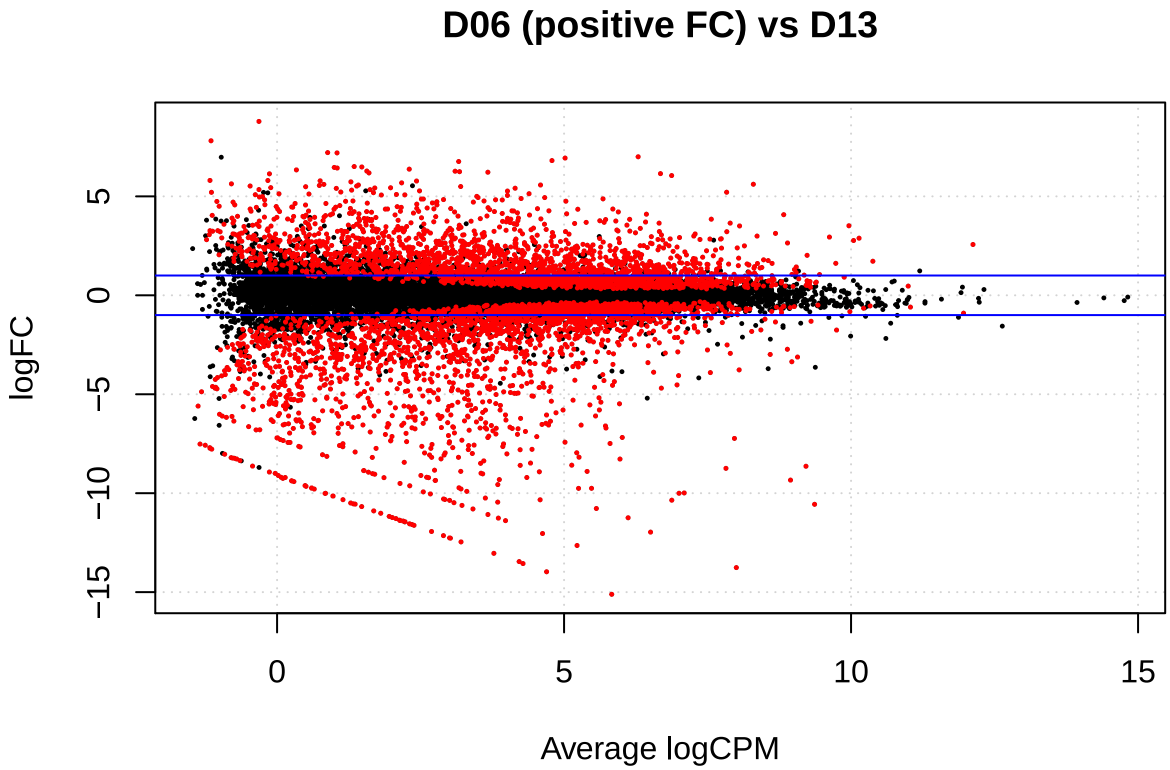


```
# results table
res6_13 <- qlf6_13$table
# restore gene column
res6_13 <- cbind(res6_13, qlf6_13$genes)

# Add column of adjusted p values
res6_13 <- cbind(res6_13, Padj = p.adjust(res6_13$PValue, method = "BH"))

# change colum order
res6_13 <- res6_13[, c(5, 1:4, 6)]
res6_13 <- res6_13[order(res6_13$Padj), ]

# output results table
write.csv(res6_13, "D06vsD13.csv", row.names = FALSE)
```

##### Differential gene expression analysis - iPSC vs D13

```
# identify genes that change expression - iPSC vs D13
qlf0_13 <- glmQLFTest(fit, contrast = contrasts[, "NCRM5vsD13"])
```

Top differentially expressed genes. Sorted by p-val. Negative Fold Change indicates upregulation at D13.

```
# top differentially expressed genes
topTags(qlf0_13, n = 20, adjust.method = "BH", sort.by = "PValue",
    p.value = 1)
```

```
## Coefficient:  -1*D13 1*NCRM5 
##                             genes      logFC    logCPM         F       PValue
## 5607    ENSG00000124942.13_AHNAK  -6.660461 10.892689 2070.3705 9.643306e-12
## 14100   ENSG00000176438.12_SYNE3  -8.126478  4.426349 2006.1028 1.107099e-11
## 222   ENSG00000008513.14_ST3GAL1  -4.952450  6.939891 1411.9595 5.145745e-11
## 9001    ENSG00000147601.13_TERF1   4.208647  8.093293 1297.1686 7.453589e-11
## 1695   ENSG00000086475.14_SEPHS1   3.782844  8.317091 1268.3845 8.221226e-11
## 4518    ENSG00000116016.13_EPAS1  -7.058607  7.504246 1232.9467 9.304367e-11
## 13106     ENSG00000171564.11_FGB -14.648557  9.043693 1200.8432 1.044052e-10
## 1635     ENSG00000084674.13_APOB  -8.052248  8.586378 1189.7740 1.087132e-10
## 38426    ENSG00000245532.5_NEAT1  -3.924231  6.715826 1169.4873 1.171902e-10
## 9756    ENSG00000154277.12_UCHL1   3.602772  7.171129 1084.1387 1.631289e-10
## 47932   ENSG00000261371.5_PECAM1  -8.388355  5.293716 1065.9341 1.756383e-10
## 3550     ENSG00000108557.17_RAI1  -5.085009  4.400158 1060.5419 1.795676e-10
## 6844    ENSG00000134138.19_MEIS2  -9.834066  5.583449 1022.1711 2.108818e-10
## 9059      ENSG00000148180.17_GSN  -3.130600  7.519266  989.6286 2.428372e-10
## 8505    ENSG00000143995.19_MEIS1  -9.245434  5.909603  988.8854 2.436340e-10
## 2528      ENSG00000101439.8_CST3  -4.195555  5.815157  973.1528 2.612793e-10
## 2181    ENSG00000100078.3_PLA2G3   6.794760  3.441192  958.2518 2.794598e-10
## 9622     ENSG00000153071.14_DAB2  -4.813937  6.607766  944.8062 2.972142e-10
## 4817   ENSG00000118495.18_PLAGL1  -6.352237  4.353784  944.5205 2.976062e-10
## 4262     ENSG00000114166.7_KAT2B  -4.702113  3.951360  912.0669 3.465909e-10
##                FDR
## 5607  9.745235e-08
## 14100 9.745235e-08
## 222   2.292371e-07
## 9001  2.292371e-07
## 1695  2.292371e-07
## 4518  2.292371e-07
## 13106 2.292371e-07
## 1635  2.292371e-07
## 38426 2.292371e-07
## 9756  2.634407e-07
## 47932 2.634407e-07
## 3550  2.634407e-07
## 6844  2.757557e-07
## 9059  2.757557e-07
## 8505  2.757557e-07
## 2528  2.757557e-07
## 2181  2.757557e-07
## 9622  2.757557e-07
## 4817  2.757557e-07
## 4262  3.050866e-07
```

```
# set p-value cutoff
p.value = 0.01

# the total number of differentially expressed genes at
# selected adjusted p-value cutoff
de <- decideTestsDGE(qlf0_13, adjust.method = "BH", p.value)
summary(de)
```

```
##        -1*D13 1*NCRM5
## Down              5171
## NotSig            8147
```

```
## Up                   4287
```

```r
# highlighted DGE genes on a plot of logfold-change versus
# log-counts-per-millions
detags <- rownames(qlf0_13)[as.logical(de)]
plotSmear(qlf0_13, de.tags = detags, main = "iPSC (positive FC) vs D13")
abline(h = c(-1, 1), col = "blue")
```

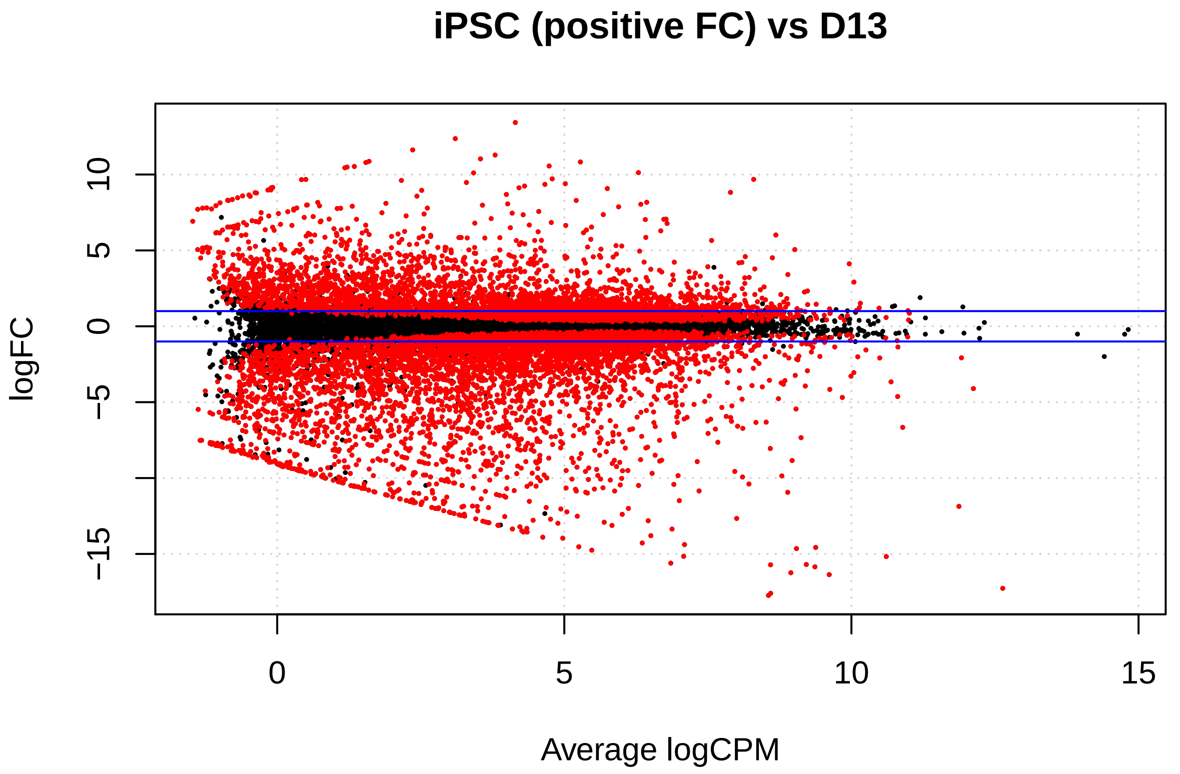


```r
# results table
res0_13 <- qlf0_13$table
# restore gene column
res0_13 <- cbind(res0_13, qlf0_13$genes)

# Add column of adjusted p values
res0_13 <- cbind(res0_13, Padj = p.adjust(res0_13$PValue, method = "BH"))

# change colum order
res0_13 <- res0_13[, c(5, 1:4, 6)]
res0_13 <- res0_13[order(res0_13$Padj), ]

# output results table
write.csv(res0_13, "iPSCvsD13.csv", row.names = FALSE)
```

##### Heatmap for variable genes (log2 counts per million)

```r
# get log2 counts per million
logcounts <- cpm(y, log = TRUE)

# rownames(logcounts) <- y$genes$genes
rownames(logcounts) <- gsub("ENSG.+?\\_", "", y$genes$genes)
```

```r
# estimate the variance for each row in the logcounts matrix
var_genes <- apply(logcounts, 1, var)
# head(var_genes)

# get the gene names for the top 500 most variable genes
select_var <- names(sort(var_genes, decreasing = TRUE))[1:500]

# subset logcounts matrix
highly_variable_lcpm <- logcounts[select_var, ]

## Get some nicer colours
mypalette <- brewer.pal(11, "RdYlBu")
morecols <- colorRampPalette(mypalette)

# plot the heatmap
heatmap.2(highly_variable_lcpm, col = rev(morecols(50)), trace = "none",
    main = "Top 500 most variable genes", scale = "row", labRow = FALSE)
```

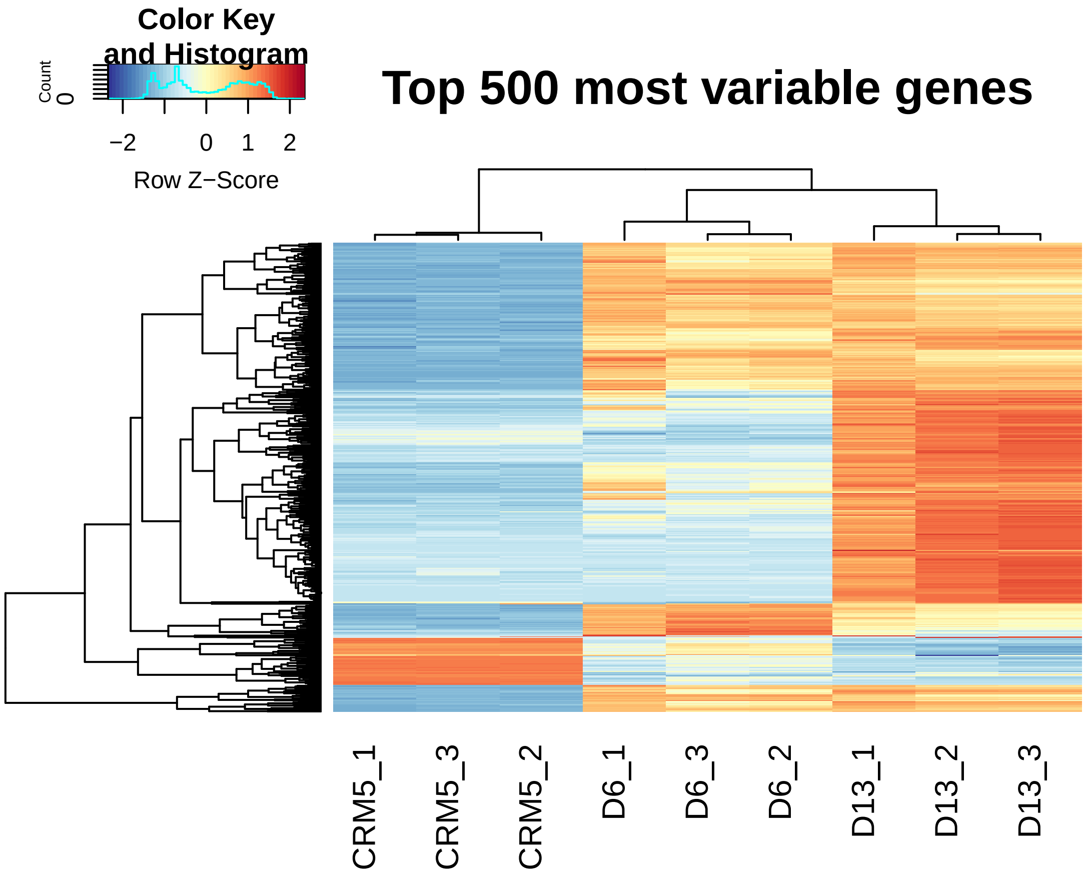


**Generate ranked gene lists for gene set enrichment analysis (GSEA)**

```r
# D6 to the left in GSEA
t <- read.csv("iPSCvsD06.csv", header = T)
# adjust gene names to work with GSEA
t$genes <- gsub("ENSG.+?\\_", "", t$genes)
```

```r
# keep sign
t$rank <- t$logFC * log10(t$PValue)
rnk <- t[c(1, 7)]

write.table(rnk, "iPSC_D06_rank.rnk", row.names = F, col.names = F,
    quote = F, sep = "\t")

# D13 to the left in GSEA
t <- read.csv("iPSCvsD13.csv", header = T)
# adjust gene names to work with GSEA
t$genes <- gsub("ENSG.+?\\_", "", t$genes)

t$rank <- t$logFC * log10(t$PValue)

rnk <- t[c(1, 7)]

write.table(rnk, "iPSCvsD13_rank.rnk", row.names = F, col.names = F,
    quote = F, sep = "\t")

# D06 to the left in GSEA
t <- read.csv("D06vsD13.csv", header = T)
# adjust gene names to work with GSEA
t$genes <- gsub("ENSG.+?\\_", "", t$genes)

t$rank <- t$logFC * -(log10(t$PValue))

rnk <- t[c(1, 7)]

write.table(rnk, "D06vsD13_rank.rnk", row.names = F, col.names = F,
    quote = F, sep = "\t")
```

```r
sessionInfo()
```

```
## R version 3.6.3 (2020-02-29)
## Platform: x86_64-apple-darwin15.6.0 (64-bit)
## Running under: macOS Catalina 10.15.7
##
## Matrix products: default
## BLAS:   /Library/Frameworks/R.framework/Versions/3.6/Resources/lib/libRblas.0.dylib
## LAPACK: /Library/Frameworks/R.framework/Versions/3.6/Resources/lib/libRlapack.dylib
##
## locale:
## [1] en_US.UTF-8/en_US.UTF-8/en_US.UTF-8/C/en_US.UTF-8/en_US.UTF-8
##
## attached base packages:
## [1] stats     graphics  grDevices utils     datasets  methods   base
##
## other attached packages:
## [1] knitr_1.29         gplots_3.1.0       RColorBrewer_1.1-2 edgeR_3.28.1
## [5] limma_3.42.2
##
## loaded via a namespace (and not attached):
##  [1] Rcpp_1.0.5         locfit_1.5-9.4     lattice_0.20-41    gtools_3.8.2
##  [5] digest_0.6.25      bitops_1.0-6       grid_3.6.3         formatR_1.7
```

```
##  [9] magrittr_1.5       evaluate_0.14      KernSmooth_2.23-17 rlang_0.4.8
## [13] stringi_1.5.3      rmarkdown_2.3      splines_3.6.3      tools_3.6.3
## [17] stringr_1.4.0      xfun_0.15          yaml_2.2.1         compiler_3.6.3
## [21] caTools_1.18.0     htmltools_0.5.0
```
