## Supplementary material for "Self-organized yolk sac-like organoids allow for scalable generation of multipotent hematopoietic progenitor cells from human induced pluripotent stem cells": Sup analysis2

### Tamaoki et al. - scRNAseq analysis

2021-03-15 22:10:20

UMI and HTO matrices necessary to rerun the presented analyses are available at GEO (accession GSE157140). Since not all analyses are deterministic we save and load objects on multiple occasions.

```
library(Seurat)
library(dplyr)
```

#### Load raw counts

```
# load 10X data ge.data <- Read10X(data.dir = '../filtered_feature_bc_matrix/')

# load in the UMI matrix ge.umis <- ge.data$`Gene Expression`

# write UMI matrix for GEO write.table(ge.umis,'UMI_mtx.txt', sep = '\t')

# load UMI matrix available at GEO, accession number GSE157140
ge.umis <- read.table("UMI_mtx.txt", sep = "\t", check.names = FALSE, header = TRUE)
ge.umis <- as.matrix(ge.umis)
ge.umis <- as(ge.umis, "sparseMatrix")

# load in the HTO matrix ge.htos <- ge.data$`Antibody Capture`

# for HTO matrix for GEO write.table(ge.htos,'HTO_mtx.txt', sep = '\t')

# load HTO matrix available at GEO, accession number GSE157140
ge.htos <- read.table("HTO_mtx.txt", sep = "\t", check.names = FALSE, header = TRUE)
ge.htos <- as.matrix(ge.htos)
ge.htos <- as(ge.htos, "sparseMatrix")

# select cell barcodes detected for both RNA and HTO
joint.bcs <- intersect(colnames(ge.umis), colnames(ge.htos))

# subset RNA and HTO counts by joint cell barcodes
ge.umis <- ge.umis[, joint.bcs]
# ge.htos <- as.matrix(ge.htos[, joint.bcs])
ge.htos <- ge.htos[, joint.bcs]

# update HTO to treatments
rownames(ge.htos) <- c("D06", "D13")

# setup Seurat object
ge.hashtag <- CreateSeuratObject(counts = ge.umis)

# log normalize RNA data
ge.hashtag <- NormalizeData(ge.hashtag)
```

```r
# find and scale variable features
ge.hashtag <- FindVariableFeatures(ge.hashtag, selection.method = "mean.var.plot")
ge.hashtag <- ScaleData(ge.hashtag, features = VariableFeatures(ge.hashtag))
```

##### Demultiplex

```r
# add HTO data as a new assay independent from RNA
ge.hashtag[["HTO"]] <- CreateAssayObject(counts = ge.htos)

# normalize HTO data, centered log-ratio (CLR) transformation
ge.hashtag <- NormalizeData(ge.hashtag, assay = "HTO", normalization.method = "CLR")

# Demultiplex cells based on HTO enrichment
ge.hashtag <- HTODemux(ge.hashtag, assay = "HTO", positive.quantile = 0.99)

# Visualize demultiplexing results Output from running HTODemux() is saved in the
# object metadata. We can visualize how many cells are classified as singlets,
# doublets and negative/ambiguous cells.

# Global classification results
table(ge.hashtag$HTO_classification.global)
```

```
##
##  Doublet Negative  Singlet
##      528      955    11303
```

```r
# Visualize enrichment for selected HTOs with ridge plots Group cells based on
# the max HTO signal
Idents(ge.hashtag) <- "HTO_maxID"
RidgePlot(ge.hashtag, assay = "HTO", features = rownames(ge.hashtag[["HTO"]])[1:2],
    ncol = 2)
```

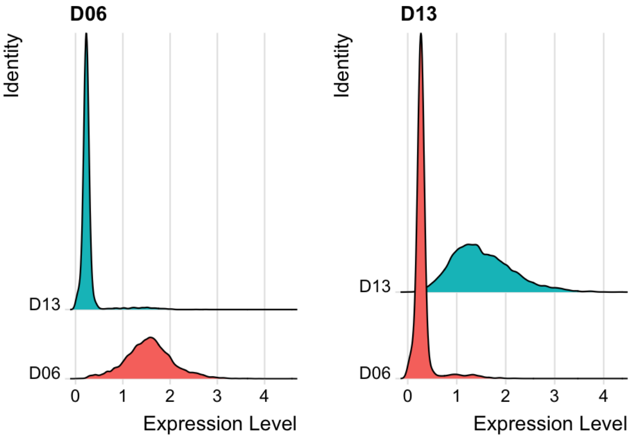


```r
DefaultAssay(object = ge.hashtag) <- "HTO"
# Visualize pairs of HTO signals to confirm mutual exclusivity in singlets
FeatureScatter(ge.hashtag, feature1 = "D06", feature2 = "D13", group.by = "HTO_classification")
```

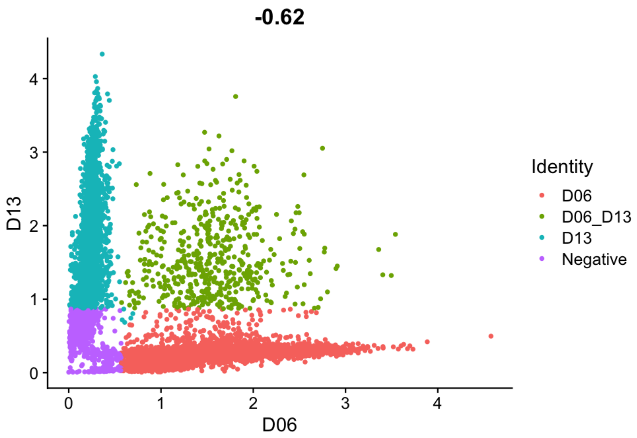


```r
# Compare number of UMIs for singlets, doublets and negative cells
Idents(ge.hashtag) <- "HTO_classification.global"
VlnPlot(ge.hashtag, features = "nCount_RNA", pt.size = 0.1, log = TRUE)
```

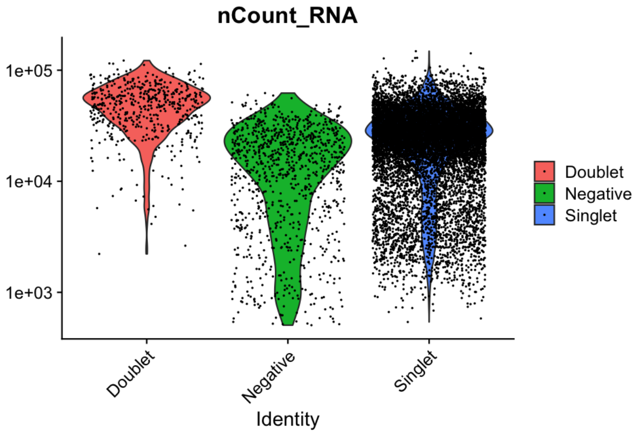


```r
# Number of UMIs/transcripts per cell
mean(ge.hashtag@meta.data$nCount_RNA)
```

```
## [1] 27204.7
```

```r
# Number of genes per cell
mean(ge.hashtag@meta.data$nFeature_RNA)
```

```
## [1] 4693.206
```

```r
# Extract the singlets
ge <- subset(ge.hashtag, idents = "Singlet")

# Number of genes with expression data
length(ge@meta.data$nFeature_RNA)
```

```
## [1] 11303
```

```r
# saveRDS(ge,'../objects/Organoids_Seurat_singlets.rds')
```

## QC

```r
# Dissociation of organoids and fluorescent activated cell sorting (FACS)
# resulted in a fraction of cells with high mitochondrial read counts
```

```r
DefaultAssay(object = ge) <- "RNA"
ge[["percent.mt"]] <- PercentageFeatureSet(ge, pattern = "^MT-", assay = NULL)

VlnPlot(ge, features = c("nFeature_RNA", "nCount_RNA", "percent.mt"), ncol = 3)
```

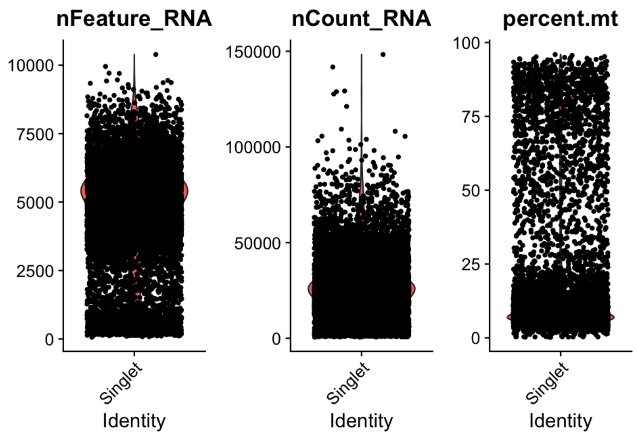


#### Filtering

```r
RidgePlot(ge, features = c("nFeature_RNA", "nCount_RNA", "percent.mt"), ncol = 3)
```

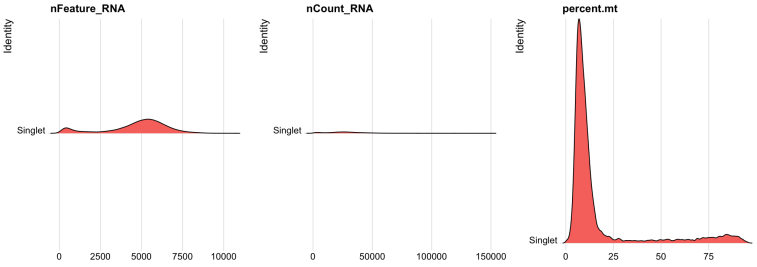


```r
# excluding cells with a mitochondrial read fraction >20%
ge <- subset(ge, subset = nFeature_RNA > 1000 & nFeature_RNA < 8000 & percent.mt <
    20 & nCount_RNA > 2000 & nCount_RNA < 80000)

# Number of genes with expression data
length(ge@meta.data$nFeature_RNA)
```

```
## [1] 9296
```

```r
RidgePlot(ge, features = c("nFeature_RNA", "nCount_RNA", "percent.mt"), ncol = 3)
```

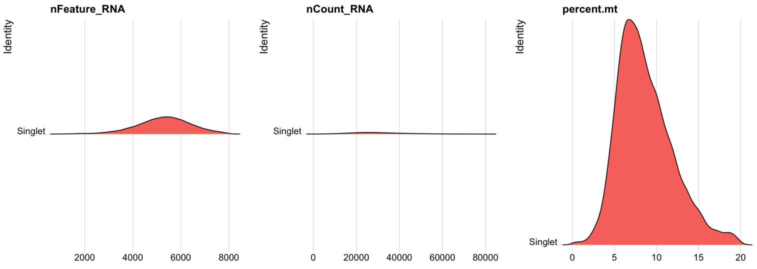


```r
# Number of day 6 cells post filtering
length(which(ge@meta.data$HTO_classification == "D06"))
```

```
## [1] 4895
```

```r
# Number of day 13 cells post filtering
length(which(ge@meta.data$HTO_classification == "D13"))
```

```
## [1] 4401
```

```r
# Number of UMIs/transcripts per cell post filtering
mean(ge@meta.data$nCount_RNA)
```

```
## [1] 30069.91
```

```r
# Number of genes per cell post filtering
mean(ge@meta.data$nFeature_RNA)
```

```
## [1] 5292.572
```

##### Combined clustering - day 6 and day 13 cells

```r
# SCTransform
ge <- SCTransform(ge, vars.to.regress = "percent.mt", verbose = FALSE)
ge <- RunPCA(ge, features = VariableFeatures(object = ge))
# PC selection
ElbowPlot(ge, ndims = 40)
```

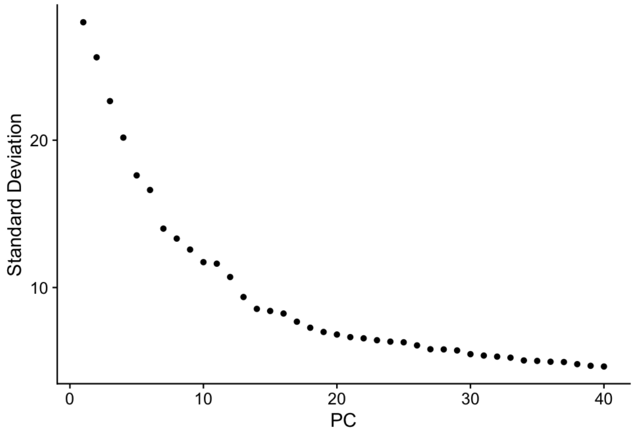


```r
# going with first 30 PCs
ge <- FindNeighbors(ge, dims = 1:30)
ge <- RunUMAP(ge, dims = 1:30)
ge <- FindClusters(ge, resolution = 0.7)
```

```
## Modularity Optimizer version 1.3.0 by Ludo Waltman and Nees Jan van Eck
##
## Number of nodes: 9296
## Number of edges: 311082
##
## Running Louvain algorithm...
## Maximum modularity in 10 random starts: 0.9058
## Number of communities: 16
## Elapsed time: 0 seconds
```

```r
# saveRDS(ge,'../objects/Organoid_Seurat_v3_res0.7.rds')
ge <- readRDS("../objects/Organoids_Seurat_v3_res0.7.rds")
```

```r
# take a look at clustering
DimPlot(ge, reduction = "umap", label = TRUE) + NoLegend()
```

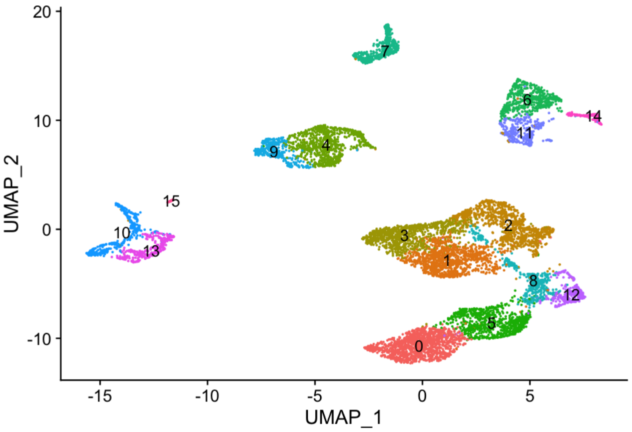


```r
# by batch
DimPlot(ge, reduction = "umap", group.by = "HTO_classification")
```

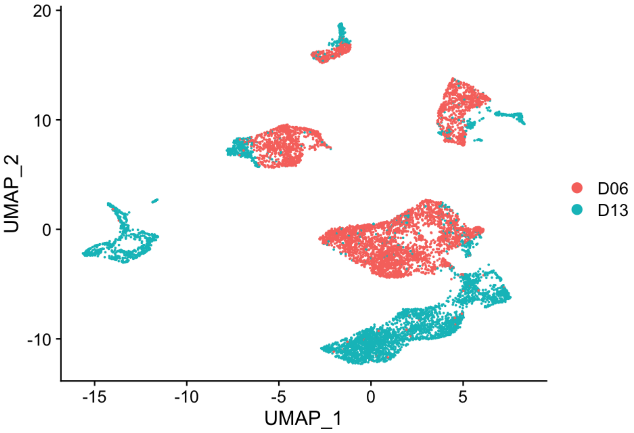


#### Cluster day 6 cells

```r
# subclustering D6
d6 <- rownames(subset(ge@assays$SCT@misc$vst.out$cell_attr, ge@assays$SCT@misc$vst.out$cell_attr$HTO_cla
    "D06"))
ge.d6 <- subset(ge, cells = d6)

ge.d6 <- SCTransform(ge.d6, vars.to.regress = "percent.mt", verbose = FALSE)
ge.d6 <- RunPCA(ge.d6, features = VariableFeatures(object = ge.d6))
ElbowPlot(ge.d6, ndims = 40)
```

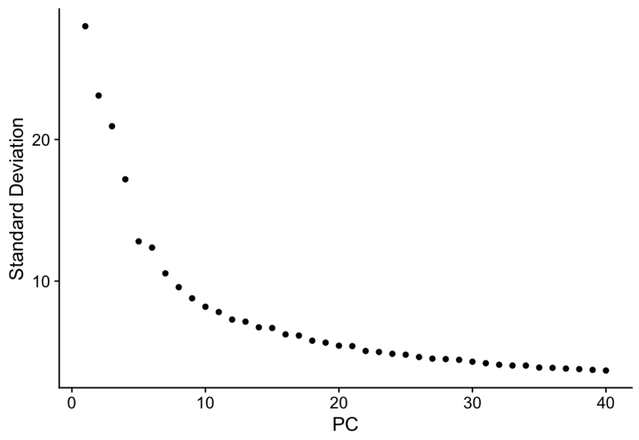


```
# going with first 21 PCs
ge.d6 <- FindNeighbors(ge.d6, dims = 1:21)
ge.d6 <- RunUMAP(ge.d6, dims = 1:21)
ge.d6 <- FindClusters(ge.d6, resolution = 0.3)
```

```
## Modularity Optimizer version 1.3.0 by Ludo Waltman and Nees Jan van Eck
##
## Number of nodes: 4895
## Number of edges: 163306
##
## Running Louvain algorithm...
## Maximum modularity in 10 random starts: 0.9170
## Number of communities: 7
## Elapsed time: 0 seconds
```

```
# saveRDS(ge.d6,'../objects/d6_subclust_0.3.rds')
ge.d6 <- readRDS("../objects/d6_subclust_0.3.rds")
# take a look at clustering
DimPlot(ge.d6, reduction = "umap", label = TRUE) + NoLegend()
```

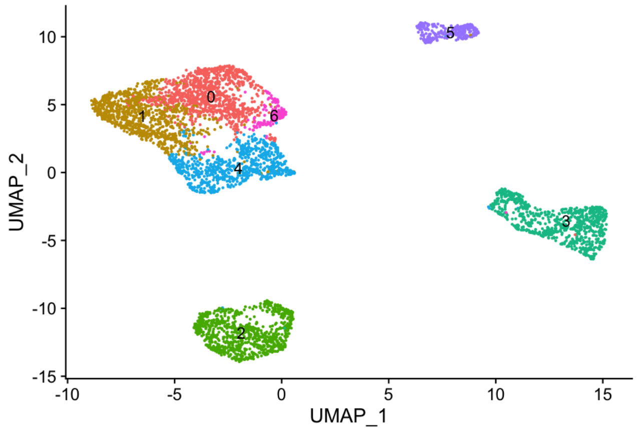


```
new.cluster.ids <- c("C0 MP1", "C1 MP2", "C2 TB", " C3 YE", "C4 MP4", "C5 HE", "C6 MP3")
names(new.cluster.ids) <- levels(ge.d6)
ge.d6 <- RenameIdents(ge.d6, new.cluster.ids)

# colors matching violin plot
colors <- c("#C49A00", "#53B400", "#A58AFF", "#F8766D", "#00B6EB", "#FB61D7", "#00C094")
DimPlot(ge.d6, reduction = "umap", label = TRUE, cols = colors) + NoLegend()
```

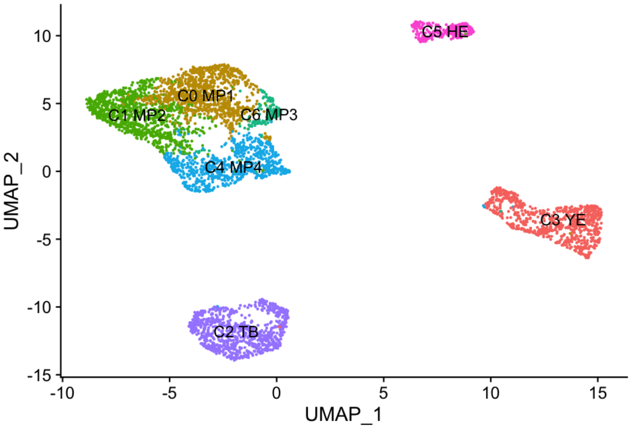


#### Identify day 6 cluster markers

```
ge.d6.markers <- FindAllMarkers(ge.d6, only.pos = TRUE, min.pct = 0.25, logfc.threshold = 0.25)
# saveRDS(ge.d6.markers,'results/d6_markers.RDS') ge.d6.markers <-
# readRDS('results/d6_markers.RDS')

# top 10 markers for each cluster
top10 <- ge.d6.markers %>% group_by(cluster) %>% top_n(n = 10, wt = avg_log2FC)
DoHeatmap(ge.d6, features = top10$gene) + NoLegend()
```

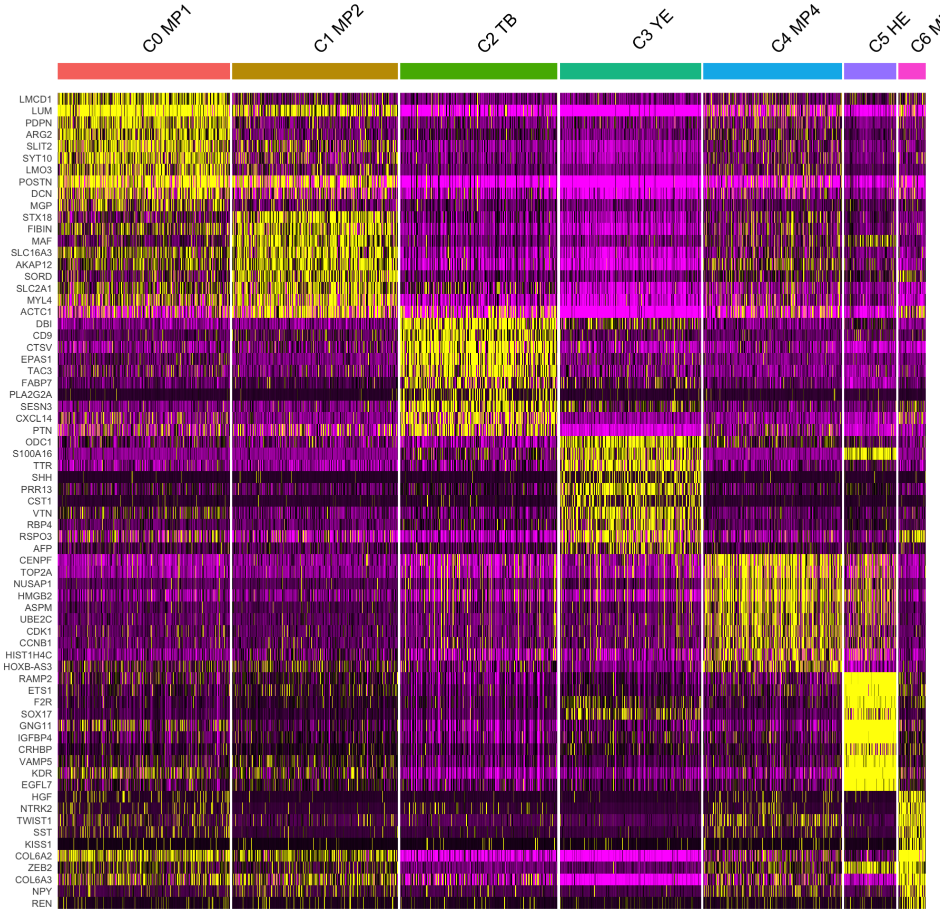

```r
top50 <- ge.d6.markers %>% group_by(cluster) %>% top_n(n = 50, wt = avg_log2FC)
df <- as.data.frame(top50)
# write.csv(df, 'results/d6_top50_marker.csv')
```

#### Cluster day 13 cells

```r
# subclustering D13
d13 <- rownames(subset(ge@assays$SCT@misc$vst.out$cell_attr, ge@assays$SCT@misc$vst.out$cell_attr$HTO_cl
    "D13"))
ge.d13 <- subset(ge, cells = d13)

ge.d13 <- SCTransform(ge.d13, vars.to.regress = "percent.mt", verbose = FALSE)
ge.d13 <- RunPCA(ge.d13, features = VariableFeatures(object = ge.d13))
ElbowPlot(ge.d13, ndims = 40)
```

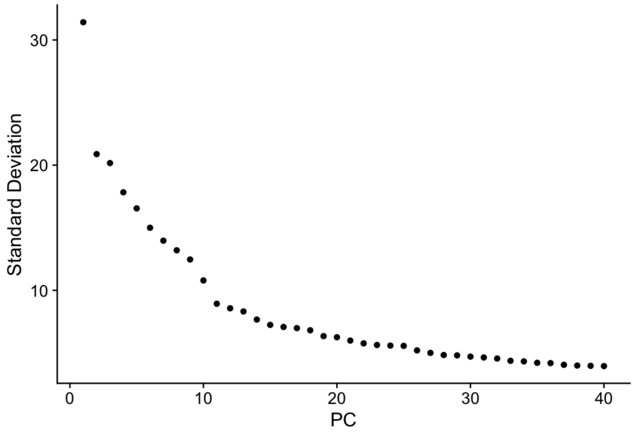


```r
# going with first 25 PCs
ge.d13 <- FindNeighbors(ge.d13, dims = 1:25)
ge.d13 <- RunUMAP(ge.d13, dims = 1:25)
ge.d13 <- FindClusters(ge.d13, resolution = 0.5)
```

```
## Modularity Optimizer version 1.3.0 by Ludo Waltman and Nees Jan van Eck
##
## Number of nodes: 4401
## Number of edges: 139161
##
## Running Louvain algorithm...
## Maximum modularity in 10 random starts: 0.9006
## Number of communities: 12
## Elapsed time: 0 seconds
```

```r
# saveRDS(ge.d13,'../objects/d13_subclust.rds')
ge.d13 <- readRDS("../objects/d13_subclust.rds")

# take a look at clustering
DimPlot(ge.d13, reduction = "umap", label = TRUE) + NoLegend()
```

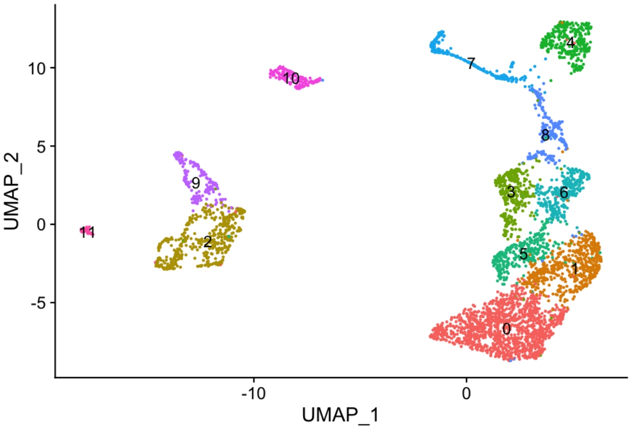


```r
cols <- c("#00C08B", "#00BA38", "#F564E3", "#DE8C00", "#00B4F0", "#00BFC4", "#B79F00",
    "#F8766D", "#7CAE00", "#C77CFF", "#619CFF", "#FF64B0")
new.cluster.ids <- c("C0 SLC", "C1 ST1", "C2 ML", "C3 MS3", "C4 TB", "C5 ST2", "C6 MS2",
    "C7 YE", "C8 MS1", "C9 HPC", "C10 EC", "C11 ER")

names(new.cluster.ids) <- levels(ge.d13)
ge.d13 <- RenameIdents(ge.d13, new.cluster.ids)

DimPlot(ge.d13, reduction = "umap", label = TRUE, cols = cols) + NoLegend()
```

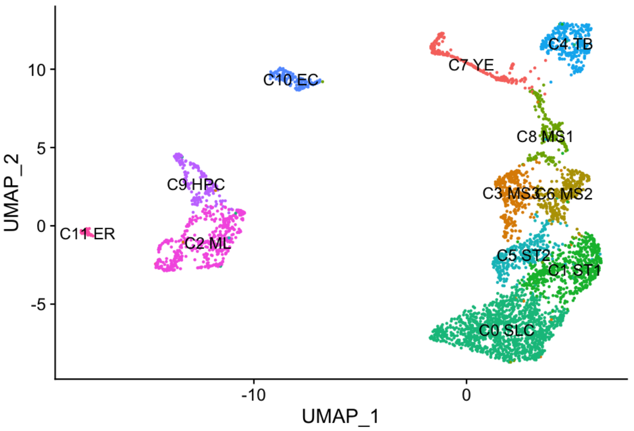


#### Identify day 13 cluster markers

```r
ge.d13.markers <- FindAllMarkers(ge.d13, only.pos = TRUE, min.pct = 0.25, logfc.threshold = 0.25)
# saveRDS(ge.d13.markers,'results/d13_markers.RDS') ge.d13.markers <-
# readRDS('results/d13_markers_newLabel.RDS')

# top 10 markers for each cluster
top10 <- ge.d13.markers %>% group_by(cluster) %>% top_n(n = 10, wt = avg_log2FC)
DoHeatmap(ge.d13, features = top10$gene) + NoLegend()
```

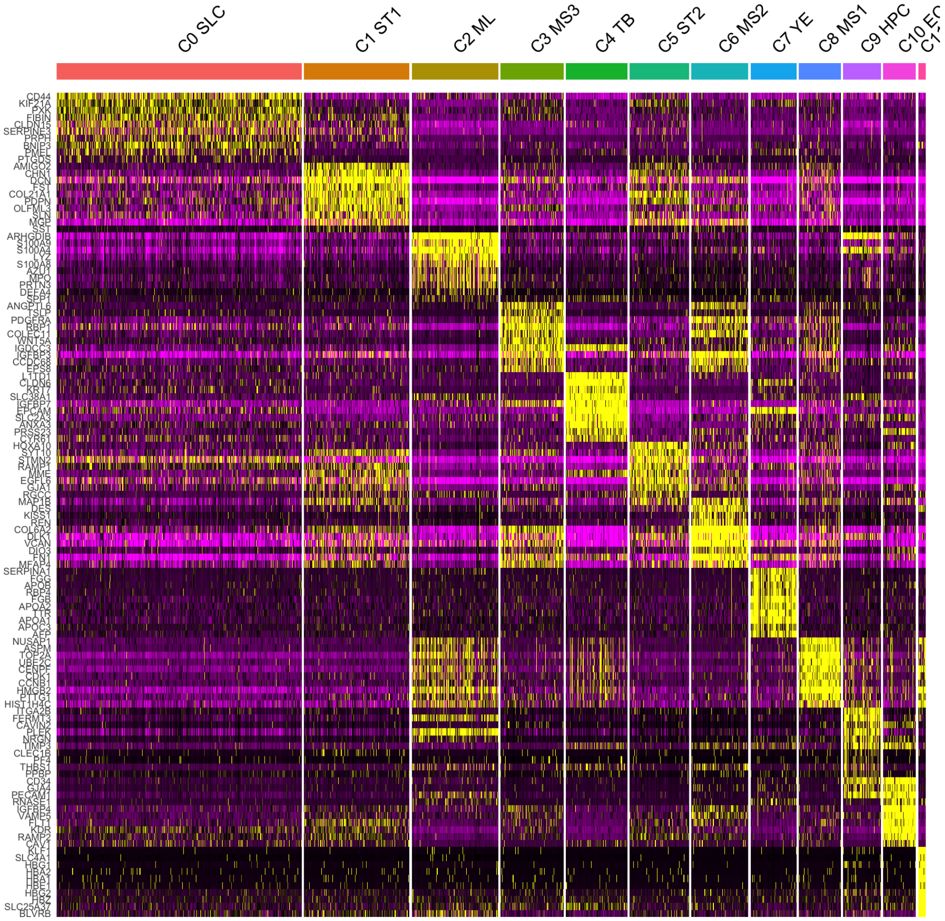


```r
top100 <- ge.d13.markers %>% group_by(cluster) %>% top_n(n = 100, wt = avg_log2FC)
df <- as.data.frame(top100)
# write.csv(df, 'results/d13_top50_markers.csv')
```

##### Map day 6 and day 13 cluster annotations to cells of the combined clustering

```r
ge.d6 <- readRDS("../objects/d6_subclust_0.3.rds")
d6.cluster.ids <- c("6_MP1", "6_MP2", "6_TB", "6_YE", "6_MP4", "6_HE", "6_MP3")
names(d6.cluster.ids) <- levels(ge.d6)
ge.d6 <- RenameIdents(ge.d6, d6.cluster.ids)
# DimPlot(ge.d6, reduction = 'umap', label = TRUE) + NoLegend()

ge.d13 <- readRDS("../objects/d13_subclust.rds")
d13.cluster.ids <- c("d13_SLC", "d13_ST1", "d13_ML", "d13_MS3", "d13_TB", "d13_ST2",
    "d13_MS2", "d13_YE", "d13_MS1", "d13_HPC", "d13_EC", "d13_ER")
names(d13.cluster.ids) <- levels(ge.d13)
ge.d13 <- RenameIdents(ge.d13, d13.cluster.ids)

d6 <- as.data.frame(ge.d6@active.ident)
d6$rows <- rownames(d6)
colnames(d6) <- c("sbcl", "rows")
d13 <- as.data.frame(ge.d13@active.ident)
d13$rows <- rownames(d13)
colnames(d13) <- c("sbcl", "rows")
```

```r
whole <- bind_rows(d6, d13)
rownames(whole) <- whole$rows

ge.w <- readRDS("../objects/Organoids_Seurat_v3_res0.7.rds")
# DimPlot(ge.w, reduction = 'umap', label = TRUE, label.size = 5) + NoLegend()

meta <- ge.w@meta.data
meta$rows <- rownames(meta)
de <- merge(ge.w@meta.data, whole, by = "row.names", all = TRUE)
# colnames(de)

new.meta <- plyr::join(meta, whole, by = "rows")
rownames(new.meta) <- new.meta$rows
new.meta$rows <- NULL

ge.w@meta.data <- new.meta
# saveRDS(ge.w,'../objects/d6_d13_with_subcl_labels.rds')
```

##### Combined clustering with annotations from day 6 and day 13 clusterings

```r
DimPlot(ge.w, reduction = "umap", group.by = "sbcl", label = TRUE) + NoLegend()
```

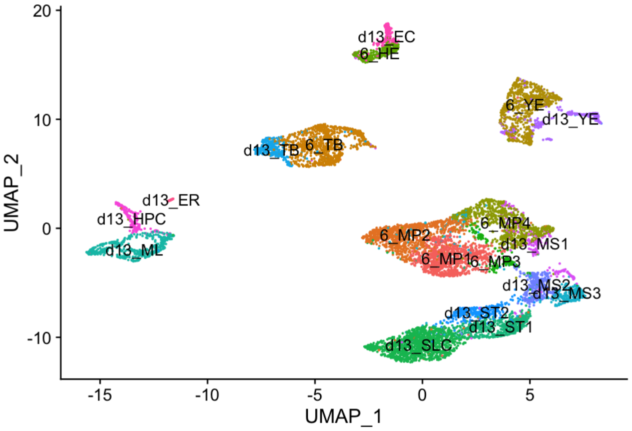


##### Subcluster hematogenic lineage cells from day 13

```r
ge <- readRDS("../objects/d13_subclust.rds")
hem <- subset(ge, idents = c("2", "9", "11"))
# DimPlot(hem, reduction = 'umap', label = TRUE) + NoLegend()

hem <- SCTransform(hem, vars.to.regress = "percent.mt", verbose = FALSE)
hem <- RunPCA(hem, features = VariableFeatures(object = hem))
ElbowPlot(hem, ndims = 50)
```

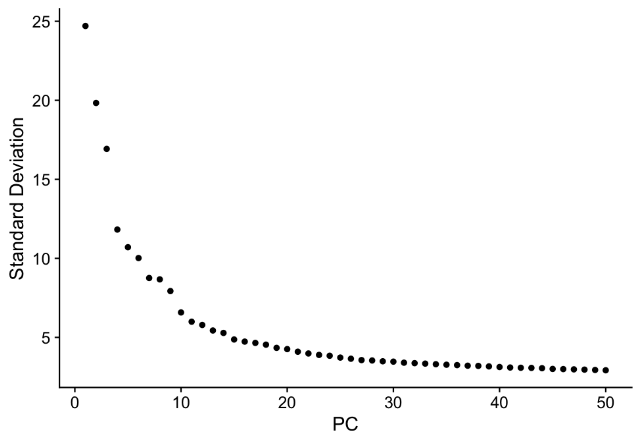


```
# going with first 30 PCs
hem <- FindNeighbors(hem, dims = 1:40)
hem <- RunUMAP(hem, dims = 1:40)
hem <- FindClusters(hem, resolution = 0.5)
```

```
## Modularity Optimizer version 1.3.0 by Ludo Waltman and Nees Jan van Eck
##
## Number of nodes: 689
## Number of edges: 19849
##
## Running Louvain algorithm...
## Maximum modularity in 10 random starts: 0.8347
## Number of communities: 6
## Elapsed time: 0 seconds
```

```
# saveRDS(hem,'../objects/d13_hem_subclust.rds')
hem <- readRDS("../objects/d13_hem_subclust.rds")
DimPlot(hem, reduction = "umap", label = TRUE) + NoLegend()
```

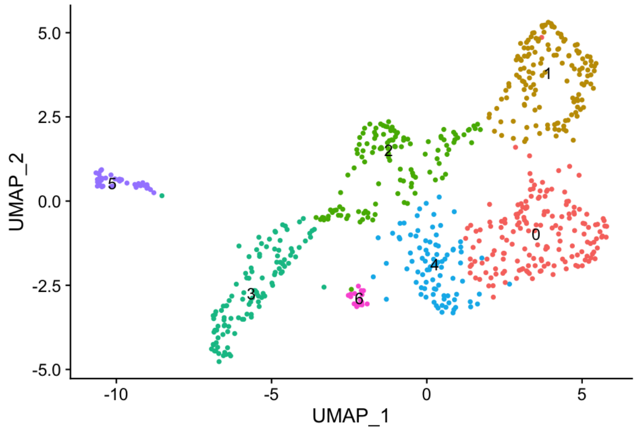


```
hem.markers <- FindAllMarkers(hem, only.pos = TRUE, min.pct = 0.25, logfc.threshold = 0.25)
# saveRDS(hem.markers,'results/d13_hem_markers.RDS') hem.markers <-
# readRDS('results/d13_hem_markers.RDS')

# top 10 markers for each cluster
top10 <- hem.markers %>% group_by(cluster) %>% top_n(n = 10, wt = avg_log2FC)
DoHeatmap(hem, features = top10$gene) + NoLegend()
```

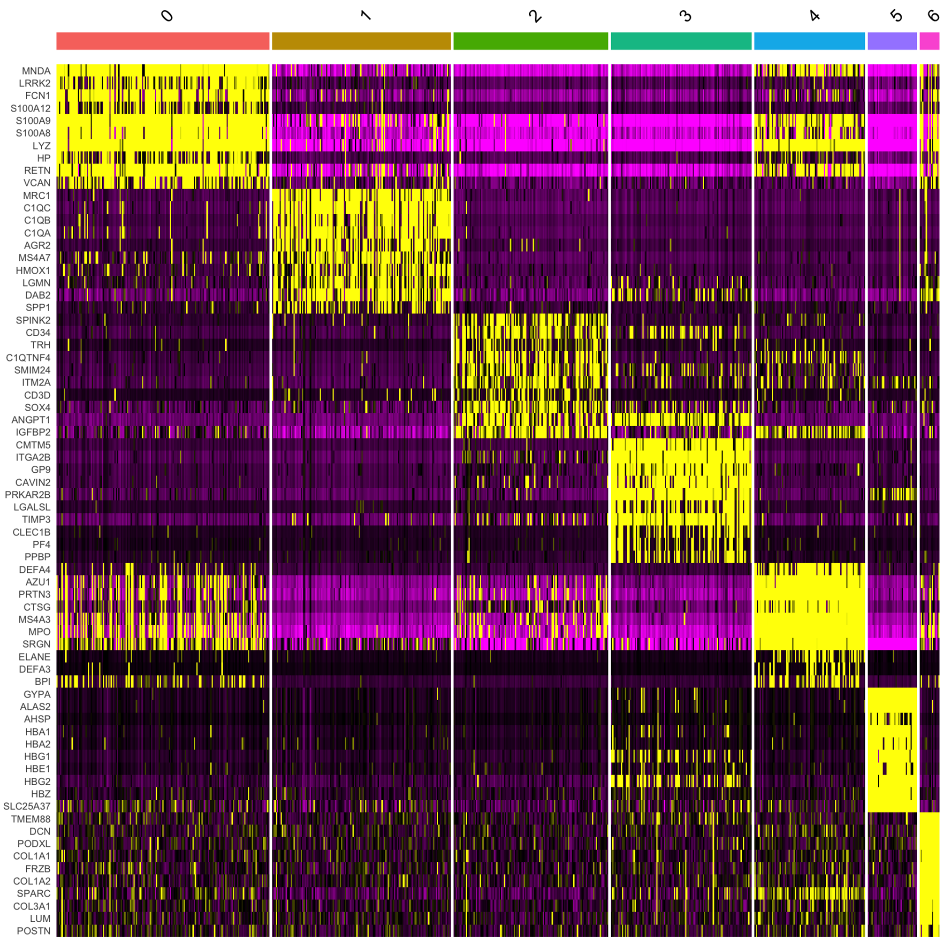


```r
top50 <- hem.markers %>% group_by(cluster) %>% top_n(n = 50, wt = avg_log2FC)
df <- as.data.frame(top50)
# write.csv(df, 'results/hem_top50_markers.txt')
```

```r
sessionInfo()
```

```
## R version 3.6.3 (2020-02-29)
## Platform: x86_64-apple-darwin15.6.0 (64-bit)
## Running under: macOS Catalina 10.15.7
##
## Matrix products: default
## BLAS:   /Library/Frameworks/R.framework/Versions/3.6/Resources/lib/libRblas.0.dylib
## LAPACK: /Library/Frameworks/R.framework/Versions/3.6/Resources/lib/libRlapack.dylib
##
## locale:
## [1] en_US.UTF-8/en_US.UTF-8/en_US.UTF-8/C/en_US.UTF-8/en_US.UTF-8
##
## attached base packages:
## [1] stats     graphics  grDevices utils     datasets  methods   base
##
## other attached packages:
## [1] dplyr_1.0.2       Seurat_3.9.9.9002 knitr_1.29
##
## loaded via a namespace (and not attached):
##   [1] nlme_3.1-148          matrixStats_0.57.0    RcppAnnoy_0.0.16
```

```
##   [4] RColorBrewer_1.1-2    httr_1.4.2            sctransform_0.3.1
##   [7] tools_3.6.3           R6_2.4.1              irlba_2.3.3
##  [10] rpart_4.1-15          KernSmooth_2.23-17    uwot_0.1.8.9001
##  [13] mgcv_1.8-31           lazyeval_0.2.2        colorspace_1.4-1
##  [16] withr_2.3.0           tidyselect_1.1.0      gridExtra_2.3
##  [19] compiler_3.6.3        formatR_1.7           plotly_4.9.2.1
##  [22] labeling_0.3          scales_1.1.1          lmtest_0.9-38
##  [25] spatstat.data_1.4-3   ggridges_0.5.2        pbapply_1.4-3
##  [28] spatstat_1.64-1       goftest_1.2-2         stringr_1.4.0
##  [31] digest_0.6.25         spatstat.utils_1.17-0 rmarkdown_2.3
##  [34] pkgconfig_2.0.3       htmltools_0.5.0       limma_3.42.2
##  [37] fastmap_1.0.1         htmlwidgets_1.5.2     rlang_0.4.8
##  [40] shiny_1.5.0           farver_2.0.3          generics_0.0.2
##  [43] zoo_1.8-8             jsonlite_1.7.1        ica_1.0-2
##  [46] magrittr_1.5          patchwork_1.0.1       Matrix_1.2-18
##  [49] Rcpp_1.0.5            munsell_0.5.0         abind_1.4-5
##  [52] reticulate_1.16       lifecycle_0.2.0       stringi_1.5.3
##  [55] yaml_2.2.1            MASS_7.3-51.6         Rtsne_0.15
##  [58] plyr_1.8.6            grid_3.6.3            parallel_3.6.3
##  [61] listenv_0.8.0         promises_1.1.1        ggrepel_0.8.2
##  [64] crayon_1.3.4          deldir_0.1-29         miniUI_0.1.1.1
##  [67] lattice_0.20-41       cowplot_1.1.0         splines_3.6.3
##  [70] tensor_1.5            magick_2.4.0          pillar_1.4.6
##  [73] igraph_1.2.6          future.apply_1.6.0    reshape2_1.4.4
##  [76] codetools_0.2-16      leiden_0.3.3          glue_1.4.2
##  [79] evaluate_0.14         data.table_1.12.8     vctrs_0.3.4
##  [82] png_0.1-7             httpuv_1.5.4          gtable_0.3.0
##  [85] RANN_2.6.1            purrr_0.3.4           polyclip_1.10-0
##  [88] tidyr_1.1.2           future_1.19.1         ggplot2_3.3.2
##  [91] xfun_0.15             rsvd_1.0.3            mime_0.9
##  [94] xtable_1.8-4          later_1.1.0.1         survival_3.2-3
##  [97] viridisLite_0.3.0     tibble_3.0.4          cluster_2.1.0
## [100] globals_0.13.1        fitdistrplus_1.1-1    ellipsis_0.3.1
## [103] ROCR_1.0-11
```
